# Hippocampal memory resilience amidst state-dependent homeostatic regulation

**DOI:** 10.64898/2026.11.23.767553

**Authors:** Refaela Atsmon, Karni Lev Bar-Or, Lee Susman, Liudmila Sosulina, Hiroshi Kaneko, Amit Zilberstein, Lior De Marcas, David Mandil, Antonella Ruggiero, Bhargavi Murthy, Leore R. Heim, Ortal Shinikamin, Ran Darshan, Alessio Attardo, Stefan Remy, Inna Slutsky

## Abstract

Whether and how the hippocampus, a central hub of learning and memory, maintains stable network activity and keeps stored memories accessible despite continual perturbations remains poorly understood. We chemogenetically activated hippocampal interneurons for days while chronically recording single-unit activity in behaving adult mice. The perturbation acutely suppressed firing rates and reorganized higher-order network features, including functional correlations, dimensionality, and dynamical stability. Despite continued inhibition, these properties recovered with distinct time courses, revealing multiscale homeostatic regulation. Correlations and dynamical stability converged to vigilance-state-dependent set points, highest during NREM sleep. Firing rate recovery was accompanied by sleep-specific changes in within-epoch dynamics: attenuated decreases during NREM sleep and a sustained reversal from decreases to increases during REM sleep, whereas increases during wake were not enhanced. CA1 pyramidal neurons increased intrinsic excitability without evidence of synaptic scaling. Retrieval of CA1-dependent memories was transiently impaired and recovered alongside network renormalization, with strength and specificity preserved. These findings reveal state-dependent constraints on homeostatic regulation, with multivariate network stabilization coinciding with memory resilience.

## Introduction

The hippocampus is a pivotal hub for experience-dependent neuronal plasticity^1^, essential for encoding and retrieval of memories^2–6^. It undergoes continual structural and functional remodeling and is subject to ongoing external and internal perturbations, yet the memories it stores must remain accessible throughout. A central question, therefore, is how hippocampal circuits maintain stable network activity while preserving access to stored information. Stability and flexibility are not independent demands: Hebbian plasticity is fundamental for memory formation and storage but poses challenges to network stability^7,8^, a plasticity–stability dilemma, recognized in both artificial and biological neural networks^9–12^. Disruption of this balance has been implicated in memory disorders including Alzheimer’s disease^13^. Understanding how hippocampal networks stabilize their activity without compromising stored memories is thus important for explaining both memory persistence and forgetting^2,3,14–16^.

Despite extensive research on hippocampal plasticity, the *in vivo* mechanisms that confer resilience - the capacity of neural circuits to recover from perturbation - remain poorly understood. Computational studies suggest that resilience arises from multiple, degenerate stabilizing processes^17–19^ operating across diverse timescales to maintain different aspects of neuronal activity^7,20^. Homeostatic plasticity offers a principled solution: slow, negative-feedback mechanisms that regulate neuronal and network activity around a set point in the face of sustained perturbations^9,21–23^. Firing rate (FR) homeostasis has been extensively studied in the primary visual cortex (V1) following sensory perturbations^24–31^, implicating both synaptic and intrinsic homeostatic mechanisms^24,25,30–34^. However, network function also depends on population-level features, including pairwise correlations, dimensionality, criticality, and dynamical stability (the relaxation timescale of population fluctuations), that jointly shape information flow and robustness and vary with vigilance state^35–37^. Whether these higher-order metrics are disrupted by prolonged inhibitory drive and, whether they recover toward state-dependent set points, remains unknown. More broadly, it is unclear how hippocampal population dynamics recover across vigilance states, and whether such recovery preserves access to previously stored memories.

Hebbian and homeostatic plasticity are thought to act in concert to support stable circuit function^11,38–40^. While Hebbian-like synaptic plasticity has been extensively studied in the adult hippocampus^1,41,42^, homeostatic mechanisms have been characterized mainly during development, often in hippocampal primary or organotypic slice cultures^43–53^. Studies in the V1 cortex show that the extent and site of expression of homeostatic regulation depend on circuit maturation^54–57^. It is therefore important to determine which homeostatic mechanisms persist in the mature hippocampus^58^ and whether vigilance state constrains them, as has been observed in developing V1^27,28^. In particular, whether sleep promotes^59–63^ or suppresses^27^ homeostatic regulation in response to inactivity in the hippocampus remains unknown.

In this study, we address three questions: (1) which features of hippocampal network activity are regulated in response to persistent perturbations; (2) how stabilizing mechanisms operate across sleep–wake states; and (3) whether these homeostatic mechanisms^64–67^ can restore hippocampal activity while preserving retrieval of previously stored memories. To this end, we combined long-term extracellular electrophysiological recordings in the hippocampal CA1 region of behaving mice with computational analyses of neuronal dynamics, brain states, and behavioral memory assays. To probe homeostatic regulation, we used continuous chemogenetic activation of hippocampal interneurons (INs), a manipulation that strongly suppresses CA1 pyramidal spiking^68^. We find that CA1 FR recovery is preferentially expressed during sleep, most prominently during REM sleep. Renormalization of excitatory network dynamics was accompanied by restored access to previously stored memories without loss of context specificity. Together, these findings reveal principles governing hippocampal stability *in vivo* and show that renormalization of network dynamics coincides with preserved memory accessibility during persistent perturbation.

## Results

### Interneuron hM3Dq activation enhances inhibitory conductance and suppresses CA1 network firing

Some physiological variables maintain baseline stability; but the key determinant of stable dynamics is their ability to recover following perturbation. Persistent perturbation is readily achieved in sensory cortices by sensory deprivation, but this approach is less straightforward for the hippocampus. To determine whether CA1 activity is homeostatically regulated at the population level, we established a perturbation designed to suppress FRs of CA1 pyramidal neurons in behaving mice. We employed a chemogenetic approach using Designer Receptors Exclusively Activated by Designer Drugs (DREADDs) to activate Gq signaling in local INs (Fig. 1a). Specifically, we expressed the Gq-coupled DREADD receptor hM3Dq under the hDlx promoter to target INs^68^. In the experimental (Dlx-Gq) group, AAV-hDlx-hM3Dq-dTomato was injected bilaterally into the dorsal CA1 of adult male mice three weeks before tetrode implantation. Histological evaluation demonstrated widespread viral expression across all CA1 layers and IN subtypes, with dTomato signals colocalizing with GAD-67–positive INs (Fig. 1b, Extended Data Fig. 1a). Control (Dlx-Cnt) mice underwent the same procedure but received a viral vector containing only the dTomato fluorescent reporter, without the DREADD-coding sequence. This design accounted for potential confounding effects of surgery, viral vector expression, and any off-target effects of systemic CNO delivery.

**Fig. 1.**
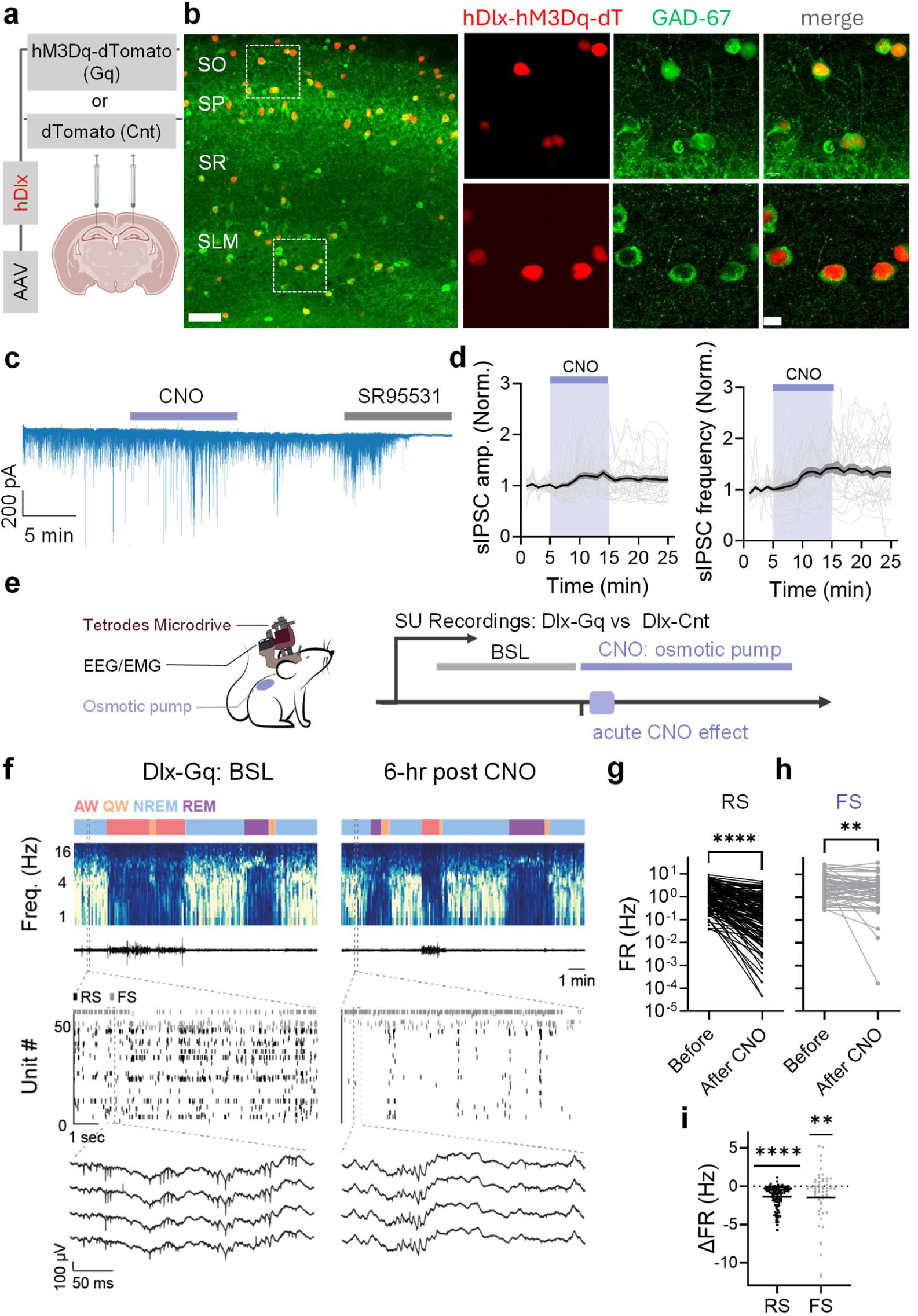
Short-term chemogenetic activation of hM3Dq in CA1 interneurons increases inhibitory synaptic currents in CA1 pyramidal neurons and suppresses their firing. **a,** Schematic of viral constructs used in each experimental group: AAV-hDlx-hM3D(Gq)-2A-NLS-dTomato (Dlx-Gq) or AAV-hDlx-HBB-chI-HA_NLS-dTomato (Dlx-Cnt). A volume of 0.5 µl was injected into the dorsal CA1 pyramidal layer of each hemisphere. **b,** Representative histological section of CA1 showing AAV-hDlx-hM3D(Gq)-2A-NLS-dTomato expression (red) in interneurons across stratum oriens (SO), stratum pyramidale (SP), stratum radiatum (SR), and stratum lacunosum-moleculare (SLM). Immunostaining for GAD-67 (green) confirmed interneuron identity; merged images demonstrate colocalization of dTomato and GAD-67. Scale bars: 50 μm (left) and 10 μm (right). **c,** Representative traces of sIPSCs recorded before and during application of 10 μM CNO. sIPSCs were abolished by subsequent application of the GABA_A_ receptor antagonist SR95531 (20 μM), confirming their inhibitory nature. **d,** Time course plots showing the amplitude (left) and frequency (right) of sIPSCs. Data were binned in 1-min intervals and normalized to the mean of the initial 5-min baseline period. **e,** Schematic of microdrive placement for single-unit recordings, EEG/EMG electrodes, and a subcutaneous osmotic pump in a mouse (left). Experimental timeline (right): short-term effect of CNO was quantified before (BSL) and 6-hr after CNO-loaded osmotic pump implantation. **f,** Representative recordings of CA1 activity before and 6 hr after CNO delivery via osmotic pump. From top to bottom: hypnogram, spectrogram, EMG, raster plot of regular-spiking (RS) units (black) and fast-spiking (FS) units (gray), and raw traces from a single tetrode. Dashed lines indicate expanded views of the corresponding sections above. **g,** Firing rates of RS units (*n* = 132 units, 5 mice, Dlx-Gq group) before and 6 hr after CNO delivery, showing a significant reduction (\*\*\*\**P* < 0.0001). **h,** Firing rates of FS units (*n* = 60 units, 5 mice, Dlx-Gq group) before and 6 hr after CNO delivery, showing heterogeneous responses with a significant overall reduction (\*\**P* = 0.0054). i, Change in firing rate (ΔFR) of RS and FS units 6 hr after CNO delivery. Both populations exhibited a negative fold change (RS: \*\*\*\**P* < 0.0001; FS: \*\**P* = 0.0073). Wilcoxon matched-pairs signed-rank test (g,h); one-sample Wilcoxon test (i).

Patch-clamp recordings in acute hippocampal slices, in the presence of AMPA and NMDA receptor blockers, confirmed that bath CNO application increased inhibitory conductance, measured as spontaneous inhibitory postsynaptic currents (sIPSCs), in CA1 pyramidal neurons from Dlx-Gq mice (Fig. 1c–d, Extended Data Fig. 1b–c). Prolonged CNO incubation for five days likewise enhanced inhibitory conductance *ex vivo* (Extended Data Fig. 1d), indicating sustained hM3Dq receptor efficacy across the perturbation period.

To ensure stable and sustained activation of hM3Dq signaling^69^ in vivo, we implanted subcutaneous osmotic pumps delivering CNO continuously (0.5 mg/kg/hr). Recordings spanned across a 2-day baseline period prior to CNO exposure (“BSL stage”), followed by 5 days of continuous CNO delivery (“CNO stage”) (Fig. 1e). To monitor FR across the sleep-wake cycle, we conducted simultaneous single-unit recordings, local field potentials (LFPs) in the CA1, electroencephalography (EEG), electromyography (EMG), and video recordings in both groups during natural behavior in their home cages (Fig. 1f). We categorized brain activity into four distinct states: active wakefulness (AW), quiet wakefulness (QW), non-rapid eye movement sleep (NREMS), and rapid eye movement sleep (REMS) (Extended Data Fig. 2a-b). Recordings resumed 30–60 min after pump implantation. Although implantation transiently disrupted behavior, Dlx-Cnt mice recovered within 6 hr, as indicated by restoration of sleep– wake architecture to baseline (Extended Data Fig. 2c,d) and unaltered performance in open-field and Delta maze assays (Extended Data Fig. 3). Consequently, the first 6 hr post-surgery were excluded from FR analyses in both groups.

Single units were classified into regular-spiking (RS) and fast-spiking (FS) based on waveform features and firing patterns (Extended Data Fig. 4). To assess the short-term effect of the perturbation on CA1 neurons *in vivo*, we analyzed FR of RS and FS units before and 6 hr after CNO administration. RS units showed a robust ∼72% reduction in FR (from 1.87 ± 0.16 to 0.52 ± 0.08 Hz; 132 units, 5 mice; Fig. 1g,i). No change in FR of RS units was observed in Dlx-Cnt group (Extended Data Fig. 1e), suggesting that pump implantation and CNO administration do not alter hippocampal neuronal firing in the absence of hM3Dq. Notably, FS units displayed a heterogeneous response to CNO, but on average exhibited paradoxical suppression of similar magnitude, from 5.06 ± 0.78 to 3.65 ± 0.69 Hz (60 units, 5 mice, Fig. 1h,i). This ‘paradoxical’ suppression of INs is consistent with inhibition-stabilized network (ISN) dynamics (see fitted models in Extended Data Fig. 5). In such a dynamical regime, increased inhibition suppresses principal-cell firing and, via reduced excitatory feedback, yields a net decrease in IN activity. These results are in line with prior *in vivo* hippocampal findings^70,71^ and early theoretical predictions of ISN models^72,73^, although they may also be explained by alternative mechanisms, such as the engagement of distinct populations of inhibitory neurons^74,75^ or overrepresentation of chain motifs^76^.

### Regulation of CA1 network FR in response to persistent IN perturbation

To test whether population-level mean activity in CA1 is homeostatically regulated, we evaluated its capacity to recover under a persistent perturbation in behaving mice (Fig. 2a). We assessed the effect of prolonged increases in inhibitory drive on RS-unit activity (24.7 ± 3.8 units per mouse per 24-h clustering block; 5 mice over 7 recording days; resulting in 119.7 ± 8.4 units per recording day across all mice). During baseline recordings, RS units exhibited a stable mean FR (MFR). Following CNO delivery, RS units in Dlx-Gq mice showed a robust acute suppression of MFR that gradually recovered to baseline levels within 2–3 days of continuous CNO exposure (Fig. 2b,d; Extended Data Fig. 6a), with quantitatively similar recovery trajectories across individual mice (Extended Data Fig. 6b). CNO had no detectable effect on RS-unit MFR in Dlx-Cnt mice throughout the experiment (Fig. 2c, Extended Data Fig. 6c). In 4 out of 5 mice, we recorded for additional two days after pump removal (22.5 ± 3.6 RS units per mouse per day; resulting in 90 ± 5 units per washout recording day across 4 mice). We found that the Dlx-Gq group exhibited an overshoot of RS-unit MFR above baseline (Fig. 2f, Extended Data Fig. 6e). This rebound is consistent with engagement of negative-feedback homeostatic mechanisms in response to prior suppression and is not readily explained by waning chemogenetic efficacy (Extended Data Fig. 1d). Similar rebounds have been reported in hippocampal cultures^50^.

**Fig. 2.**
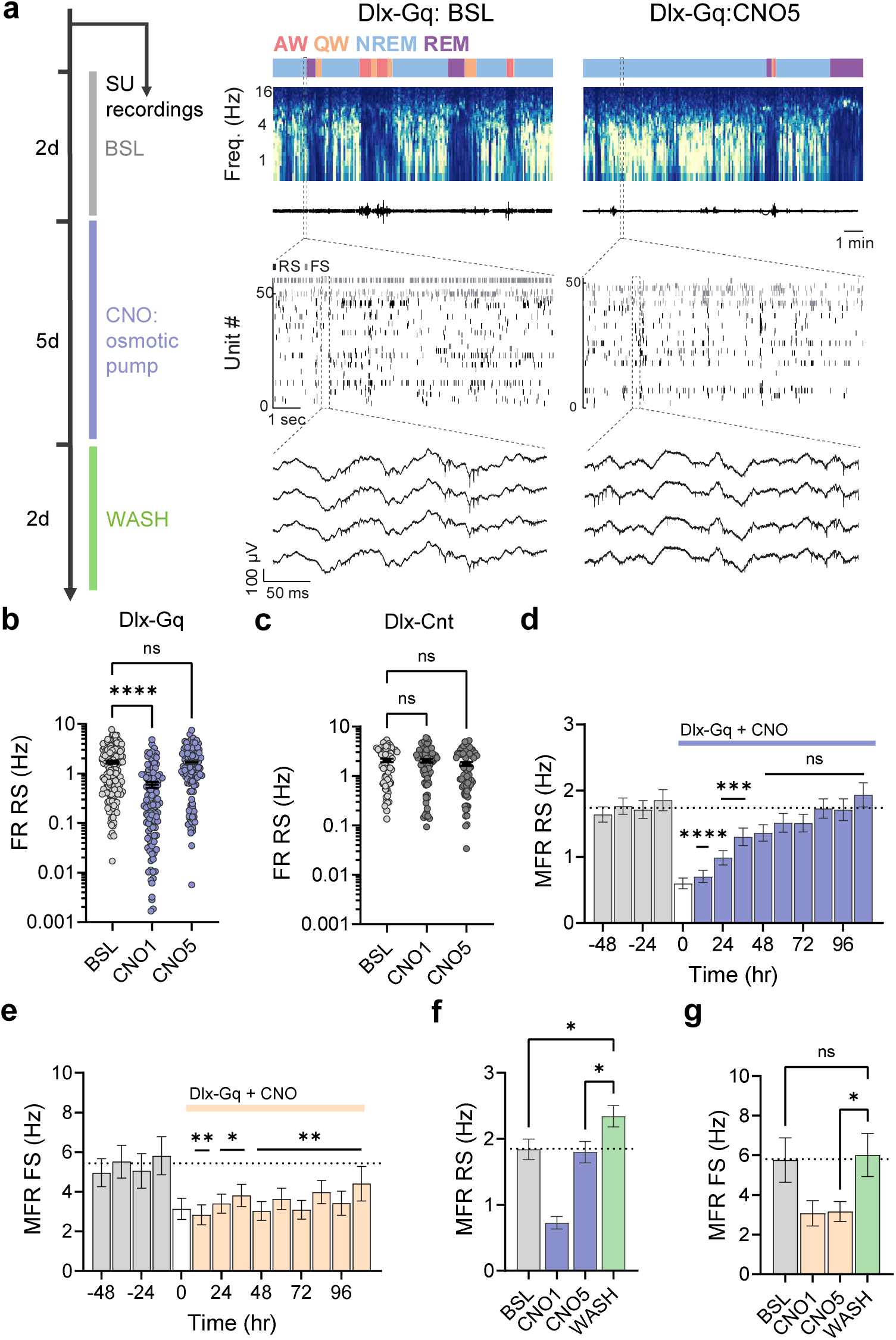
MFR of CA1 RS units is homeostatically restored following persistent chemogenetic perturbation of INs, whereas FS units fail to recover. **a,** Left: Experimental timeline: after 2 days of baseline (BSL) recordings, a CNO-loaded osmotic pump was implanted for 5 days; recordings continued for two additional days after pump removal (WASH). Right: Representative recordings of CA1 activity at baseline (BSL) and on day 5 following CNO delivery via osmotic pump (CNO5). From top to bottom: hypnogram, spectrogram, EMG, raster plot of regular-spiking (RS; black) and fast-spiking (FS; gray) units, and raw traces from a single tetrode. Dashed lines indicate expanded views of the corresponding sections above. **b**, FR of RS units in the Dlx-Gq group during 12-hr windows across baseline (BSL), day 1 of chronic CNO administration (CNO1), and day 5 of chronic CNO administration (CNO5). FR was reduced on CNO1 (\*\*\*\**P* < 0.0001) and recovered by CNO5 (*P* > 0.99). For pairwise statistical contrasts, see Table S1a. **c**, FR of RS units in the Dlx-Cnt group across BSL, CNO1, and CNO5. CNO administration had no effect on FR (CNO1: *P* > 0.99; CNO5: *P* = 0.34). pairwise contrasts in Table S1b. **d**, MFR of RS (24.7 ± 3.8 units per mouse per day; 119.7 ± 8.4 units per recording day across 5 mice; 838 units in total for 7 recording days) in the Dlx-Gq group during BSL (gray) and CNO exposure (purple). The first 6 hr post-CNO (white) were excluded from analysis. RS MFR decreased during 0–48 hr post-CNO (\*\*\*\**P* ≤ 0.0001) but recovered by 48–108 hr (*P* = 0.46). For pairwise statistical contrasts, see Table S1c. **e**, The same as (d) but for FS units (12.1 ± 1.3 units per mouse per day; 58.3 ± 3.3 units per recording day across 5 mice, 408 units in total for 7 recording days). FS MFR decreased during 0-48 hr (\**P* ≤ 0.02) and remained suppressed throughout 48–108 hr (\*\**P* = 0.007), with no recovery until pump removal. For pairwise statistical contrasts, see Table S1d. **f**, RS MFR increased after pump removal (WASH, 24 hr post-removal, *n = 4 mice, 22.5 ± 3.6 units per mouse per day*; 90 ± 5 units per day for all mice), relative to both BSL (BSL vs. WASH, \**P* = 0.043) and CNO5 (CNO5 vs. WASH, \**P* = 0.036) in Dlx-Gq group. For pairwise statistical contrasts, see Table S1e. **g**, FS MFR returned to baseline levels following pump removal (BSL vs. WASH, *P* = 0.97; CNO5 vs. WASH, \**P* = 0.046, *n* = 4 mice, 9.8 ± 0.3 units per mouse per day, 39 ± 2 per day for all mice) in Dlx-Gq group. For pairwise statistical contrasts, see Table S1f. Linear mixed-effects regression (LMER; b-g). Data are presented as mean ± s.e.m.

In contrast to the recovery of RS units, paradoxical inhibition of FS units (12.1 ± 1.3 units per mouse per day, 5 mice over 7 recording days; resulting in 58.3 ± 3.3 units per recording day across all mice) persisted throughout the perturbation (Fig. 2e). The absence of renormalization in FS-unit activity confirms sustained effectiveness of the perturbation during CNO administration. CNO did not alter FS-unit MFR in Dlx-Cnt mice (Extended Data Fig. 6d). During the washout period after pump removal, the Dlx-Gq group exhibited a return of FS-unit MFR to baseline (9.8 ± 0.3 units per mouse per day; resulting in 39 ± 2 units per recording day across 4 mice; Fig. 2g).

Together, these results indicate that RS-unit MFR is regulated around a network-level set point in CA1, underscoring the capacity of excitatory CA1 circuits to compensate for a persistent IN perturbation in behaving mice.

### Regulation of CA1 FR state-space dynamics in response to persistent IN perturbation

Computational capacity, and likely also homeostatic mechanisms, rely not only on mean population activity but also on the structure of fluctuations across time and neurons. We therefore examined CA1 FR dynamics at a finer scale, focusing on activity patterns unfolding over seconds to minutes, and at the properties of co-fluctuations of neurons in the population.

We first looked at a network property associated with Hebbian plasticity, the functional connectivity among neurons, quantified by pairwise Pearson correlation coefficients (CCs) between RS units (Fig. 3a). Mean CCs were stable during baseline, but increased sharply following IN activation, consistent with enhanced synchrony. Over the course of 3–4 days, similar to the recovery of mean FR, correlations gradually renormalized to baseline (Fig. 3b).

**Fig. 3.**
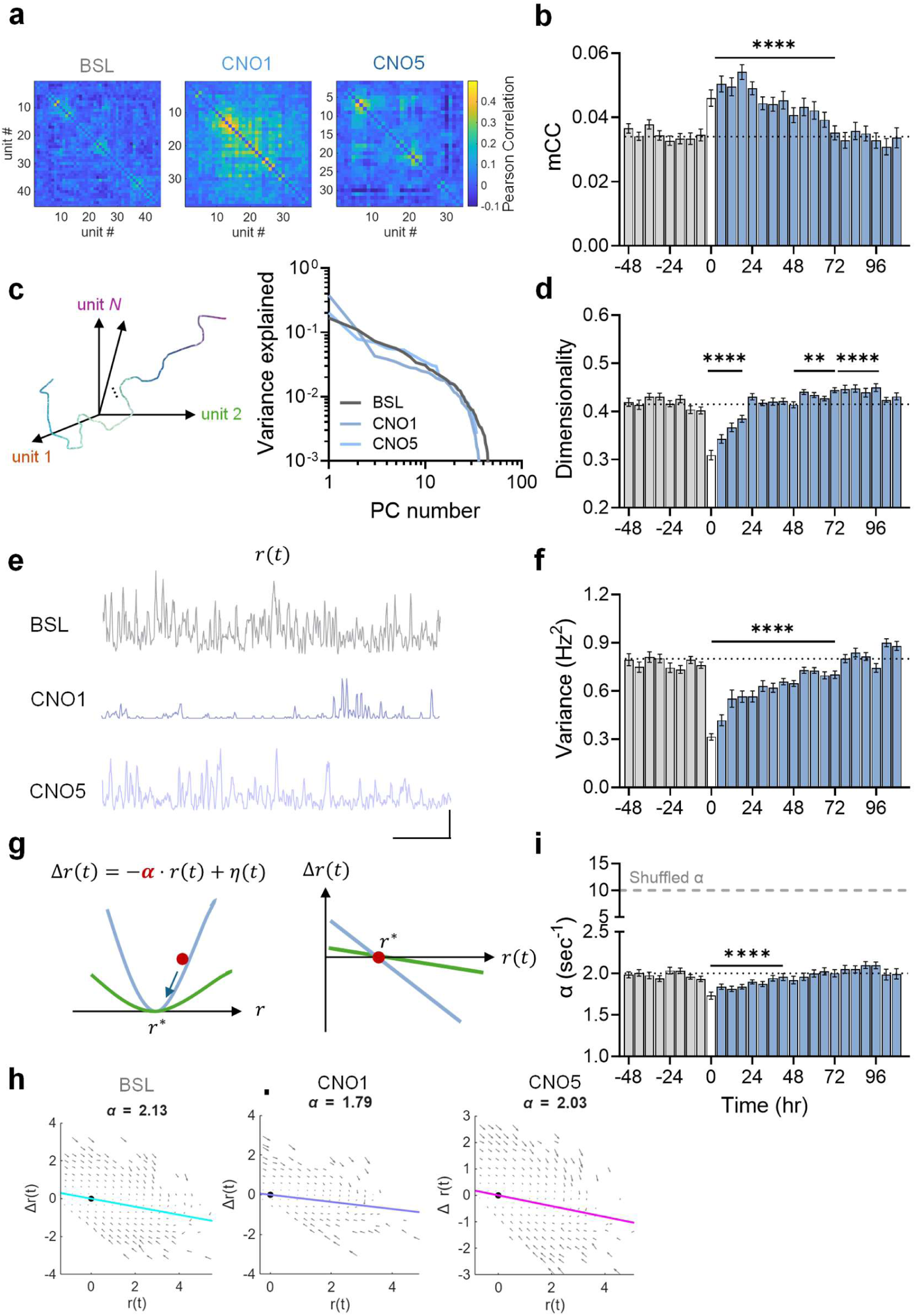
Regulation of CA1 state-space dynamics on fast and slow timescales. **a**, Representative pairwise correlation matrices from one mouse in a 15-min window during BSL, CNO day 1, and CNO day 5. **b**, Mean correlation coefficient (mCC) across active cells (15-min windows, *n* = 5 mice) during 48 h of BSL (grey) and after CNO (blue). mCC increased during CNO1–2 (\*\*\*\**P* < 0.0001) and returned to baseline by CNO5 (*P* > 0.99; pairwise contrasts in Table S2a). **c**, Population activity trajectories in neuronal state space (left) and variance explained by PCs for three example windows (right; same as in a). **d**, Linear embedding dimensionality (as a fraction of active units) across BSL and after CNO. Dimensionality decreased on CNO1 (\*\*\*\**P* < 0.0001), recovered by CNO2 (*P* > 0.99), overshoot during CNO3-4 (\*\**P* = 0.004 and \*\*\*\**P* < 0.0001, respectively) and recovered back to baseline by CNO5 (*P* = 0.235). Pairwise contrasts in Table S2b. **e**, Examples of population-averaged FR traces at 100-ms resolution from the same three windows (scale bars: 2 Hz, 100 ms). **f**, Variance of population-averaged FR (15-min windows, *n* = 5 mice) across BSL and after CNO. Variance decreased during CNO1–2 (\*\*\*\**P* < 0.0001) and recovered by CNO5 (*P* = 0.1; pairwise contrasts in Table S2c). **g**, Illustration of the quantification of dynamical stability: within each 15-min time window we model the instantaneous MFR, r(t), as a noisy linear process, the stability of which can be thought of as the steepness of an energy valley around its temporal mean (left). The steepness of the valley is estimated by the 1-dimensional flow maps: the slope of the change in r vs r itself (right). A steeper (negative) slope corresponds to higher stability. **h**, Example flow maps and corresponding α values from the same three windows. **i**, Stability index α (15-min windows, *n* = 5 mice) across BSL and after CNO, compared with shuffled controls (dashed). Stability decreased during CNO1–2 (\*\*\*\**P* < 0.0001) and renormalized by CNO5 (*P* > 0.99; Table S2d). In panels b, d, f, and i, 15-min windows were grouped into 6-h bins for statistics and visualization. The 6-h period immediately after pump implantation (white bar) was excluded. Time windows: BSL, –48 to 0 h; CNO1, 0– 24 h; CNO2, 24–48 h; CNO5, 96–108 h. Linear mixed-effects regression. Analyses were performed in 15-min windows, and statistical comparisons were conducted across consecutive 6 hr blocks. Data are presented as mean ± s.e.m..

Next, we characterized the complexity of population activity trajectories by analyzing population FR trajectories as they traversed state space (Fig. 3c, left). We did this by estimating the linear embedding dimensionality^77^ (Fig. 3c, right), defined as the number of principal components required to explain >80% of the variance in population dynamics, normalized to the number of active recorded units. Dimensionality was calculated after centering the data and is therefore independent of MFR changes. Following IN perturbation, dimensionality dropped sharply, returned to baseline within ∼24 hours, slightly overshot during the subsequent 2 days, and recovered back to baseline by the last day of perturbation (Fig. 3d). Together, the increase in pairwise correlations and the decrease in dimensionality indicate a network-level transient shift toward more synchronous, low-dimensional dynamics that is subsequently compensated.

We then analyzed fluctuations in the population-averaged FR (MFR) at high temporal resolution. The temporal variance of the MFR decreased significantly following IN perturbation and recovered after 3 days (Fig. 3e-f).

To assess stability more directly, we applied a dynamical systems approach, estimating the restorative forces that “pull” dynamic FR trajectories toward their mean (Fig. 3g; Methods). Specifically, we fitted a linear autoregressive model⁶⁶ to the temporal evolution of the mean FR and quantified the slope of Δr as a function of r, yielding the dynamical stability index α (Fig. 3h). Smaller α values correspond to weaker dynamical stability (Extended Data Fig. 7a-b). A representative flow map computed for Δr(t) as a function of r(t) for CA1 RS neurons showed a baseline stability index of α = 2.13 s⁻¹ (slope; Fig. 3h). Across mice, α was stable at baseline (1.98 ± 0.02 s⁻¹ compared with shuffled α values 9.8 ± 0.009 s⁻¹; Fig. 3i and Extended Data Fig. 7c). Shuffling time points within each 15-min window abolished structure in the flow fields and confirmed that α is not trivially explained by temporal correlations (Extended Data Fig. 7c). Unlike other FR measures, α showed a weaker response to perturbation: ∼10% decrease following hM3Dq-dependent IN activation (Fig. 3h-i). Similarly, hM4Di-dependent IN suppression resulted in a pronounced increase in the FR of RS units (Extended Data Fig. 8a-b) with only a ∼12% increase in dynamical stability (Extended Data Fig. 8c). Notably, variations in dynamical stability and FR levels need not covary. Indeed, intrahippocampal administration of the NMDAR open-channel blocker MK-801 led to a ∼60% decrease in the FR of RS units (Extended Data Fig. 8d-e), accompanied by a concurrent ∼20% increase in dynamical stability (Extended Data Fig. 8f), indicating that NMDAR-mediated excitatory currents normally destabilize spontaneous FR dynamics. Thus, dynamical stability increased following both an increase (Dlx-hM4Di) and a decrease (NMDAR blockade) in FR, suggesting that dynamical stability depends on synaptic receptor engagement rather than FR level per se.

Together, these results demonstrate that, in addition to MFR, multiple parameters of FR dynamics, including functional correlations, dimensionality, variance, and dynamical stability, are differentially sensitive to persistent IN activation and recover on distinct timescales.

### Regulation of state-dependent homeostatic set point by the perturbation

Sleep and wake states provide distinct neuromodulatory tones and network regimes, each characterized by distinct activity dynamics, raising the question of whether population-level parameters, such as MFR, are governed by state-specific set points. In other words, does each vigilance state independently modulate these parameters, or do they converge toward a shared stable target across states? Importantly, modulation by state alone is not sufficient to establish the existence of a stable set point; rather, stability must be demonstrable under perturbation. By examining FR dynamics during IN perturbation across vigilance states, we test whether state-dependent modulations converge back to their pre-perturbation values, thereby demonstrating the robustness of set-point regulation.

We first addressed the question of whether changes in network coordination and temporal structure were modulated at all by vigilance state. To do so, we quantified a range of statistical features of population activity, including FR variability, dimensionality, functional correlations, and dynamical stability during baseline across three states – AW, NREMS, and REMS (Fig. 4a). Multinomial logistic regression revealed that these features carried predictive information about vigilance states (Fig. 4b), with MFR contributing little unique predictive information (near chance; Methods, Fig. 4c). Among the features, pairwise correlations and α were most informative (Fig. 4c), uniquely contributing to identification of the three states and showing consistent state-dependent modulation (Extended Data Fig. 9). This agrees with previous studies showing that CA1 network dynamics display state-dependent modulation of correlations and stability across vigilance states^78^. Mean pairwise correlations were significantly higher during NREMS than in AW, with the lowest values during REMS (Extended Data Fig. 9a, Fig. 4d-f). These differences were stable across days, suggesting a persistent reorganization of functional interactions within the network by state. Dynamical stability was highest during NREMS relative to AW and REMS (Extended Data Fig. 9b, Fig. 4g-i), indicating state-dependent adjustments in intrinsic dynamics.

**Fig. 4.**
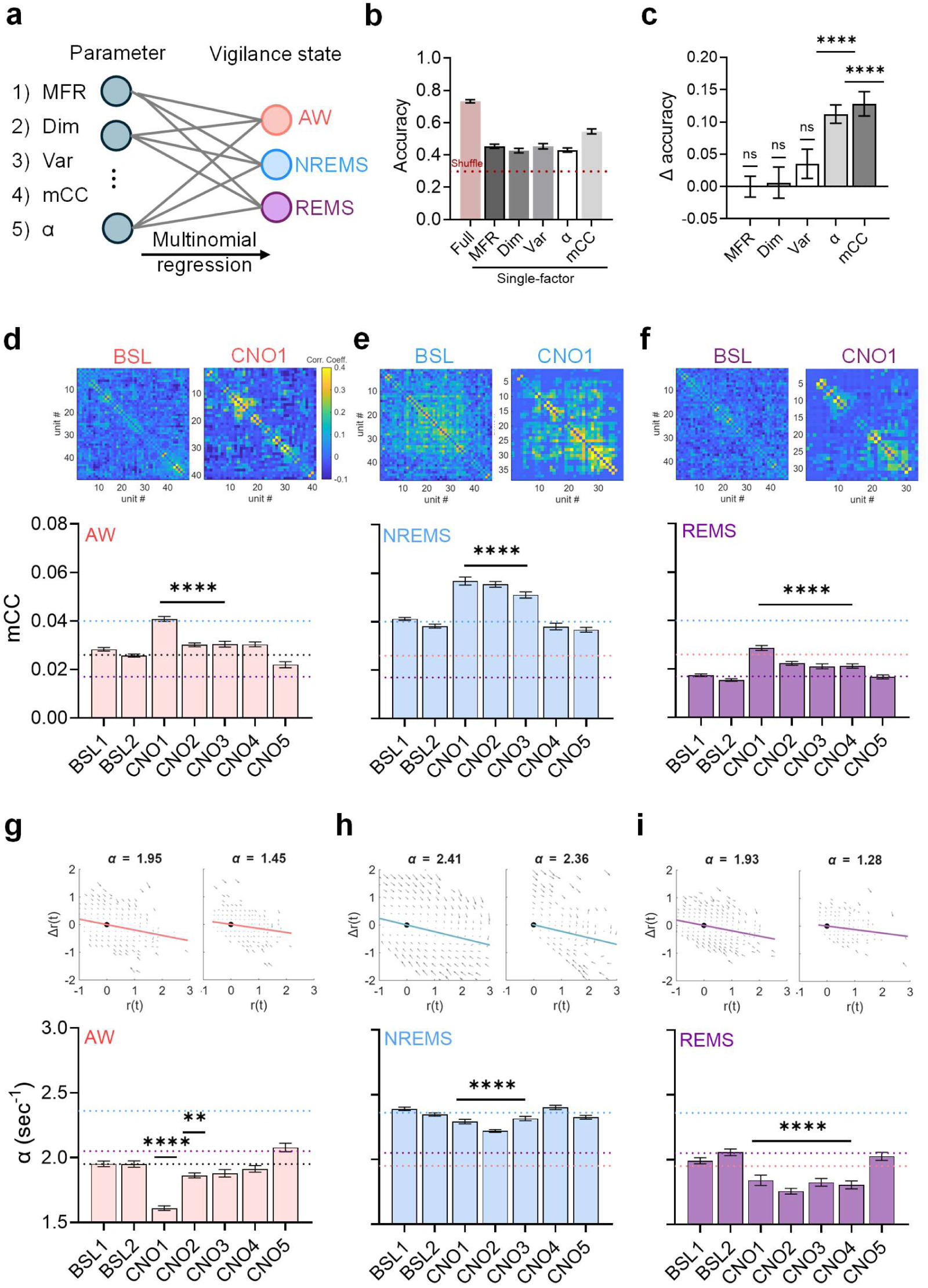
Pairwise correlations and dynamical stability exhibit state-specific set points that are stabilized in response to the perturbation. **a,** Schematic of multinomial regression model testing the contribution of single FR-derived parameters—mean firing rate (MFR), dimensionality (Dim), variance of mean FR (Var), mean correlation coefficient (mCC), and stability index (α) - to classification of vigilance states (AW, NREM, REM). **b,** Classification accuracy using each single parameter compared to the full model. c, Change in classification accuracy (Δ accuracy: full accuracy – accuracy of model omitting the parameter), showing mean correlation and α as the most informative single parameters for state discrimination (α and mCC: \*\*\*\**P <* 0.0001, variance: *P* = 0.18, dimensionality: *P* = 0.8, MFR: *P* = 0.99). **d-f,** Pairwise correlation structure across CA1 RS units (example matrices at BSL and in CNO day 1, top) and mean correlation coefficient (mCC, bottom) during baseline (BSL) and following CNO across AW (d), NREM (e), and REM (f). mCC increases significantly under IN perturbation across vigilance states (AW: CNO1-3 \*\*\*\**P* <0.0001; NREM CNO1-3 \*\*\*\**P* <0.0001; REM: CNO1-4 \*\*\*\**P* <0.0001; pairwise contrasts in Table S3a-c). **g-i**, State-specific analysis of dynamical stability (α). Example flow fields in BSL and in CNO day 1 (top) illustrate Δr(t) as a function of r(t), with slopes defining α. Population summary (bottom) shows that α is significantly reduced during AW (g) NREM (h) and REMS (i) and recover by CNO5 (AW: CNO1 \*\*\*\**P* <0.0001, CNO2 \*\**P* = 0.003; NREM: CNO1-3 \*\*\*\**P* <0.0001; REM: CNO1-4 \*\*\**\*P* < 0.0001; pairwise contrasts in Table S3d-f). One-sample Wilcoxon test (c), Linear mixed-effects regression (LMER; d-i); Data are presented as mean ± s.e.m.

Next, we asked whether pairwise correlations and the dynamical stability index α - the features that most contributed to vigilance-state classification - remain stable following perturbation. Pairwise correlations increased significantly following CNO administration in all states, then returned toward state-specific baselines, with the fastest recovery during wakefulness (Fig. 4d-f). In contrast, dynamical stability was robustly perturbed during wake and REMS, but the magnitude of the perturbation was modest during NREMS (Fig. 4g-i). Thus, among vigilance states, NREMS exhibits the highest dynamical stability and the lowest sensitivity to chemogenetic IN activation. Together, these findings indicate that functional pairwise correlations and temporal dynamics are stabilized around vigilance-state-specific set points in the face of the perturbation.

### Upward FR recovery is expressed preferentially during sleep

Previous studies suggest that sleep supports homeostatic regulation of synaptic structure and function, counterbalancing Hebbian-like plasticity that accumulates during wakefulness^60,79^. Conversely, an alternative hypothesis posits that FR compensation for a hypoactivity-inducing perturbation is suppressed during sleep but promoted during active wakefulness^27^. To address this controversy, we examined whether upward FR homeostasis in excitatory CA1 neurons occurs during specific brain states (Fig. 5a).

**Fig. 5.**
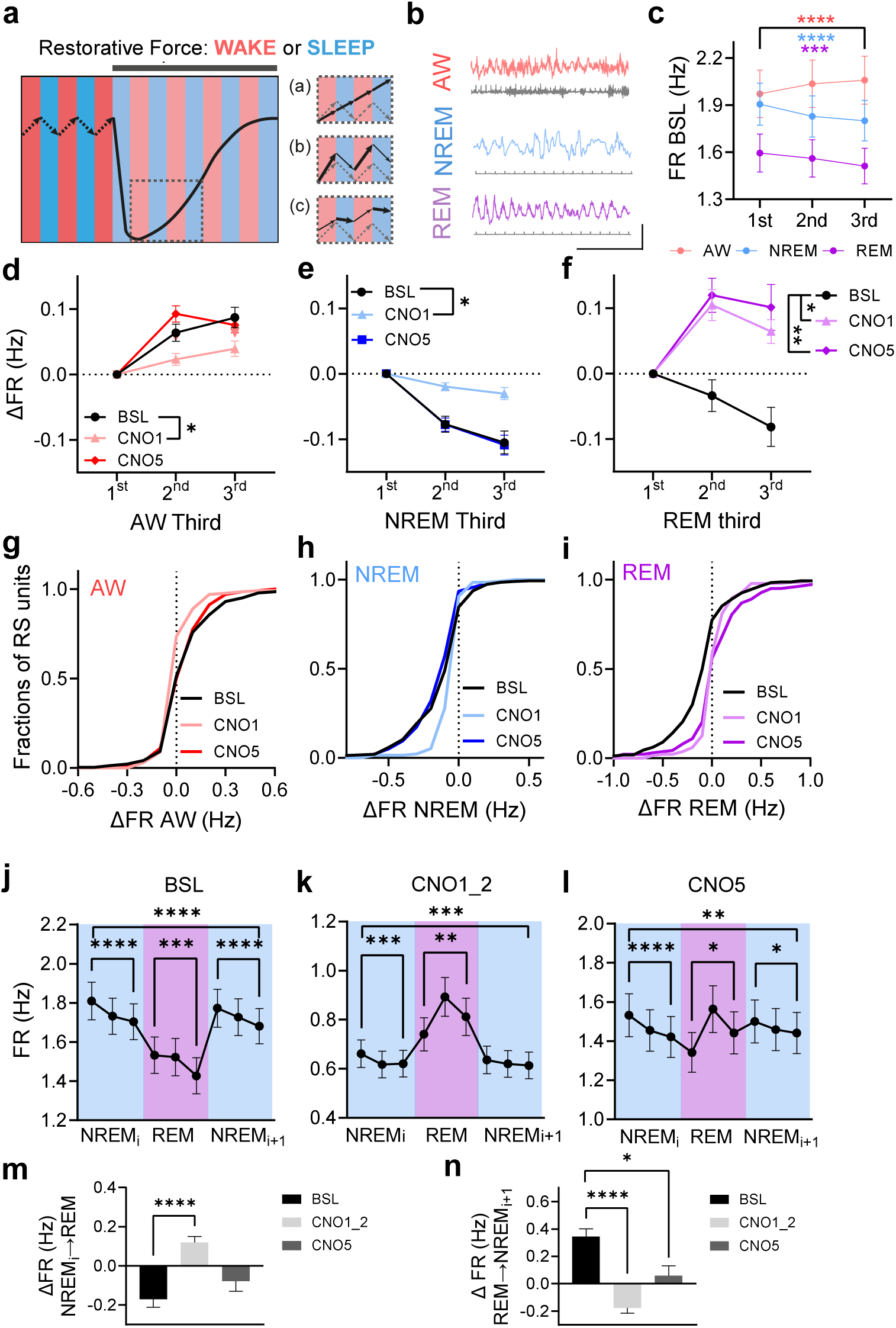
Homeostatic recovery of population FRs is preferentially expressed during sleep. **a**, Schematic illustration of hypothetical models of MFR recovery. The main plot shows baseline FR increases in wake (red) and decreases in sleep (blue), followed by MFR suppression at perturbation onset and subsequent recovery. Insets illustrate three possible recovery mechanisms: (a) continuous recovery across states, (b) recovery driven by enhanced increases during wake, or (c) recovery driven by reduced suppression during sleep. Gray arrows indicate baseline dynamics for reference. **b**, Representative CA1 LFP (top) and EMG (bottom) traces during AW (pink), NREMS (blue), and REMS (purple). Scale bars: 0.1 mV, 1 s. **c**, Baseline within-epoch FR dynamics of RS units per state. Baseline FR dynamics of RS units during AW show progressive increases (\*\*\*\**P* < 0.0001), FR dynamics during NREMS show decreases (\*\*\*\**P* < 0.0001) and FR dynamics during REMS show decreases (\*\*\**P* = 0.00011), pairwise contrasts in Table S4a. **d-f**, Average within-epoch FR change during AW, NREMS, and REMS. Data are plotted across epoch thirds (1–3) relative to the first third, with each line representing a single day. The last point corresponds to the within-epoch change used for statistical comparisons (last third – first third)**. d**, Average within-epoch FR change during AW exhibited a significant suppression on CNO1, (CNO1 \**P* = 0.016, CNO5 *P* = 0.7). **e**, Average within-epoch FR decrease during NREMS showed a significant attenuation on CNO1 (CNO1 \**P* = 0.015, CNO5 *P* = 0.34). **f**, Average within-epoch FR change during REMS shifts from within-epoch decreases at baseline to increases across all CNO days (CNO1: \**P* = 0.01, CNO5: \*\**P* = 0.005), pairwise contrasts in Table S4b. **g-i**, Cumulative distributions of within-epoch FR changes during AW, NREMS, and REMS. **g**, AW: (BSL vs CNO1 \*\*\*\**P* < 0.0001, BSL vs CNO5 *P* > 0.99). **h**, NREMS: (BSL vs CNO1: *P* < 0.0001; BSL vs CNO5: *P* = 0.052). **i**, REMS: (BSL vs CNO1 and BSL vs CNO5: \*\*\*\**P* < 0.0001). **j-l**, FR dynamics of RS units across NREM_i_–REM–NREM_i+1_ triplets computed per neuron across the first, middle, and last thirds of each state and averaged across sequences. **j**, baseline (BSL, 48 hr; 79 triplets; NREM_i_ *\*\*\*\*P* < 0.0001, REM \*\*\**P* = 0.0005, NREM_i+1_ \*\*\*\**P* < 0.0001; triplet_end-start_ \*\*\*\**P* < 0.0001). **k**, CNO days 1–2 (57 triplets; NREM_i_ *\*\*\*P* = 0.0008, REM \*\**P* = 0.003, NREM_2_ *P* = 0.08; triplet_end-start_ \*\*\**P* = 0.0005). **l**, CNO day 5 (90 triplets; NREM_i_ *\*\*\*\*P* < 0.0001, REM \**P* = 0.04, NREM_i+1_ \**P* = 0.03; triplet_end-start_ \*\**P* = 0.002), pairwise contrasts in Table S4c. **m,n**, Transition magnitudes in NREM_i_–REM–NREM_i+1_ triplets were quantified as the difference between the first third of the current state and the last third of the preceding state. Both the NREM_i_→REM and REM→NREM_i+1_ transitions shifted significantly on CNO1-2 (*P* < 0.0001). On CNO5, the NREM_i_→REM transition was not significantly different from baseline (*P* = 0.1), whereas the REM→NREM_i+1_ remained significantly reduced compared to baseline (*P* < 0.05), pairwise contrasts in Table S4d. Linear mixed-effects regression (LMER; c-f,j-n); Kolmogorov–Smirnov tests with Bonferroni correction (g-i). Data are presented as mean ± s.e.m across neurons.

We first analyzed sleep-wake architecture throughout the experiment (Fig. 5b, Extended Data Fig. 10). Given that CNO may have off-target effects on sleep^80^, we compared its effects in the Dlx-Cnt and Dlx-Gq groups across the duration of CNO pump delivery. The perturbation did not alter the fraction of total time spent in each state per day (Extended Data Fig. 10a) nor overall sleep-wake architecture (Extended Data Fig. 10b,c). Furthermore, no changes were observed in power spectral density across vigilance states (Extended Data Fig. 11), with the exception of a transient increase in slow-wave activity during NREMS in the Dlx-Gq group (Extended Data Fig. 11b).

The FR was uniformly suppressed by CNO delivery across all states and returned to state-specific baseline values (Extended Data Fig. 12a,b) as observed for global MFR regulation (Fig. 2d). To investigate the contributions of specific brain states to MFR recovery, we quantified within-epoch FR changes for each neuron as the difference between the last and first third of each epoch, after dividing the epoch into three equal segments (Extended Data Fig. 12c). These per-epoch changes were then averaged across all epochs of a given state (AW, NREMS, REMS) across a single day. Analyses of wake were restricted to AW due to the short duration of the majority of QW epochs. Transitions between states were defined as the difference between the first third of an epoch and the last third of the preceding epoch.

At baseline, RS units displayed the following FR dynamics and their mean value: increases during waking and an overall decrease during sleep (Fig. 5c). This fundamental pattern of opposing regulation across vigilance states has been reported in both hippocampus and neocortex^81–85^, highlighting that FRs undergo systematic modulation across the sleep-wake cycle. While our findings align with this general principle, we observed heterogeneous FR dynamics in CA1 mouse hippocampus during NREMS epochs (Fig. 5h), similar to data in mouse cortex^86^, with an overall average decrease in FR (Fig. 5c,e), in contrast to rat hippocampus where FR increases during NREMS epochs have been reported^82,84^.

Given these baseline FR dynamics, several mechanisms could drive MFR recovery in response to the perturbation, including FR increases that surpass baseline dynamics during wakefulness, reduced FR suppression during sleep, or both (Fig. 5a). State-specific FR analysis revealed distinct patterns of FR recovery following CNO administration. Within-epoch FR dynamics changed during MFR restoration in both wakefulness and sleep, but in qualitatively different ways. During AW, FR increases were initially attenuated but gradually returned to baseline levels over subsequent days (Fig. 5d,g). This effect remained consistent when normalizing the FR change by the averaged epoch FR (FR gain; Extended Data Fig. 12f), indicating the lack of compensatory wake-related FR enhancement during the perturbation, in comparison to its baseline within-epoch dynamics. In AW, the changes on CNO1 occurred mainly among units displaying within-epoch increases (Fig. 5g).

In NREMS, within-epoch FR suppression was initially attenuated on CNO1 and gradually returned to baseline levels (Fig. 5e,h), resulting in a smaller net FR decrease during this state. The changes in NREMS on CNO1 were mainly among units displaying within-epoch decreases (Fig. 5h), reflecting attenuated suppression. However, FR gain analysis revealed a more negative modulation on CNO1 (Extended Data Fig. 12f), indicating that the smaller absolute FR decrease during NREMS is partly accounted for by the lower overall firing imposed by the perturbation. This limits interpretation of the magnitude of the NREMS effect, although the net FR decrease accumulated across NREMS epochs is nonetheless reduced. Once MFR normalization was achieved, within-epoch FR dynamics in AW and NREMS returned to baseline patterns, underscoring the coordinated regulation of firing across vigilance states.

Most notably, REMS exhibited a dramatic reversal of baseline dynamics, shifting from FR suppression to sustained FR enhancement that did not fully renormalize (Fig. 5f,i). This pattern was preserved when FR changes were expressed as gain (Extended Data Fig. 12f), confirming a shift from baseline suppression to enhancement. CNO induced a rightward shift in the FR-change distribution, mainly among units showing within-epoch decreases, as in NREMS (Fig. 5i). Thus, the largest upward shift in within-epoch FR dynamics occurred during REMS, despite its baseline tendency toward downward within-epoch FR dynamics. These state-specific effects were robust to temporal binning, with similar results obtained using epoch thirds, fixed length start-end bins, or cumulative sums of within-epoch changes across all epochs (Extended Data Fig. 13a–c). This global analysis of NREMS and REMS epochs was further supported by the analysis of a subset of sleep episodes, namely NREMS–REMS– NREMS triplets^82,84^. Triplet analysis revealed similar baseline FR dynamics and similar modulation by CNO, with pronounced effects during REMS (Fig. 5j-l, Extended Data Fig. 14). Both the NREM_i_→REM (Fig. 5m) and REM→NREM_i+1_ (Fig. 5n) transitions shifted significantly during the acute phase of the perturbation. On CNO day5, the NREM_i_→REM transition was not significantly different from baseline, whereas the REM→NREM_i+1_ remained significantly reduced compared to baseline. Notably, the transition from NREMS to REMS, which showed a decrease in FRs under baseline conditions in agreement with earlier reports^82,84^, exhibited an increase in FR during CNO1–2, which could contribute to MFR recovery (Fig. 5m).

Together, these analyses indicate that the changes in within-epoch FR dynamics accompanying upward MFR recovery were restricted to sleep, whereas within-epoch increases during wake were instead transiently reduced. At baseline, AW was the only state exhibiting net upward within-epoch dynamics; these increases were not enhanced at any stage of the perturbation (Fig. 5d,g), arguing against a contribution of wake dynamics to recovery. In NREMS, the baseline downward dynamics were attenuated during the early phase of the perturbation, reducing the net FR decrease accumulated across NREMS epochs (Fig. 5e,h). In REMS, within-epoch dynamics reversed from net decreases at baseline to net increases and remained shifted throughout the perturbation, the largest deviation observed in any state (Fig. 5f,i). Both sleep states therefore deviated from their baseline dynamics in the direction of elevated firing, whereas wake dynamics did not. This dissociation indicates that upward MFR homeostasis is not a passive consequence of the baseline state-dependence of FR. Because NREMS occupies a far larger fraction of each day than REMS and comprises many more epochs (Extended Data Fig. 10a,c), the smaller per-epoch change in NREMS and the larger per-epoch change in REMS could both contribute to the net upward shift in MFR. Resolving their relative contributions will require state-selective manipulations.

### Intrinsic homeostatic plasticity of CA1 pyramidal neurons is induced by persistent IN perturbation

Having observed recovery of FR properties at the population level, we next investigated the underlying homeostatic mechanisms. In V1, homeostatic synaptic scaling in layers 4 and 6 elicited by monocular deprivation or dark rearing is restricted to an early critical period^54,56^. By contrast, adult layer 2/3 excitatory neurons exhibit either the absence^26^ or the presence^57,87^ of excitatory synaptic scaling, depending on the perturbation. Moreover, structural dendritic spine plasticity in V1 layer 2/3 neurons is likewise age-dependent^30^. Homeostatic regulation can also involve adjustments in intrinsic excitability^66,67,88^. However, although such changes contribute to homeostatic regulation of layer 2/3 excitatory neurons during the critical period^89^, they are not evident in adult mice^26,57^.

To examine the effect of persistent IN hM3Dq activation on intrinsic electrophysiological properties of CA1 pyramidal neurons, we implanted subcutaneous osmotic pumps delivering continuous CNO in Dlx-Cnt and Dlx-Gq mice. After five days of CNO delivery, hippocampal slices were prepared for patch-clamp recordings.

We elicited action potentials (APs) in response to increasing somatic current injections ranging from −200 to +500 pA and constructed firing–current (F–I) curves in the presence of postsynaptic receptor blockers. Following chronic CNO administration, CA1 pyramidal neurons from Dlx-Gq mice exhibited a significant leftward shift in the F–I relation compared with Dlx-control mice (Fig. 6a,b). Passive intrinsic properties, determined from the voltage responses to somatic current injections, such as resting membrane potential (RMP, Fig. 6c), input resistance (Fig. 6d), and AP threshold (Fig. 6e) were unaffected by the perturbation. Notably, the duration of spiking increased following the perturbation (Fig. 6f). These findings identify increased intrinsic excitability of CA1 pyramidal neurons as a candidate cellular mechanism that could contribute to homeostatic restoration of CA1 population firing after persistent IN activation.

**Fig. 6.**
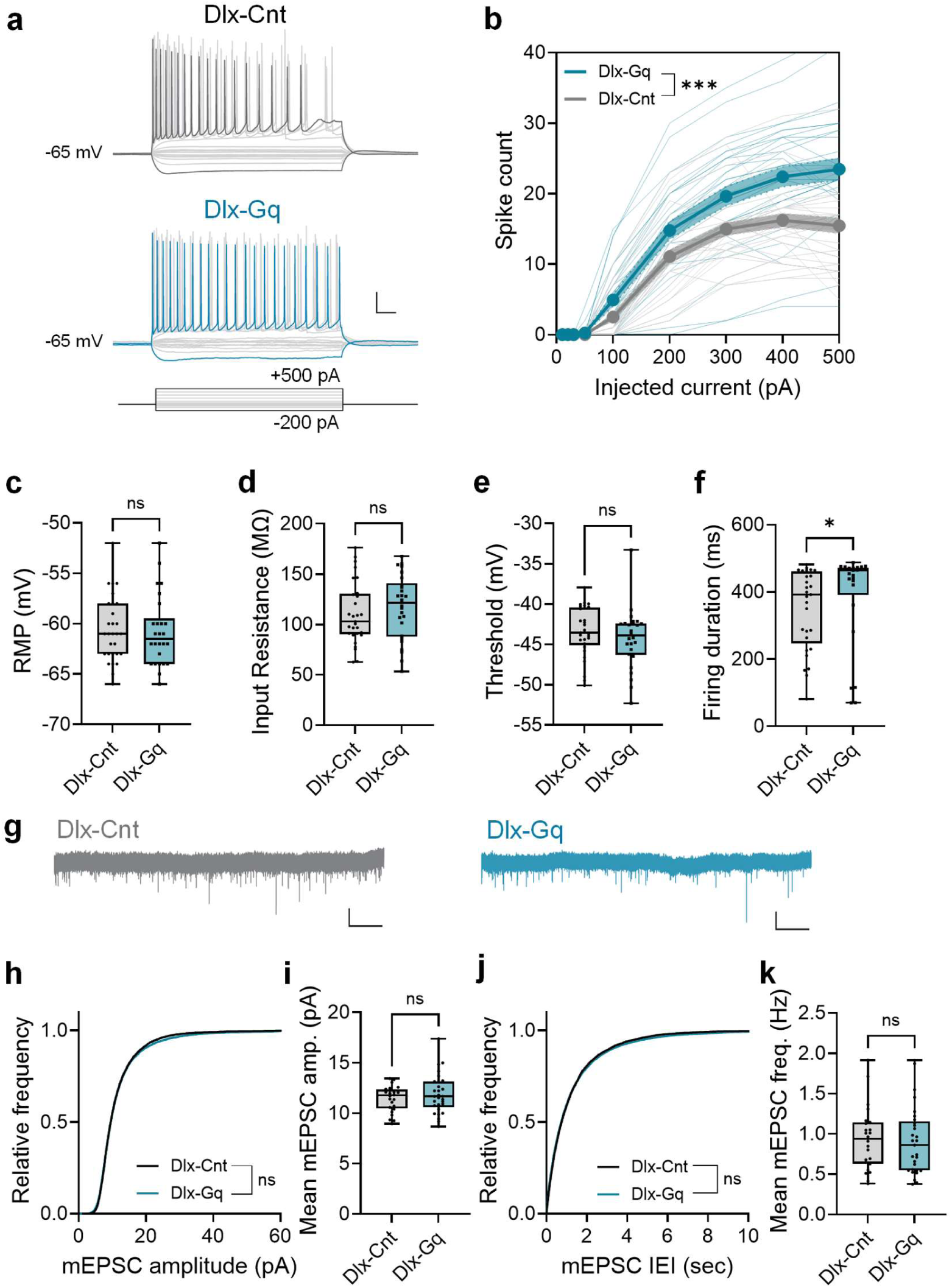
Chronic chemogenetic IN perturbation induces intrinsic but not synaptic homeostatic plasticity in CA1 pyramidal neurons. **a**, Representative traces evoked by step current injections in CA1 pyramidal neurons from Dlx-Cnt (top) and Dlx-Gq (bottom) mice after 5 days of CNO delivery via osmotic pump. Scale bars: 20 mV, 100 ms. **b**, F*–*I curves in CA1 pyramidal neurons from Dlx-Cnt (light gray) and Dlx-Gq (light blue) mice. Bold lines indicate mean ± SEM. Spike counts across depolarizing currents were higher in Dlx-Gq than Dlx-Cnt after 5 days of CNO (\*\*\**P* = 0.001, *n* = 31 neurons, 3 mice per group). **c-f**, Intrinsic electrophysiological properties of CA1 pyramidal neurons after 5 days CNO in Dlx-Cnt (*n* = 29 neurons, 3 mice) and Dlx-Gq (*n* = 26 neurons, 3 mice) groups. Group comparison: resting membrane potential (RMP, C, *P* = 0.84), input resistance (d, *P* = 0.45), spike threshold (e, *P* = 0.21), and firing duration (f, \**P* = 0.024). **g**, Representative traces of mEPSCs recorded from Dlx-Cnt (left) and Dlx-Gq (right) mice after 5 days of CNO. **h-k**, mEPSC analysis in Dlx-Cnt (*n* = 24, 3 mice) and Dlx-Gq (*n* = 29, 3 mice) groups. **h**, Cumulative distribution of mEPSC amplitude did not differ between Dlx-Cnt and Dlx-Gq groups (*P* = 0.16). **i**, Mean mEPSC amplitude did not differ between Dlx-Cnt and Dlx-Gq groups (*P* = 0.60). **j**, Cumulative distribution of mEPSC inter-event-interval (IEI) did not differ between Dlx-Cnt and Dlx-Gq groups (*P* = 0.08). **k**, Mean mEPSC frequency did not differ between Dlx-Cnt and Dlx-Gq groups (*P* = 0.74). Two-way ANOVA with Sidak’s multiple comparison test (b); unpaired Student’s t-test (c,d); Kolmogorov–Smirnov test (h,j); Mann-Whitney U test (e,f,i,k).

Having observed homeostatic plasticity of intrinsic excitability, we next asked whether synaptic homeostatic mechanisms also contributed to recovery of CA1 activity. To test this, we recorded mEPSCs from CA1 pyramidal neurons in hippocampal slices prepared from Dlx-Cnt and Dlx-Gq mice five days after CNO delivery (Fig. 6g). We observed no significant change in either mEPSC amplitude (Fig. 6h,i) or frequency (Fig. 6j,k) in slices from Dlx-Gq mice compared with Dlx-Cnt mice after five days of CNO delivery. These results provide no evidence for a contribution of quantal excitatory synaptic transmission to the homeostatic recovery of CA1 network activity in adult mice. Thus, under these conditions, intrinsic homeostatic plasticity of CA1 pyramidal neurons is the only candidate effector mechanism we detected, whereas changes in quantal excitatory transmission were not evident.

### Memory retrieval amidst homeostatic recovery

The observed recovery of RS CA1 network activity following IN perturbation is thought to be mediated by a wide range of homeostatic mechanisms^64–66,90^, such as an increase in intrinsic excitability, which can potentially be detrimental to some hippocampal functions^13,50^ and can interfere with memory storage or retrieval by rendering previously stored information inaccessible. A key question is whether homeostatic mechanisms can preserve the hippocampus’s ability to retrieve stored memories while simultaneously stabilizing network activity in response to perturbation^91^. Specifically, do the mechanisms that renormalize network activity in CA1 circuits also restore CA1-dependent memory retrieval, or might they compromise critical cognitive functions despite reestablishing baseline activity levels?

To investigate how homeostatic processes affect the stability and accessibility of hippocampal memory, we focused on retrieval of contextual fear memory^92^ (CFM, Fig. 7a). We examined whether CA1-dependent CFM retrieval^93,94^ is maintained during chemogenetic IN perturbation and amidst FR homeostasis. In a separate cohort of Dlx-Gq and Dlx-Cnt mice, we designed the following behavioral experiment (Fig. 7a): four weeks after AAV injections into dorsal CA1, both groups underwent contextual fear conditioning (memory formation phase) and were tested for retrieval at three time points — at baseline (Retrieval 0), on the first day of CNO application (Retrieval 1), and on the fifth day of CNO application (Retrieval 2) — in the original context (CtxA) and a novel context (CtxB).

**Fig. 7.**
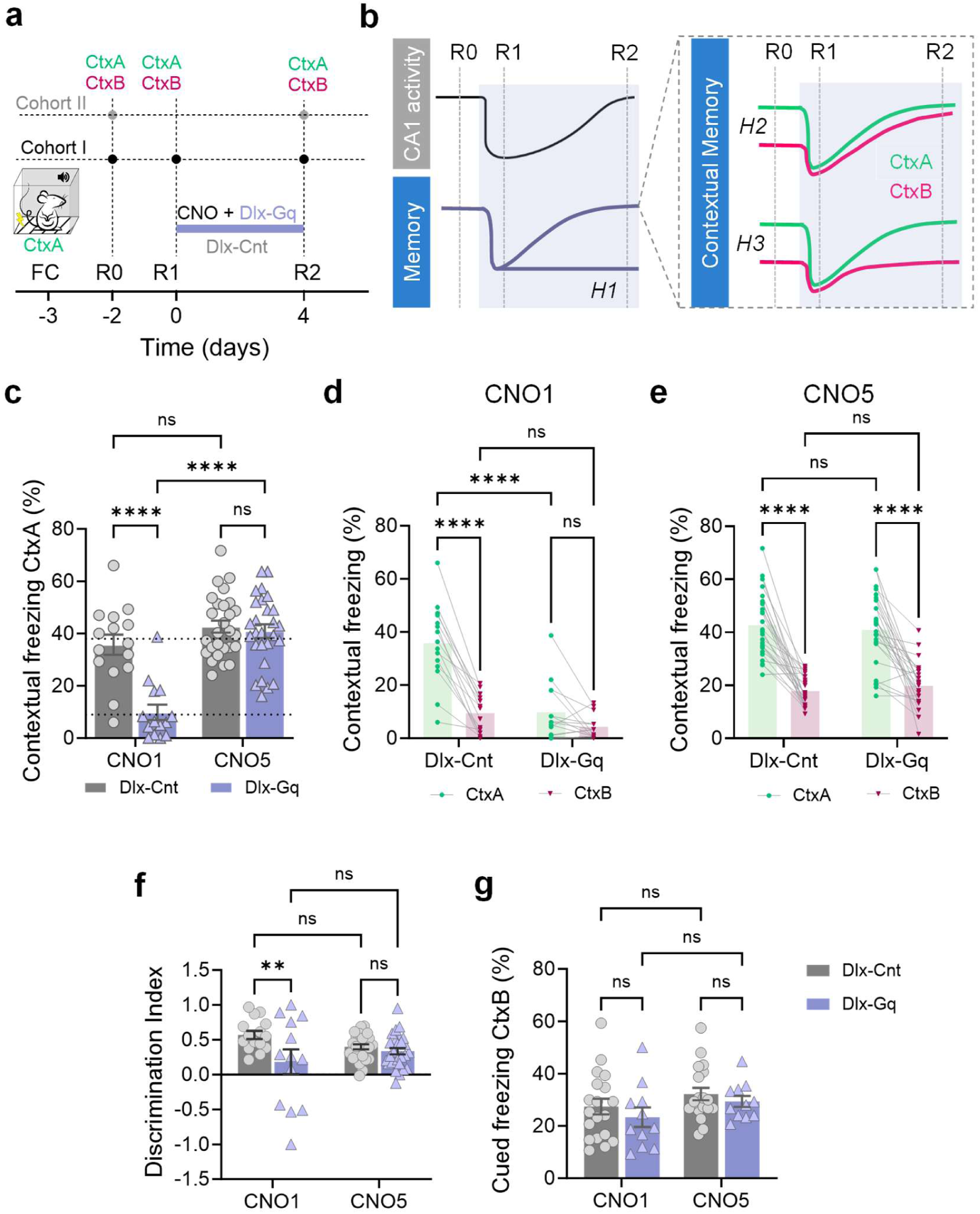
CA1-dependent contextual fear memory retrieval is impaired and subsequently recovers during persistent chemogenetic IN perturbation. **a,** Experimental design: Dlx-Gq and Dlx-Cnt mice underwent fear conditioning (FC) in Context A. Retrieval sessions were conducted in Context A (CtxA) and Context B (CtxB). Cohort II did not undergo Retrieval 1 (R1) at CNO1. **b,** Schematic illustration of hypotheses for memory retrieval outcomes: (H1) No recovery of retrieval, memory remains inaccessible; (H2) Recovery with impaired context discrimination, leading to generalization; (H3) Recovery with preserved context discrimination, maintaining specificity. **c,** Contextual freezing in CtxA for Dlx-Gq and Dlx-Cnt groups at Ret1 (CNO1) and Ret2 (CNO5). At Ret1: Dlx-Gq (*n* = 14) vs. Dlx-Cnt (*n* = 15); at Ret2: Dlx-Gq (*n* = 27) vs. Dlx-Cnt (*n* = 27); Dlx-Gq vs. Dlx-Cnt at CNO1: \*\*\*\**P* < 0.0001; at CNO5: *P* = 0.62; Dlx-Gq CNO1 vs. CNO5: \*\*\*\**P* < 0.0001; Dlx-Cnt CNO1 vs. CNO5: *P* = 0.1. **d**, Contextual freezing in CtxA and CtxB for Dlx-Gq and Dlx-Cnt at Ret1 (CNO1). Dlx-Cnt CtxA vs. CtxB: \*\*\*\**P* < 0.0001; Dlx-Gq CtxA vs. CtxB: *P* = 0.07; CtxA Dlx-Cnt vs. Dlx-Gq: \*\*\*\**P* < 0.0001; CtxB Dlx-Cnt vs. Dlx-Gq: *P* = 0.182. **e**, Same as in d for Ret2 (CNO5). Dlx-Cnt CtxA vs. CtxB: \*\*\*\**P* < 0.0001; Dlx-Gq CtxA vs. CtxB: \*\*\*\**P* < 0.0001; CtxA Dlx-Cnt vs. Dlx-Gq: *P* = 0.54; CtxB Dlx-Cnt vs. Dlx-Gq: *P* = 0.47. **f**, Discrimination index for Dlx-Gq and Dlx-Cnt groups at Ret1 (CNO1) and Ret2 (CNO5), calculated as (freezing in CtxA − freezing in CtxB) ÷ (freezing in CtxA + freezing in CtxB). Dlx-Cnt vs. Dlx-Gq at CNO1: \*\**P* = 0.002; at CNO5: *P* = 0.46; Dlx-Cnt CNO1 vs. CNO5: *P* = 0.1; Dlx-Gq CNO1 vs. CNO5: *P* = 0.16. **g**, Cued freezing for Dlx-Gq (n = 11) and Dlx-Cnt (n = 19) groups at Ret1 and Ret2. Dlx-Gq vs. Dlx-Cnt at CNO1: *P* = 0.34; at CNO5: *P* = 0.51; Dlx-Gq CNO1 vs. CNO5: *P* = 0.21; Dlx-Cnt CNO1 vs. CNO5: *P* = 0.19. Two-way ANOVA with Fisher’s LSD (c-g), Data are presented as mean ± s.e.m across mice.

Indeed, one day after the acquisition phase (Retrieval 0), Dlx-Gq and Dlx-Cnt mice showed comparable freezing during CFM retrieval which was significantly lower in a novel context in comparison to a familiar context (Extended Data Fig. 15a), indicating that learning and memory in the hippocampus, including encoding and consolidation processes, were equivalent between groups, and that memory retrieval was context-specific prior to the perturbation.

Next, three days after the acquisition, osmotic pumps were implanted to deliver CNO for five days. Previous studies have shown that acute suppression of CA1 network activity - whether by directly silencing pyramidal neurons^94^ or by activating parvalbumin-positive CA1 interneurons^95^ - impairs memory retrieval. As expected, on the first day of the perturbation, when MFR of CA1 RS neurons was maximally suppressed, the Dlx-Gq group exhibited reduced freezing compared to the control group (Fig. 7c), indicating impaired CFM retrieval.

Does memory retrieval remain impaired during persistent IN perturbation? We evaluated memory retrieval on CNO5 following MFR renormalization, to test the following competing hypotheses (Fig. 7b): (1) Failure to recover retrieval of memory specific to the conditioned context; (2) Recovery of conditioned-context memory retrieval at the cost of memory specificity; (3) Restoration of both the ability to retrieve the conditioned context and distinguish between contexts, consistent with memory resilience.

Remarkably, freezing in the Dlx-Gq group recovered by the last day of the perturbation (Fig. 7c). Notably, the ability to discriminate novel (CtxB) and familiar (CtxA) contexts was not impaired when network activity recovered (CNO5), remaining indistinguishable between Dlx-Gq and Dlx-Cnt groups (Fig. 7d-f). This finding indicates that maintaining memory strength did not come at the cost of losing context specificity^96^. Furthermore, memory retrieval remained indistinguishable from that of the Dlx-Cnt after CNO washout (Extended Data Fig. 15b). To rule out the possibility that impaired Retrieval 1 on CNO1 could affect Retrieval 2 on CNO5, we added another cohort of mice (cohort II) that did not undergo Retrieval 1 (Fig. 7a). Freezing on CNO5 showed no difference between Dlx-Gq and Dlx-Cnt mice, regardless of whether they underwent both Retrieval 1 and 2 or only Retrieval 2. A mild reduction in freezing was observed in Dlx-Gq mice that experienced both retrieval sessions compared with those with only Retrieval 2 (Extended Data Fig. 15c), consistent with an effect on memory reconsolidation^97^. Combined results (Cohort I and Cohort II) show that freezing recovered to control levels in Dlx-Gq group (Fig. 7c-f).

To ensure that CFM retrieval remained hippocampus-dependent throughout the experiment^94,98^, we tested a separate group of mice for CFM retrieval seven days after the first retrieval. Mice were injected intraperitoneally with CNO or saline 30 minutes before retrieval. As expected, CNO-injected mice exhibited impaired retrieval compared to saline-injected mice (Extended Data Fig. 15d), confirming the continued involvement of the hippocampus in CFM retrieval throughout the experimental timeline.

To verify the spatial specificity of the perturbation, we tested mice for auditory-cued memory, which is known to be dependent on the amygdala but not on the hippocampus^99^. Chemogenetic activation of local hippocampal INs had no effect on cued fear memory in a novel context (Fig. 7g and Extended Data Fig. 15e). Additionally, this perturbation did not influence exploratory behavior in the open field, as evidenced by no differences in the total distance traveled, velocity, or percentage of time spent in the center between the Dlx-Gq and Dlx-Cnt groups on the first day of the perturbation (Extended Data Fig. 15f-h).

Overall, these results suggest a temporal link between the recovery of CA1 network activity and the recovery of memory retrieval, indicating that access to previously stored memories can be restored without loss of context specificity during persistent perturbation.

## Discussion

By tracking hippocampal networks during a sustained inhibitory perturbation, we find that CA1 network activity recovery is multivariate, state-resolved, and behaviorally meaningful. Intrinsic homeostatic plasticity in pyramidal neurons coincides with coordinated renormalization of RS population-level statistics toward vigilance-state–specific set points, and retrieval of previously stored memories recovers without loss of specificity. Together, these results provide a framework for understanding hippocampal resilience as structured, state-dependent regulation of network dynamics, rather than simple rate renormalization, consistent with preserved cognitive function.

A fundamental role of memory retrieval is to ensure the seamless transmission of information across time. Yet how homeostatic mechanisms elicited by ongoing perturbations operate in the hippocampus, and how they interact with this critical hippocampal function, remains largely unknown. We find that homeostatic mechanisms in the CA1 hippocampus not only stabilize neuronal networks composed of highly dynamic single neurons^100–102^, but can coincide with the restoration of access to seemingly ‘inaccessible’ memories^103–105^. This recovery from transient amnesia is particularly striking because it occurs during periods associated with homeostatic plasticity that might otherwise interfere with memory functions^13,40^. Although classic homeostatic models emphasize firing-rate stabilization, whether such mechanisms safeguard higher-order functions has remained an open question^106^. Here, we show that network-dynamics homeostasis in CA1 is compatible with preserved memory accessibility during persistent perturbation.

Homeostatic processes operate on timescales of hours to days, whereas behavior unfolds over seconds to minutes^106^. To understand the balance between homeostatic processes that maintain dynamic stability and the network functions they support, it is crucial to unpack the long-term dynamics of the population mean FR across both spatial and temporal dimensions. Our analyses show that the variance of the MFR and its dynamical stability were perturbed and subsequently recovered in response to elevated inhibitory drive. While dynamical stability showed only mild sensitivity to chemogenetic modulation of INs, it was highly sensitive to NMDA receptor–dependent conductance. In addition, our findings reveal that both dimensionality and pairwise correlation strength are sensitive to an increase in inhibitory drive and display precise homeostatic recovery, preserving the complexity of firing dynamics. These higher-order population statistics – dimensionality and correlation – have been proposed as indicators of learning efficiency^107^. Whether the homeostatic regulation of dimensionality and correlations contributes to memory resilience by enabling multiple functional circuit configurations amidst homeostatic forces remains to be explored in future studies.

To elucidate the underlying principles of FR homeostasis in the hippocampus *in vivo*, we explored the role of sleep in the homeostatic recovery of hippocampal activity. Our findings indicate a dual role for sleep in homeostatic control: it shapes both the set points and the regimes in which effector mechanisms are expressed. At the level of set-point regulation, analyses revealed bidirectional coupling between vigilance states and specific features of FR dynamics, such as dynamical stability and functional pairwise correlations. These parameters not only differ at baseline but also renormalize to state-specific set points following the perturbation, with recovery kinetics that differ across brain states. At the effector level, homeostatic recovery of CA1 MFR is preferentially expressed during sleep following persistent IN activation. While FR increases occurring during AW did not exceed baseline dynamics and therefore did not contribute to recovery beyond normal wake-associated changes, upward FR recovery toward baseline was most prominently expressed during REM sleep. FR dynamics at NREM-to-REM transitions and during REM epochs exhibited a robust shift from net decreases to net increases following chemogenetic IN perturbation. These results imply that intra-state FR dynamics are not fixed but are instead modifiable, providing a potential framework for understanding the high heterogeneity of intra-state FR dynamics reported in the literature^27,81,82,84,85^. Further studies are required to establish a causal role of REMS in homeostasis of hippocampal activity and function to test if it extends its established functions in memory-related processes, including learning^108,109^, consolidation^110–112^, and forgetting^15,113^. Speculatively, the suspension of thermoregulation during REMS^114^, one of the most energetically costly peripheral processes in endotherms^115^, may bias network-level recovery toward this state, consistent with an energy-allocation framework^116^.

An important question raised by our findings is whether homeostatic regulatory mechanisms are universal across brain regions or instead adapted to the functional requirements of specific circuits. While network-wide FR statistics and functional pairwise correlations appear homeostatically regulated in both CA1 and V1^24–27,29,117^, the underlying mechanisms differ substantially between regions. Specifically, FS neuron populations did not renormalize in adult CA1 in our work, mirroring adult V1^26^ but contrasting with juvenile V1^24^. Furthermore, in juvenile V1, FR recovery is gated during active wakefulness in response to monocular deprivation during the critical period^27^, whereas in CA1, the upward homeostatic FR response was preferentially expressed during sleep following chemogenetic IN perturbation. Moreover, whereas chemogenetic suppression of neurons in adult V1 elicits synaptic, but not intrinsic, homeostatic plasticity^57^, persistent chemogenetic IN activation in adult CA1 selectively upregulates intrinsic excitability in pyramidal neurons (Fig. 6), consistent with homeostatic plasticity observed in adult CA1 pyramidal neurons following persistent blockade of sodium channels in vivo^58^. Further studies are required to assess potential contributions of inhibitory synapses and neurons to the renormalization of CA1 network dynamics following chemogenetic IN activation. Together, these findings support the view that homeostatic regulation employs distinct cellular mechanisms depending on brain region, perturbation, and developmental stage, underscoring the need for continued investigation to determine which principles are broadly generalizable and which are context-dependent or shaped by circuit architecture.

Our work emphasizes the importance of temporal resolution when studying neural circuit functions. Homeostatic mechanisms that restore mnemonic processes necessitate careful consideration of perturbation duration and timing in behavioral assessments. Genetic tools allow for cell-type and region-specific gene expression control^118^ but typically induce effects over extended periods, permitting ample time for homeostatic compensation. Consequently, the absence of behavioral deficits in these contexts may indicate compensation rather than a lack of impact. On the other hand, pharmacological inhibitors allow for rapid assessments but lack target specificity, whereas optogenetics offers fast, cell-type-specific modulation without enabling long-term assessment. Determining necessity therefore requires designs that separate a mechanism’s immediate contribution to mnemonic processing from its susceptibility to homeostatic compensation over time.

We considered three hypotheses to describe how physiological and behavioral responses are coordinated to meet homeostatic challenges in hippocampal circuits:

1. *Unstable memory as a trade-off for network homeostasis*: This hypothesis posits that restoring network excitability may come at the expense of hippocampal functions, such as memory formation, consolidation, or retrieval. While this trade-off might preserve overall network stability, it may lead to specific functional failures.
2. *Trade-off between memory stability and specificity*: In this scenario, homeostatic mechanisms restore general stability at the cost of memory precision, leading to memory generalization. Specific features of memories are sacrificed to maintain overall network activity. For example, disruptions to homeostatic synaptic scaling^119^, loss of presynaptic GABA(B) receptors^120^, or reduced GABA release from FS INs under stress^121^ are all associated with a loss of memory specificity.
3. *Cognitive resilience hypothesis*: This hypothesis suggests that homeostatic mechanisms can fully restore memory retrieval, maintaining both strength and specificity. Such resilience underscores the brain’s ability to adaptively resolve disruptions while preserving critical functions.

Our experiments in wild-type adult mice support the cognitive resilience hypothesis: homeostatic recovery of network dynamics coincides with preserved accessibility and context-specific precision of previously stored memories despite persistent perturbation of local inhibition. Network-level homeostasis compensates for sustained disruptions, parallel to recovery of memory access without loss of fidelity. This resilience was demonstrated for retrieval of a memory acquired before the perturbation. Stages that depend on fast-spiking, parvalbumin-positive interneurons, such as consolidation and learning^78,122,123^, may be less resilient, since FS-unit activity did not renormalize during persistent IN perturbation (Fig. 2e). We also observe intrinsic homeostatic plasticity in CA1 pyramidal neurons, however, establishing its necessity or sufficiency for behavioral recovery will require pathway-specific manipulations that isolate homeostatic mechanisms from Hebbian plasticity, an important but challenging goal. Notably, intrinsic excitability has been extensively implicated in memory allocation, with more excitable neurons more likely to be recruited into memory ensembles and shaping ensemble interactions and memory linking processes across time^124–130^. It will therefore be interesting to test whether learning during CNO5, when intrinsic excitability is elevated relative to baseline, differs from controls, providing a tractable avenue to explore additional functional implications of homeostatic plasticity for learning. This work lays a foundation for understanding how homeostatic processes preserve memory retrieval in dynamic neural systems. It motivates theoretical and experimental tests of how population-level statistics and dynamical stability interact across vigilance states, how sleep constrains effector mechanisms, and how memory remains accessible amidst ongoing homeostatic regulation.

## Methods

### Experimental animals

All procedures were approved under a Tel Aviv University (TAU) Institutional Animal Care and Use Committee protocol or by the government of Saxony-Anhalt under the license 203.m-42502-2-1737 LIN_GÜ. All experiments were performed on 4-6 month-old C57BL/6 male mice (Envigo, Israel; Charles River, Germany). All animals were kept in a normal light/dark cycle (12 h /12 h, lights on at 7AM), at room temperature (∼21-23 °C) and had free access to food and water. Mice were singly housed after the implantation of tetrodes.

### Viruses

We used sAAV-5/2-hDlx-HBB-chI-HA_hM3D(Gq)_2A_NLS_dTomato (Dlx-Gq group, v128, ETH Zurich Viral Vector Facility) or the control viral vector sAAV-5/2-hDlx-HBB-chI-HA_NLS_dTomato (Dlx-Cnt group, made by ETH Zurich Viral Vector facility).

### Stereotaxic surgery for viral injections

Mice were initially anesthetized with 5% isoflurane by volume for induction, followed by an intraperitoneal injection of ketamine/xylazine (100 mg/kg ketamine and 8 mg/kg xylazine). The animals were then head-fixed in a stereotaxic apparatus (David Kopf Instruments) and maintained under anesthesia via continuous inhalation of isoflurane (0.5–1%) throughout the surgery. Eye ointment (Duratears, Vetmarket) was applied to prevent corneal dehydration, and body temperature was monitored and maintained at 34°C using a heating pad (FHC, DC temperature controller). At the onset of the procedure, mice received a subcutaneous injection of Carprofen (5 mg/kg) to minimize inflammation and pain. A small craniotomy was performed, and 500 nL (at 0.1 μl/min flow rate) of viruses were injected bilaterally into the CA1 pyramidal at the following coordinates: −2.18 mm AP, −1.5 mm ML, and −1.3 mm DV to bregma. The incision was sutured, disinfected with Betadine, and the mice were given a subcutaneous injection of buprenorphine (0.02 mg/kg) for postoperative analgesia.

### Electrophysiological recordings in behaving mice

#### Tetrode implantation

Two to three weeks after virus injection, mice were implanted with a custom-made microdrive (custom printed circuit board and drive, Rogat, Carmiel, Israel), along with neck muscle electrodes for EMG and frontal EEG screw. The microdrive contained a movable assembly of four tetrodes, which were connected to the recording setup via an Omnetics head stage connector (Connector Corporation, Minneapolis, MN, USA). Tetrodes were constructed from four twisted 17-μm polyimide-coated platinum-iridium (90%–10%) wires (California Fine Wire) and mounted in groups of four into microdrives with a single turning screw and no separation between tetrodes. The electrode tips were plated with platinum to reduce electrode impedances to 150–300 kΩ at 1 kHz.

#### Surgical procedures

Mice were anaesthetized by continuous isoflurane inhalation and head fixed to a stereotaxic apparatus. The animal’s eyes were covered in ointment (Duratears, Vetmarket) and body temperature was maintained at 34°C by a heating pad (FHC, DC temperature controller). Four holes were drilled in the skull: above the right and left cerebellum for ground screws, above the left frontal cortex for an EEG screw (1.5 mm anterior, 1.5 mm lateral to bregma) and another above the right dorsal CA1 for the tetrode microdrive (1.94 mm AP, 1.5 mm ML, 1 DV to bregma). All craniotomies were drilled with a 0.7 mm burr. Bipolar stainless-steel electrodes (0.127 mm diameter) were implanted in the cervical muscles for EMG recordings. All implants were secured to the skull with dental cement and to each other with light curing glue to reduce the construct weight. After surgery mice were injected subcutaneously with 0.125 mg/kg carprofen and 0.05 mg/kg buprenorphine. Mice were allowed to recover in their home cage for 5-7 days before additional handling or experimentation. During this time the animal’s weight and overall wellbeing were monitored daily.

#### Tetrode positioning and mice habituation

Following postoperative recovery, mice were housed in recording cages with *ad libitum* access to food and water and connected to the recording system. Tetrode microdrives were lowered once daily in 25–100 μm increments, with 24-hr interval between adjustments, until the CA1 stratum pyramidale was reached (typically within 1–2 weeks). During this period, mice were habituated to the recording apparatus, cage, room, and experimenter. CA1 stratum pyramidale was identified by (1) calculated implant depth, (2) ripple oscillations during NREM sleep, and (3) a high ratio of RS to FS units (see below). Additional depth indicators included the transition from cortex to white matter (low spiking activity) and from white matter to CA1 stratum oriens (increased FS activity). Experimental recordings commenced only after tetrodes reached the stratum pyramidale. At the conclusion of the experiment, mice were anesthetized and small electrolytic lesions (30 μA, 20 s) were made to allow histological verification of tetrode placement.

#### Data acquisition

Following recovery period, the mice were transferred to the experimental room and connected to the recording system. Raw data were sampled at 24 kHz using a Neurophysiology Workstation (RZ5D base processor and PZ5 NeuroDigitizer amplifier, Tucker-Davis Technologies Inc.). Once the CA1 pyramidal layer was targeted, the experimental protocol commenced. Mice were recorded in their familiar home cages for 48 hours. On the third day, during the first half of the light cycle, the mice underwent implantation of an osmotic pump containing CNO (see below). Mice were then returned to their home cages and reconnected to recording system for continuous monitoring over the following five days. On the sixth day, the pump was removed under isoflurane anesthesia and mice were subsequently recorded for an additional 48 hours.

#### Chronic CNO delivery

Mice were anesthetized with isoflurane (1–2%), and a mid-scapular incision was made on the back of the animal to implant subcutaneously an osmotic pump (ALZET® pumps, 1007D model). The pump, pre-loaded with clozapine N-oxide dihydrochloride (Tocris, cat. no. 6329) and pre-incubated overnight to ensure immediate drug delivery, was implanted to deliver CNO at a constant rate of 0.5 mg/kg per hour. Following implantation, the incision was sutured, disinfected, and lidocaine cream was applied for analgesia.

#### Intraperitoneal CNO injection

For the experiments involving intraperitoneal injection of CNO, CNO (AK Scientific) was dissolved in dimethyl sulfoxide (DMSO) to a concentration of 5mg/ml and then further diluted in PBS to a concentration of 0.5 mg/ml. CNO was injected intraperitoneally 40 min to 1h before testing, at 5 mg/kg.

#### Scoring of vigilance states

EEG, LFP and EMG signals were subsampled from the raw data to 1250 Hz by a low-pass sinc filter with a cutoff frequency of 450 Hz. EMG signals were also high-pass filtered at 10 Hz. Vigilance states were classified in 1 sec epochs as active wake (low-amplitude de-synchronized EEG with high EMG activity), quiet wake (low-amplitude de-synchronized EEG with tonic EMG absent of movement-associated bursts), NREMS (high-amplitude synchronized EEG at low frequencies (<4 Hz) with low EMG activity), and REMS (prominent EEG activity in the theta band (4-10 Hz) with low EMG activity). Classification occurred semi-automatically using the AccuSleep MATLAB package^131^ with subsequent manual curation. Brief awakenings (wake periods shorter than 16 s occurring within NREMS episodes) were identified and removed, along with the corresponding spikes, without terminating the surrounding NREMS episode, the remaining NREMS segments were merged. Similarly, quiet wake epochs shorter than 16 s were merged with adjacent active wake epochs. Only state epochs longer than 60 s were included in the analyses.

For analyses of firing rate and network properties, vigilance states were classified as active wake, quiet wake, NREMS, and REMS (Fig. 5–6, Extended Data Fig. 9, 11-14). For sleep– wake architecture analyses, states were classified into three categories: wake (combining active and quiet wake), NREM, and REM sleep (Extended Data Fig. 2, 10).

#### Spectral analysis

Power spectral densities (PSDs) were calculated with the MATLAB function “pwelch” (hanning window of 2048 samples, with 312 samples overlap) for each state epoch and averaged across epochs. PSD of each state calculated across state epochs occurred in the first three hours of light phase in baseline day, CNO day 2 and day 5 and was normalized to the sum of power for each state.

#### Single-unit analysis

Spikes were detected as threshold crossings four standard deviations (s.d.) above the whitened data^132^. Spike waveforms (40 samples symmetrical around the peak) were semi-automatically clustered using KlustaKwik followed by manual curation using Klusters^133,134^, with spike-sorting sessions performed in 12–24 hr blocks. Clusters were defined as well-isolated single units and included in the analysis only if they fulfilled the following criteria: (1) the presence of a refractory period (less than 0.5% of inter-spike intervals < 2 ms^135^; (2) an isolation distance > 10^136^. Clusters were semi-automatically separated to regularly-spiking (RS) and fast-spiking (FS) units by inspection of their waveform, autocorrelograms, and burst index using the Cell Explorer MATLAB package^137^. For analyses of continuous recordings without state separation, neuronal spike times were converted to firing rates using 60-s bins and averaged over the relevant time windows for firing-rate analyses (Fig. 1-2, Extended Data Fig. 6, 8). For network-level analyses (mCC, dimensionality, variance, and stability index α), spike times were converted to firing rates using 100-ms bins and subsequently averaged over 15-min intervals (Fig. 3-4, Extended Data Fig. 7-9).

#### Pairwise correlations

We computed Pearson correlation coefficients between all pairs of single-unit FR trajectories within a 15 min time window in the following manner: we first omitted cells that did not fire at all within the 15 min time window, we then z-scored each unit (subtracted the time-average FR and divided by it’s temporal standard deviation). We then computed the temporal covariance matrix on this data (using the built-in Matlab function “cov”) and subtracted the diagonal (since it is all ones). A representative example correlation matrix is shown in Fig. 3a. Fig. 3b reports the average of all elements of this matrix.

#### Linear embedding dimensionality

We computed dimensionality in 15 min time windows using single-unit FR trajectories, where FRs were calculated in bins of 100 ms. We first centered the data by subtracting the mean of each unit over 15 min and remove units that had 0 spikes during this window. Principal Component Analysis was then performed on the centered activity for each window separately, where single-unit FRs are the variables and 100 ms time bins are the cases. The result was a spectrum of variance explained along each Principal Component (PC). We denote the variance along the kth PC as v_k_ such that for a network with N units there are N variance values v_1_, …, v_N_ in descending order, the total variance is given by 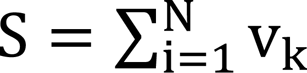, and the cumulative variance as a function of dimension is given by:

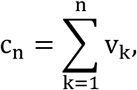

Then, dimensionality is defined as the minimal PC index for which the cumulative variance is equal to or greater than 80% of the total variance:

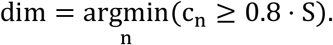

#### Dynamical stability

We computed dynamical stability indexes α for the centered FR data (such that each unit had zero mean) over the 15 min time windows using single-unit FR trajectories, where FRs were calculated in bins of 100 ms, yielding a total of T = 9000 time-points. For each time-point 1 ≤ t ≤ T within the window we average over all units to obtain the instantaneous mean FR, r(t). Then, for each t > 1 we compute the temporal difference of r(t):

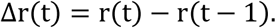

For each time window, these two quantities yield two time-matched vectors of length T − 1. The stability index α is then given by the slope of the linear fit between these two vectors, Δr and r.

#### Analysis of FR statistics per vigilance state

To assess the contribution of different brain states to upward FR homeostasis, we analyzed daily recordings during baseline, and the first, second, and last day of CNO treatment. For each neuron, all AW, NREMS, and REMS epochs lasting at least 60 s were extracted. Each epoch was normalized in length by dividing it into three equal parts, and firing rates were calculated for the first, middle, and last third. For each epoch, within-epoch change was quantified as the difference between the firing rates in the last and first third. These values were then averaged across all epochs of the same state to obtain the mean within-epoch change per neuron.

State transitions were defined as the difference between the firing rate in the first third of a given epoch and the last third of the preceding epoch. Transition values were then averaged across all relevant transitions for each neuron. Because each third corresponded to ≥20 seconds of data, these transitions reflected averaged dynamics rather than abrupt changes.

For NREM_i_–REM–NREM_i+1_ triplet analysis, only triplets containing both NREM and REM epochs with a minimum duration of 60 s per state were included. Brief awakenings of <16 s following REM were permitted, whereas spikes occurring during brief awakenings within NREM or immediately after REM were excluded from FR calculations. FR was computed for each neuron by dividing each state epoch within a triplet into three equal thirds and then averaging across all triplets.

Within-epoch firing rate changes were also quantified using a normalized measure -FR gain, calculated for each epoch as (FR_last_ – FR_first_)/ (FR_last_ + FR_first_) where FR_first_ and FR_last_ correspond to the firing rates in the first and last thirds of the epoch, and subsequently averaged across epochs for each neuron. To assess the robustness of state-dependent FR changes across analysis parameters, we also calculated within-epoch changes using constant 20-s bins at the start and end of each epoch, instead of normalizing by epoch duration. In addition, we quantified cumulative within-epoch changes by summing changes across all epochs of a given state for each neuron.

#### Patch clamp recordings and analysis in hippocampal slices

An osmotic pump delivering CNO was implanted in mouse 4–5 weeks after viral injection for chronic experiments. Mice were anesthetized with isoflurane and decapitated either 4–5 weeks after viral injection (acute experiments) or 5 days (111-113 hours) after implantation of an osmotic pump (chronic experiments). The brains were rapidly extracted and immediately sectioned into 300-µm-thick coronal slices using a vibratome (VT-1200S, Leica, Germany) in ice-cold sucrose-based cutting solution composed of (in mM): 60 NaCl, 2.5 KCl, 1.25 NaH₂PO₄, 26 NaHCO₃, 1 CaCl₂, 5 MgCl₂, 100 sucrose, and 20 glucose, continuously oxygenated with 95% O₂ and 5% CO₂. Slices were allowed to recover for 30 minutes at 34°C before being transferred to standard artificial cerebrospinal fluid (ACSF) containing (in mM): 125 NaCl, 3 KCl, 1.25 NaH₂PO₄, 26 NaHCO₃, 2.6 CaCl₂, 1.3 MgCl₂, and 15 glucose, also bubbled with 95% O₂ and 5% CO₂, and maintained at room temperature until use.

Whole-cell patch-clamp recordings were performed on hippocampal CA1 pyramidal neurons under either voltage- or current-clamp mode using an upright fluorescence microscope (SliceScope, Scientifica, UK). Expression of dTomato in the hippocampus was visually confirmed prior to recording.

Patch pipettes (3–5 MΩ) were fabricated with a horizontal puller (DMZ Universal Puller, Zeitz, Germany) and filled with intracellular solutions appropriate to the experiment. For acute experiments targeting spontaneous inhibitory postsynaptic currents (sIPSCs), the pipette solution contained^138^ (in mM): 30 K-gluconate, 110 CsCl, 0.1 CaCl₂, 10 HEPES, 4 Mg-ATP, 0.3 Na-GTP, 1.1 EGTA, and 4 QX-314. For chronic experiments, targeting both miniature excitatory postsynaptic current (mEPSC) and intrinsic excitability, the solution contained (in mM): 140 K-gluconate, 7 KCl, 5 HEPES, 0.5 MgCl₂, 5 phosphocreatine, and 0.16 EGTA.

Recordings were acquired using an amplifier (ELC-03XS, npi, Germany), digitized at 50 kHz via an A/D converter (LIH 8+8, HEKA, Germany), and recorded using custom software written in Igor Pro (WaveMetrics, USA). During experiments, slices were continuously perfused with oxygenated ACSF maintained at 33°C.

Pharmacological agents were included in the ACSF according to experimental requirements. For recordings of sIPSCs with acute CNO application ACSF was supplemented with 10 µM NBQX and 50 µM D-AP5. After a 5–10 minute baseline period, 10 µM clozapine-N-oxide (CNO) was bath-applied for 10 minutes via the perfusion system. For mEPSC recordings with chronic CNO application, ACSF contained 50 µM D-AP5, 20 µM SR95531, and 1 µM tetrodotoxin (TTX). For assessing intrinsic electrophysiological properties in chronic experiments, ACSF included 10 µM NBQX, 50 µM D-AP5, and 20 µM SR95531.

Cells were clamped at –60 mV during sIPSC recordings, at –70 mV during mEPSC recordings in voltage-clamp mode, and clamped at –65 mV in current-clamped mode for recordings assessing intrinsic properties. Liquid junction potentials were not corrected.

Events of sIPSCs and mEPSCs were detected using the Event Detection module in NeuroMatic^139^ v3.0s running on Igor Pro. Only recordings longer than 10 minutes were included in the analysis of mEPSC amplitude and frequency. Intrinsic membrane properties were subsequently analyzed using custom Python script. The resting membrane potential was recorded immediately after establishing the whole-cell configuration, under current clamp conditions without current injection. The input resistance was determined from the linier relationship between the steady-state voltage deflection and the injected current. This was accomplished using 500 ms current steps ranging from −30 to +30 pA in 10 pA increments, with the calculation performed on the stable phase (400–500 ms after current onset) of the voltage response. The action potential threshold was defined as the membrane potential at the onset of the action potential. This measurement was taken in response to a 500 ms current step at the rheobase, which represents the minimal current required to elicit a single action potential. The firing duration was measured as the time between the peak of the first and the last action potential elicited by a 500 ms current step equal to three times the rheobase current. All data were visualized using Igor Pro, GraphPad Prism, and/or custom-made Python tools.

### Behavioral assays

#### Handling

Three to four weeks after viral injection (as described above), mice were pair-housed and handled for 10 minutes daily over three consecutive days. For the next two days, they were transported via wheeled cart to the experimental floor, where they remained for 10 minutes to habituate them to the environment and transport route.

#### Fear Conditioning Paradigm

The fear conditioning (FC) apparatus consisted of a conditioning cage (25X25X36 cm) with a grid floor wired to a shock generator surrounded by an acoustic chamber. For CFC experiments, two distinct contexts were used: Context A for acquisition and Context B as a neutral, different environment. Context A was rectangular, with white light, black and white striped walls, a grid floor, and a vanilla scent. Mice were placed in Context A for 150 seconds, followed by a 20-second pure tone (2.9 kHz), after which a 2-second foot shock (0.64 mA) was delivered. This procedure was repeated twice with 45-second intervals. Thirty seconds after the third shock, the mice were returned to their home cage. For contextual FC testing, mice were returned to Context A, and freezing was measured over 5 minutes. For auditory cued FC testing, mice were placed in Context B, a round cage with red plastic walls, a white plastic floor, green light, and a rum scent. Freezing was recorded for 2.5 minutes, followed by a 2.9 kHz tone for 2.5 minutes, during which conditioned freezing was measured. Freezing behavior, defined as complete immobility, was continuously measured by an experienced experimenter blinded to treatment groups.

Mice were trained and tested as follows: On Day 1, acquisition occurred in Context A. The next day (Retrieval 0, “baseline”), contextual recall in Context A and cued testing in Context B were performed 3.5 hours apart. Two days later, an osmotic pump with CNO was implanted. Four to seven hours post-implantation, contextual recall (Retrieval 1, “CNO day 1”) and cued testing were conducted. Four days later, contextual recall (Retrieval 2, “CNO day 5”) and cued testing were repeated. Two days after pump removal, mice underwent a final contextual recall (Retrieval 3, “Washout”) and cued test, 3.5 hours apart.

Dlx-Gq and Dlx-Cnt groups were divided into two subgroups: one underwent retrieval 1 + 2 (at CNO day 1 and day 5) where the second underwent only retrieval 2 at CNO day 5 (Extended Data Fig. 15).

#### Histology and confocal imaging

At the end of the experiments, mice were transcardially perfused with phosphate-buffered saline (PBS), followed by cold 4% paraformaldehyde (PFA). Brains were removed and post-fixed in 4% PFA for 24 h. 60-µm coronal sections were prepared using a vibrating microtome (Leica VT1200). Free-floating sections were washed in PBS, blocked for 2 h in PBS containing 1% bovine serum albumin (BSA) and 0.3% Triton X-100, and then incubated for 2 days at 4 °C with mouse anti-GAD67 primary antibody (1:200, Sigma-Aldrich, MAB5406). Sections were then washed in PBS and incubated for 2 h at room temperature with Alexa Fluor 488 goat anti-mouse secondary antibody (1:500, Abcam, ab150113) in PBS containing 1% BSA. Finally, sections were washed in PBS and mounted on slides with DAPI-containing mounting medium (Sigma-Aldrich, MBD0020).

Confocal fluorescence images were acquired using an FV1000 confocal microscope (Olympus, Japan) with 20× and 60× objectives (the latter imaged through a water droplet). Z-stack images were collected with slice intervals of 4 µm (×20) or 3 µm (×60). Maximum-intensity projections were generated from the z-stacks using the ImageJ program.

### Statistical analysis

Unless otherwise specified, data analysis was conducted using custom-written code in MATLAB (MathWorks, Natick, MA). Statistical analyses were performed using MATLAB (MathWorks, Natick, MA) and Graph-Pad Prism 10.2.2 software. Linear mixed effects models were performed to assess the influence of experimental days and behavioral state, on the dependent variable(s) while accounting for variability across animals and neurons. Analyses were conducted using the “fitlme’ function in MATLAB (MathWorks, Natick, MA). The models formulas are described in the supplementary information tables S1-4. For each contrast the model returned the estimated difference in response variable (Estimate), its standard error (SE), the mean ratio and p-values. P-values were adjusted using Bonferroni correction. Data are presented as mean ± standard error of the mean (SEM) unless stated otherwise. Statistical significance markers are: ns, not significant, \**P* < 0.05, \*\**P* < 0.01, \*\*\**P* < 0.001, \*\*\*\**P* < 0.0001. Sample sizes, *P*-values, and statistical tests for each experiment are provided in the respective figure legends. All experiments were performed with at least four mice.

## Data availability

The authors declare that the main data supporting the findings of this study are available within the article and its Supplementary Information files. Datasets generated during this study and any additional information required to reanalyze the datasets will be available upon request from the lead contact.

## Code availability

All original code will be deposited at Zenodo and publicly available as of the date of publication.

## Acknowledgments

We thank Yuval Nir, Shimon Marom, Inbal Goshen, Dori Derdikman, Noam Ziv, Boaz Styr, Naama Brenner, Omri Barak, and Michael London for their thoughtful discussion and comments on an early version of this manuscript. This work was funded by research grants awarded to I. Slutsky from the European Research Council (grants 724866 and 101097788), the Israel Science Foundation ISF, 1663/18 and 2634/24), the Deutsche Forschungsgemeinschaft (DFG, grant 448865644 with A.A.), the BIRAX (The Britain-Israel Research and Academic Exchange) and The Parasol Foundation Trust, and the Rosetrees Trust (grant CF-2022-2\129). S. Remy was supported by the Deutsche Forschungsgemeinschaft (DFG CRC 1436, DFG RTG 2413 SynAGE). R.D. was supported by the Israel Science Foundation (ISF, 2097/24 and 2994/24) and the Alon scholarship for the integration of outstanding faculty. R.A. is grateful to the Sagol School of Neuroscience for a PhD fellowship; K.L.B.O. is grateful to the Sagol School of Neuroscience for an MSc fellowship and to the Sieratzki Institute for Advances in Neuroscience for a PhD fellowship; L.S. acknowledges the Center for the Physics of Biological Function at Princeton University for an independent postdoctoral fellowship; B.M. was funded by the DFG (grant 672192 to A.A. with I.S.). This work was conducted in partial fulfillment of the requirements for the PhD degree by R.A. at the Gray Faculty of Medical and Health Sciences and the Sagol School of Neuroscience, Tel Aviv University.

## Author contributions

Conceptualization: IS, RA, KLBO; Methodology: RA, LRH, KLBO, OS, LS, LSo, AA, SR; Experimentation: RA, KLBO, LSo, HK, AZ, DM, AR, BM; Formal analysis: RA, KLBO, LS, LSo, HK, DM, AR, LRH; Modeling: LDM, LS, RD; Funding acquisition and supervision: IS, SR, RD, AA; Writing – all the authors.

## Supplementary Information

### Extended Data

**Extended Data Fig. 1.**
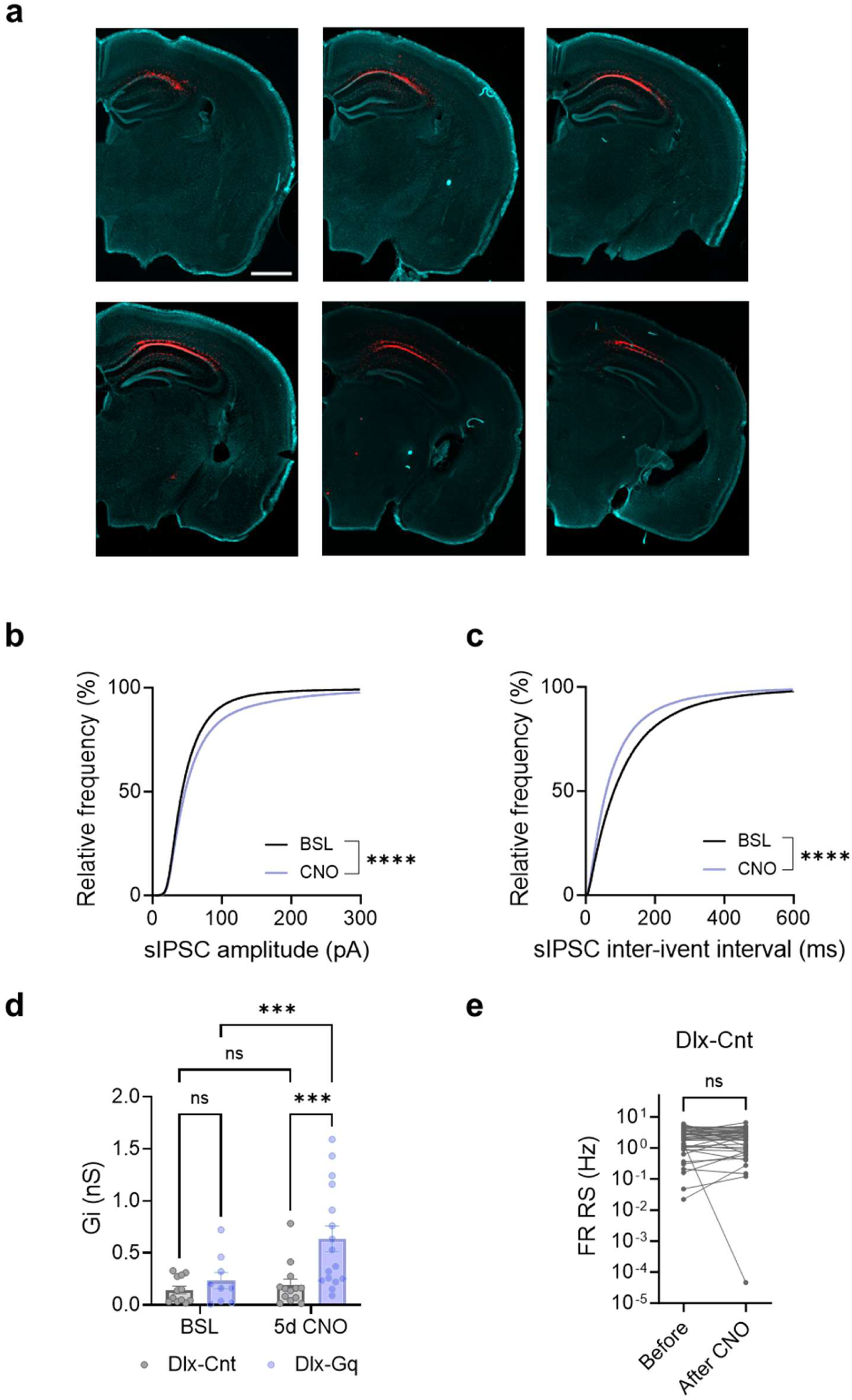
Effect of hM3Dq activation in interneurons on sIPSCs in excitatory neurons (related to Fig. 1). **a,** Representative low-magnification histological sections of hippocampal CA1 showing AAV-hDlx-hM3D(Gq)-2A-NLS-dTomato expression (red) in interneurons and DAPI nuclear staining (cyan), illustrating viral spread along the anterior-posterior axis. Six coronal sections are shown (section thickness: 120 µm; inter-section interval: 240 µm). Scale bar, 1 mm. **b-c,** Cumulative probability plots of amplitude (a) and inter-event interval (IEI, b) of sIPSCs under baseline (BSL, black) and CNO (purple) conditions. BSL data represent a 5-minute window before CNO application, while CNO data represent the final 5 minutes of the 10-minute CNO application period (*n* = 31 neurons, 5 mice, BSL Vs CNO : sIPSC amplitude \*\*\*\**P* < 0.0001; sIPSC IEI \*\*\*\**P <* 0.0001). **d,** Integrated inhibitory conductance 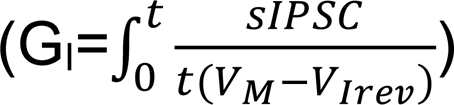 increased after 5 days of 10 µM CNO incubation in Dlx-Gq cultures (\*\*\**P* = 0.0003) but not in Dlx-Cnt hippocampal cultures (*P* = 0.76). **e**, FR of RS units (*n* = 39 units, 3 mice, Dlx-Cnt group) before and 6 hr after CNO delivery via osmotic pump, did not show a significant reduction (*P* = 0.33). Kolmogorov-Smirnov test (b and c); mixed-effect model with uncorrected Fisher’s LSD (d); Wilcoxon matched-pairs signed rank test (e).

**Extended Data Fig. 2.**
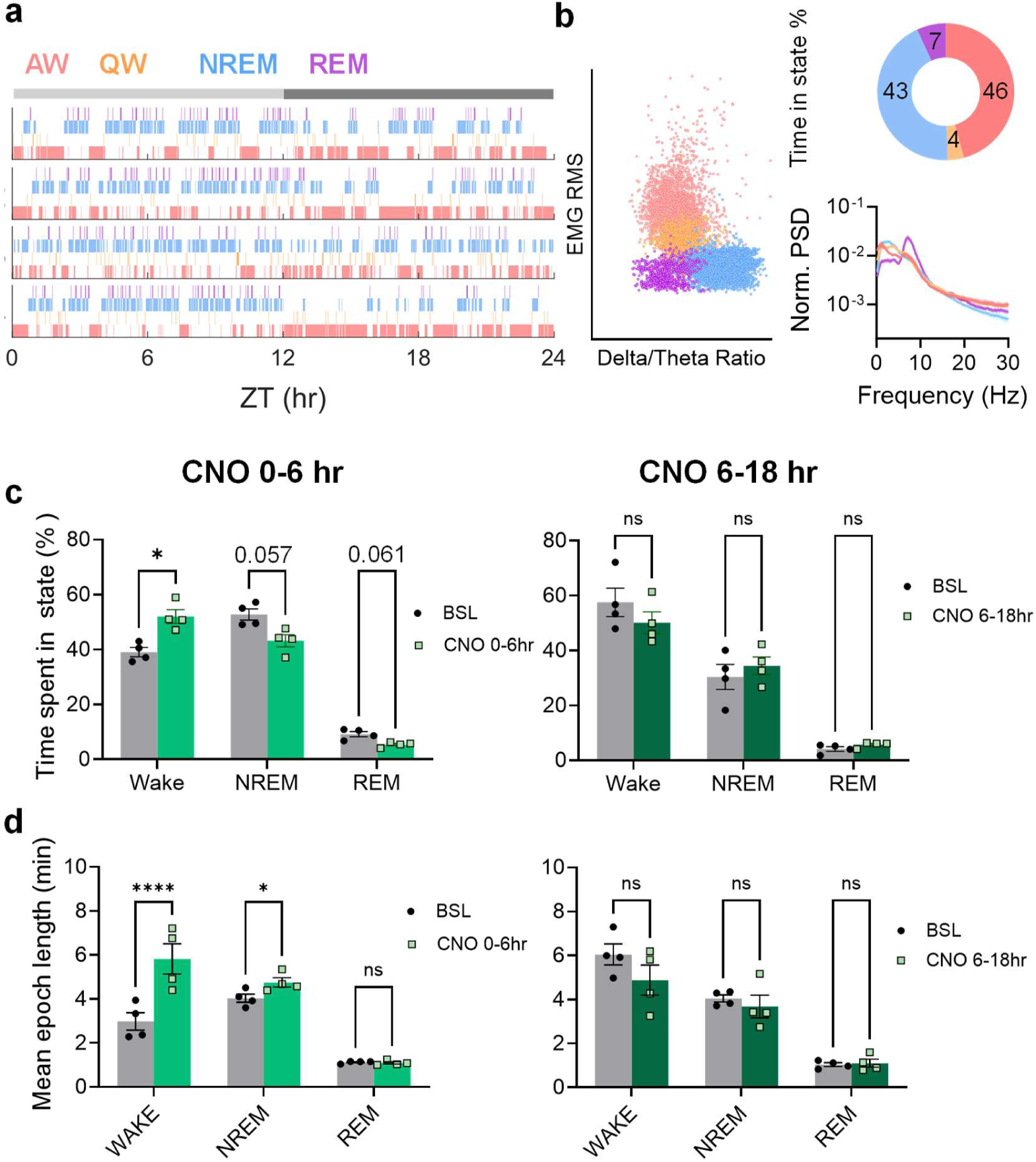
Sleep–wake cycle is transiently affected during the first 6 hr after osmotic pump implantation in Dlx-Cnt mice (related to Fig. 1). **a**, Individual hypnograms of Dlx-Cnt mice across a 24-hr undisturbed baseline. Vigilance states are color-coded as active wake (AW, pink), quiet wake (QW, orange), NREM sleep (blue), and REM sleep (purple). Light and dark phases are indicated by the grey bar above. **b**, Left : scatter plot of vigilance state separation based on EMG RMS and delta/theta ratio in 1-s windows. Right :proportion of time spent in vigilance states during 24 hr of baseline (top) and LFP spectral power normalized to total power (mean ± s.e.m.) across vigilance states (bottom). **c**, Percentage of time spent in WAKE, NREM, and REM across baseline (BSL, grey) and first six hours (left, light green) and 6-18hr (right, dark green) after CNO pump implantaion. (WAKE — BSL vs. CNO 0-6 hr, \**P* = 0.021; CNO 6-18 hr, *P* = 0.66; NREM — BSL vs. CNO 0-6 hr, *P* = 0.057; CNO 6-18 hr, *P* = 0.88; REM — BSL vs. CNO 0-6 hr, *P* = 0.061; CNO 6-18 hr, *P* = 0.45). **d**, Mean epoch length of WAKE, NREM, and REM across baseline (BSL, grey) and first six hours (left, light green) and 6-18hr (right, dark green) after CNO pump implantaion. (WAKE — BSL vs. CNO 0-6 hr, \*\*\*\**P* < 0.0001; CNO 6-18 hr, *P* = 0.20; NREM — BSL vs. CNO 0-6 hr, \**P* = 0.041; CNO 6-18 hr, *P* = 0.90; REM — BSL vs. CNO 0-6 hr, *P* = 0.99; CNO 6-18 hr, *P* = 0.99). Two-way ANOVA with Sidak’s multiple comparisons tests (c,d).

**Extended Data Fig. 3.**
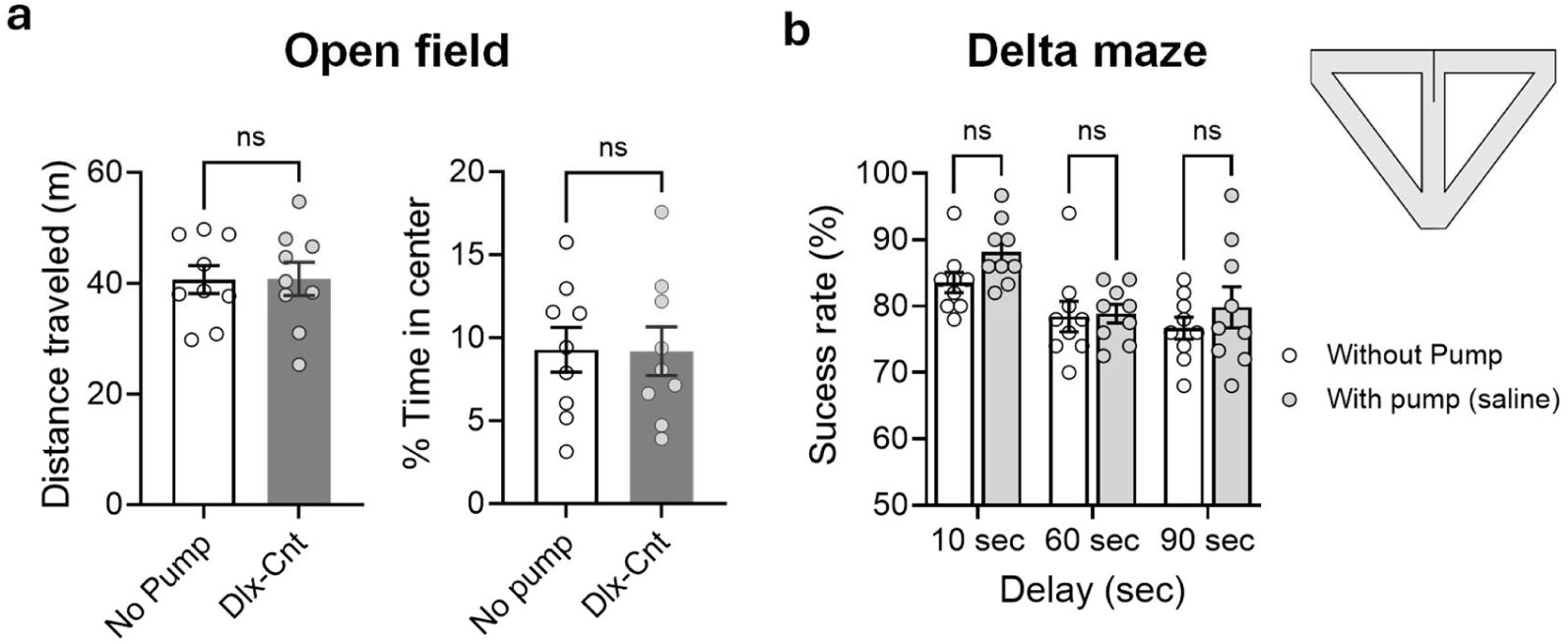
Osmotic pump implantation does not affect locomotor activity or spatial working memory (related to Fig. 1). **a,** Open-field performance in Dlx-Cnt mice implanted with CNO-filled osmotic pumps (tested 4–7 h post-implantation) compared with non-implanted controls (*n* = 9 per group). Distance traveled and percentage of time spent in the center of the arena were not significantly different (*P* > 0.9 for both). **b**, Spatial working memory in Dlx-Gq mice assessed on a separate day before implantation and 4–7 h after implantation of a saline-filled osmotic pump (*n* = 9). Performance across 10-, 60-, and 90-s delays did not differ significantly between conditions (10 s: *P* = 0.06; 60 s: *P* = 0.85; 90 s: *P* = 0.19). Mann–Whitney test (a); repeated-measures two-way ANOVA followed by Tukey’s test (b).

**Extended Data Fig. 4.**
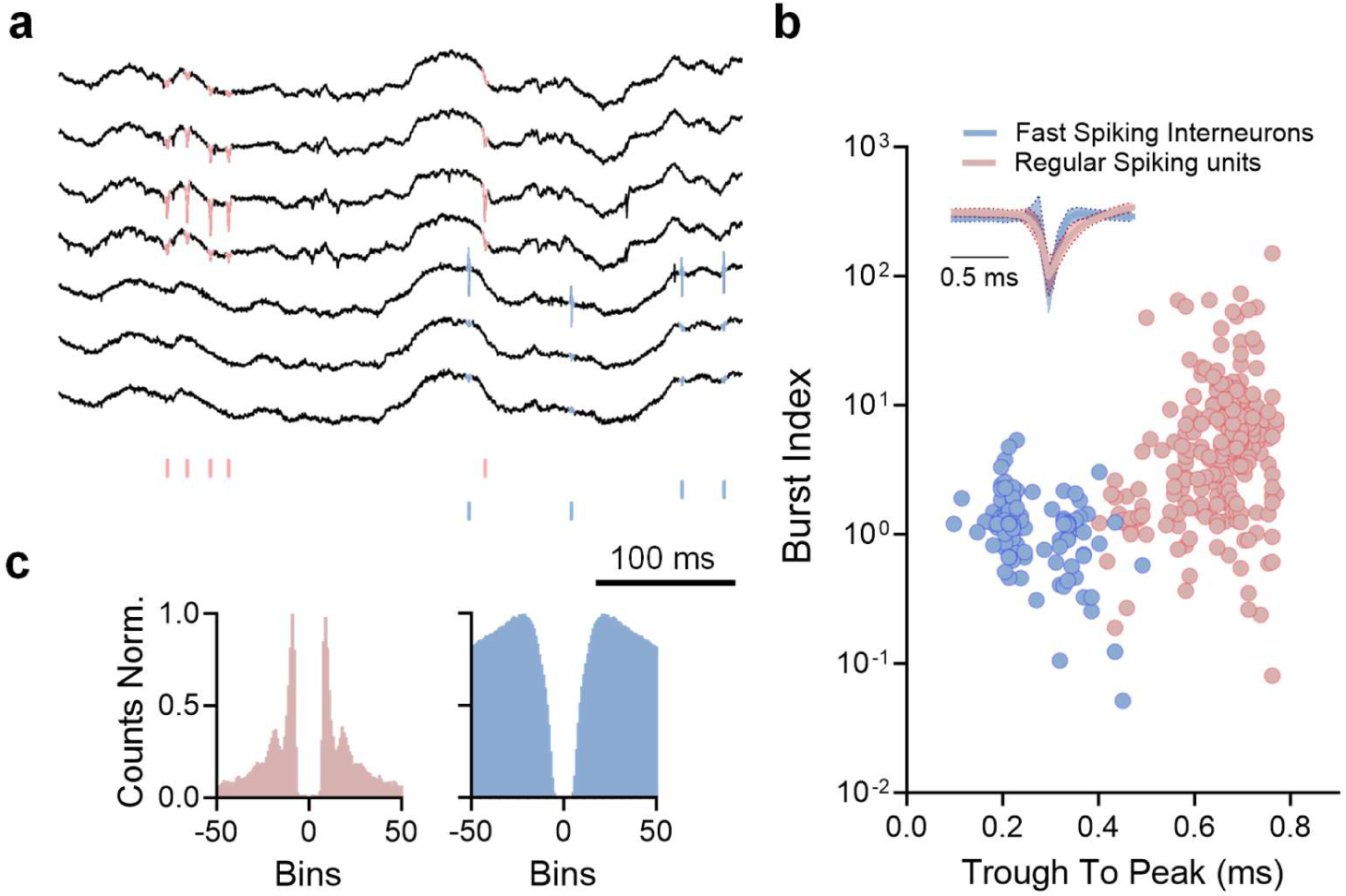
Cell type classification from extracellular recording in CA1 hippocampus (related to Fig. 1). **a**, Raw data traces of extracellular recordings from two tetrodes, spikes are classified as RS neurons (pink) or FS interneurons (blue). Scale bar: 100 ms. **b**, Scatter plot displaying separation of RS and FS units based on waveform trough-to-peak duration and burst index. *Inset*: Average z-scored waveform for each group, shaded area - SD. **c**, Representative auto-correlogram of an RS (*left*) and an FS (*right*) unit.

**Extended Data Fig. 5.**
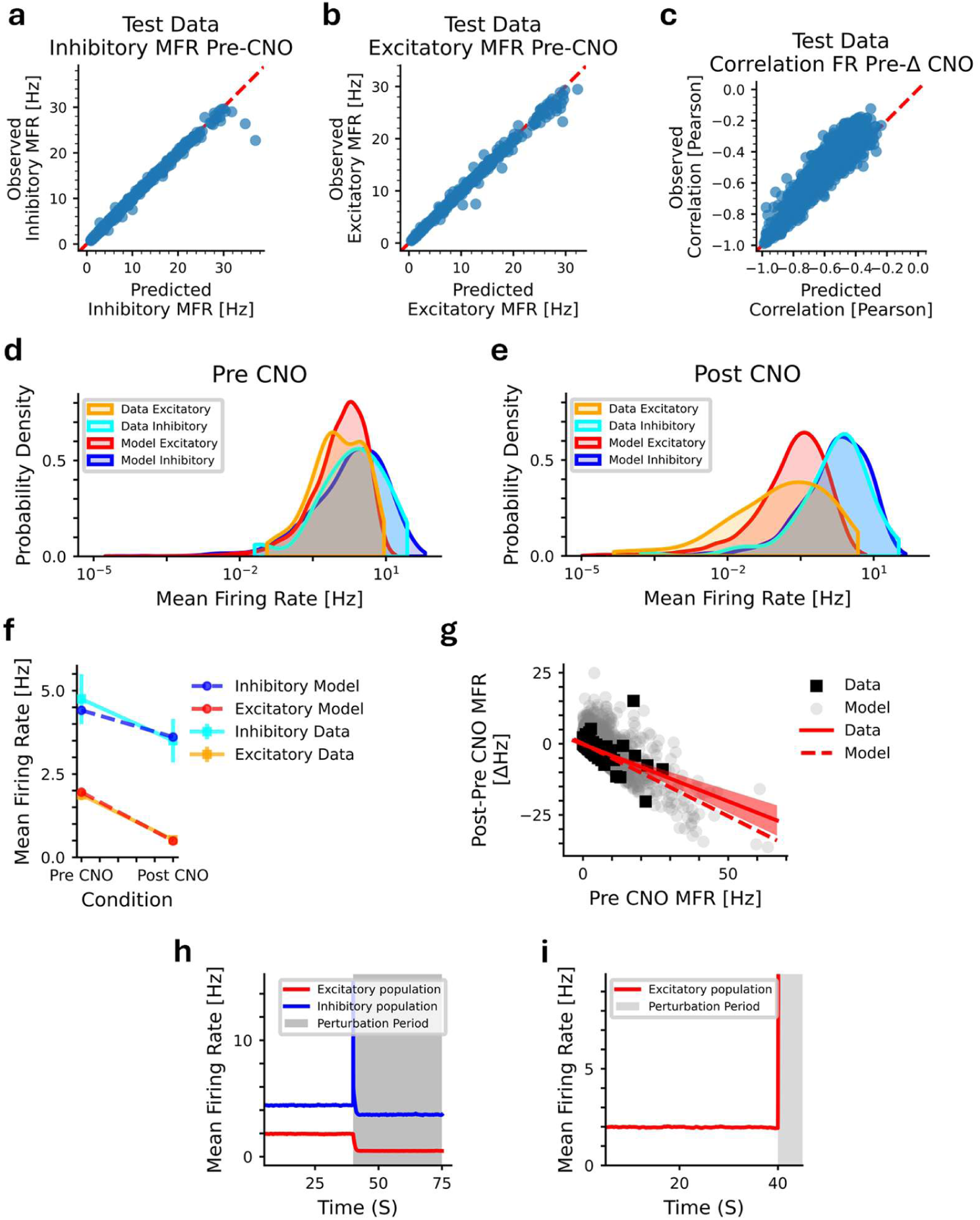
Paradoxical effect explained by inhibitory-stabilized networks (ISN, related to Fig. 1). **a-c,** Simulation-based Inference reproduces key statistics on a simulated test dataset: (a) inhibitory MFR (R²=0.98), (b) excitatory MFR (R²=0.97), (c) correlation of all cells between baseline firing rates and CNO-induced changes (see g) (R² = 0.54). **d**, Distribution of neuronal FRs on CNO day 1 for data (N = 120 excitatory, 66 inhibitory) and fitted model (N = 7500 excitatory, 2500 inhibitory), before CNO application. **e**, Corresponding post CNO distributions. **f**, Comparison between population MFR data (SEM error bars) and model (same as in e). **g**, Correlations between neuronal FRs before CNO (pre CNO) and the difference in FRs after CNO pump implementation (Post-Pre CNO). Shaded area: 95% CI of data correlation (r=−0.40, 95% CI [−0.51, −0.27]); model r = −0.59. **h**, Time course of population MFRs in the model. Note that, as expected, the inhibitory MFR transiently increases before stabilizing below baseline^1^. **i**, Excitatory MFR with and without inhibitory synapses. Paradoxical effects in I/E networks require an inhibitory-stabilized regime^2^; removing inhibition leads to excitatory divergence. Same model used in d-i.

**Extended Data Fig. 6.**
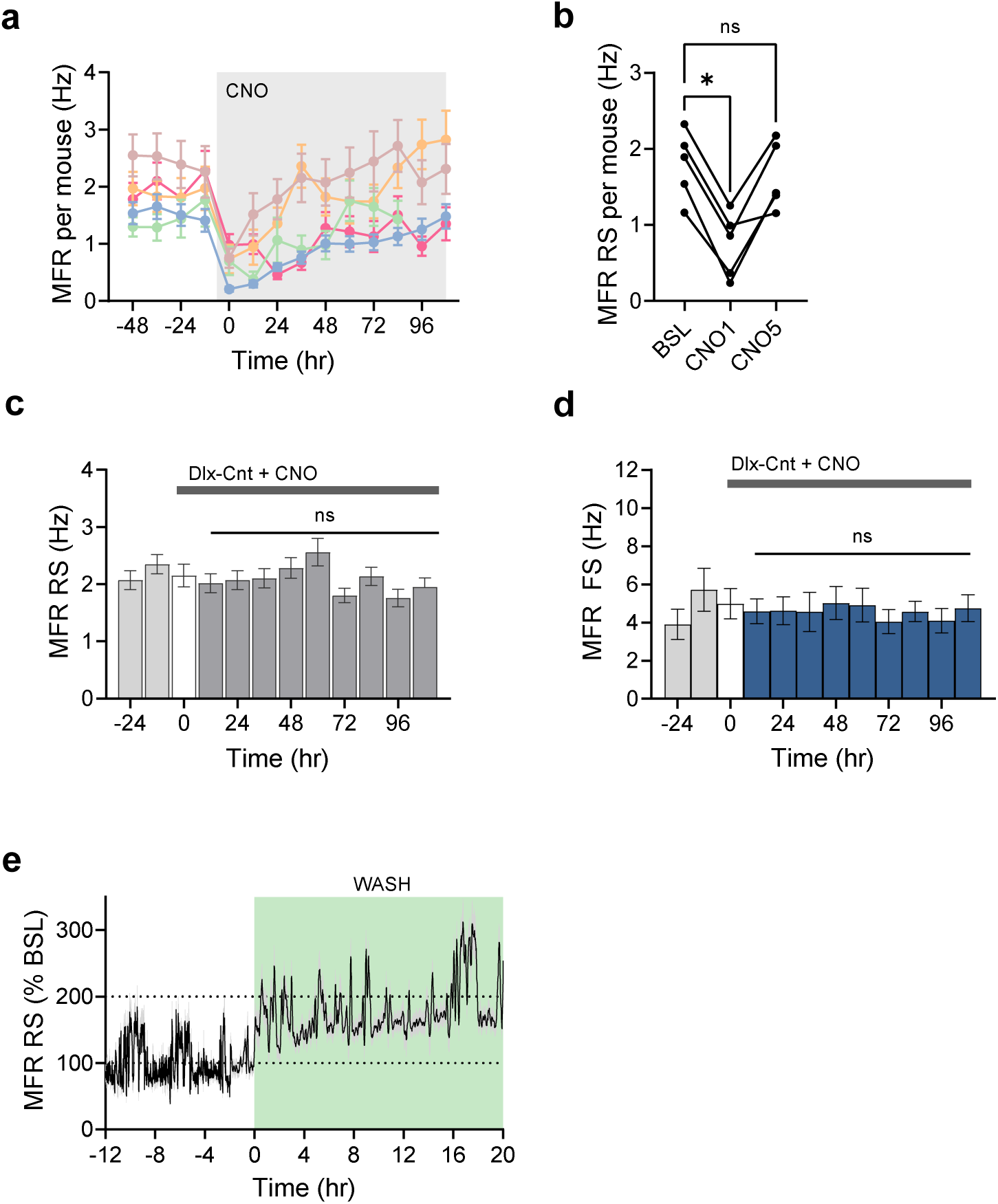
MFR of CA1 RS and FS neurons following hM3Dq INs activation (related to Fig. 2). **a,** Per-mouse MFR dynamics across the experiment (*n* = 5 mice). **b**, Per-mouse MFR statistics (the same data as in a). **c**, MFR of RS units recorded from Dlx-Cnt mice (18.8 ± 6.4 units per mouse per day; 97.6 ± 12.5 units per recording day across 4 mice; 586 units in total for 6 recording days) during baseline (BSL, light grey) and after CNO administration via osmotic pump (dark grey), first 6 hr post-CNO (white) were excluded from analysis. No significant changes in MFR were observed (0–108 hr: *P* ≥ 0.435). **d**, Same as in (c), FS neurons recorded from four Dlx-Cnt mice (10.7 ± 3.6 units per mouse per day; 59.8 ± 5.1 units per recording day across 4 mice; 359 units in total for 6 recording days). MFR did not change following CNO administration (0–108 hr: *P* > 0.999). **e**, Representative trace of normalized MFR (as a percentage of baseline, hours: −12 - 0) of RS units recorded from one mouse (32 ± 1 units) during the 12 hours preceding the removal of the osmotic pump and the 20 hours following removal. One-way ANOVA Friedman test (b); linear mixed effects regression (LMER) (c-d), For pairwise statistical contrasts, see Table S1g-h).

**Extended Data Fig. 7.**
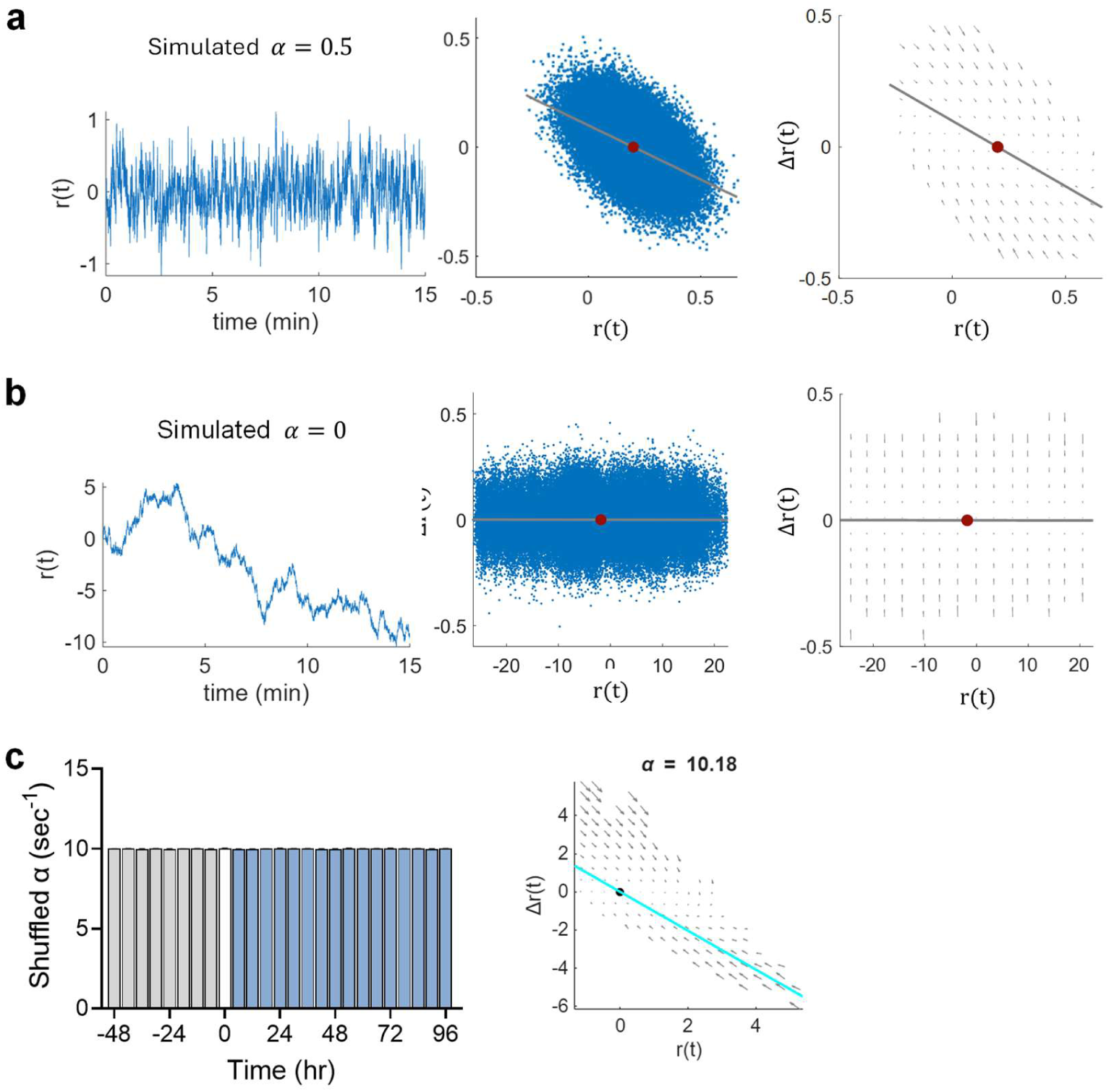
Estimating the stability index *α* on simulated dynamics (related to Fig. 3). **a,** Simulated trajectories of the dynamics Δ*r*(*t*) = −α ⋅ *r*(*t*) + *η*(*t*) with α = 0.5 and *η*(*t*) white noise with 0 mean and variance 1. Left: example trace of the population mean activity *r*(*t*). Middle: relationship between *r*(*t*) and its temporal derivative Δ*r*(*t*), showing a clear negative slope (of 0.5) consistent with stable dynamics. Right: corresponding flow field representation. **b**, Simulations with no stability (α = 0). Left: activity trace exhibits random-walk behavior. Middle: Δ*r*(*t*) shows no systematic relationship with *r*(*t*). Right: flow field is flat, indicating absence of restoring dynamics. **c**, Shuffled control analysis. Left: α values estimated from temporally shuffled data across all mice and recording days, binned into 6h windows. No systematic change was observed across baseline or CNO time points (*P* > 0.23 for all comparison). Right: example Δ*r*(*t*) versus *r*(*t*) scatter for shuffled data, showing an artificially inflated α estimate that does not capture the underlying temporal structure. Linear mixed effects regression (LMER; c), For pairwise statistical contrasts, see Table S2e).

**Extended Data Fig. 8.**
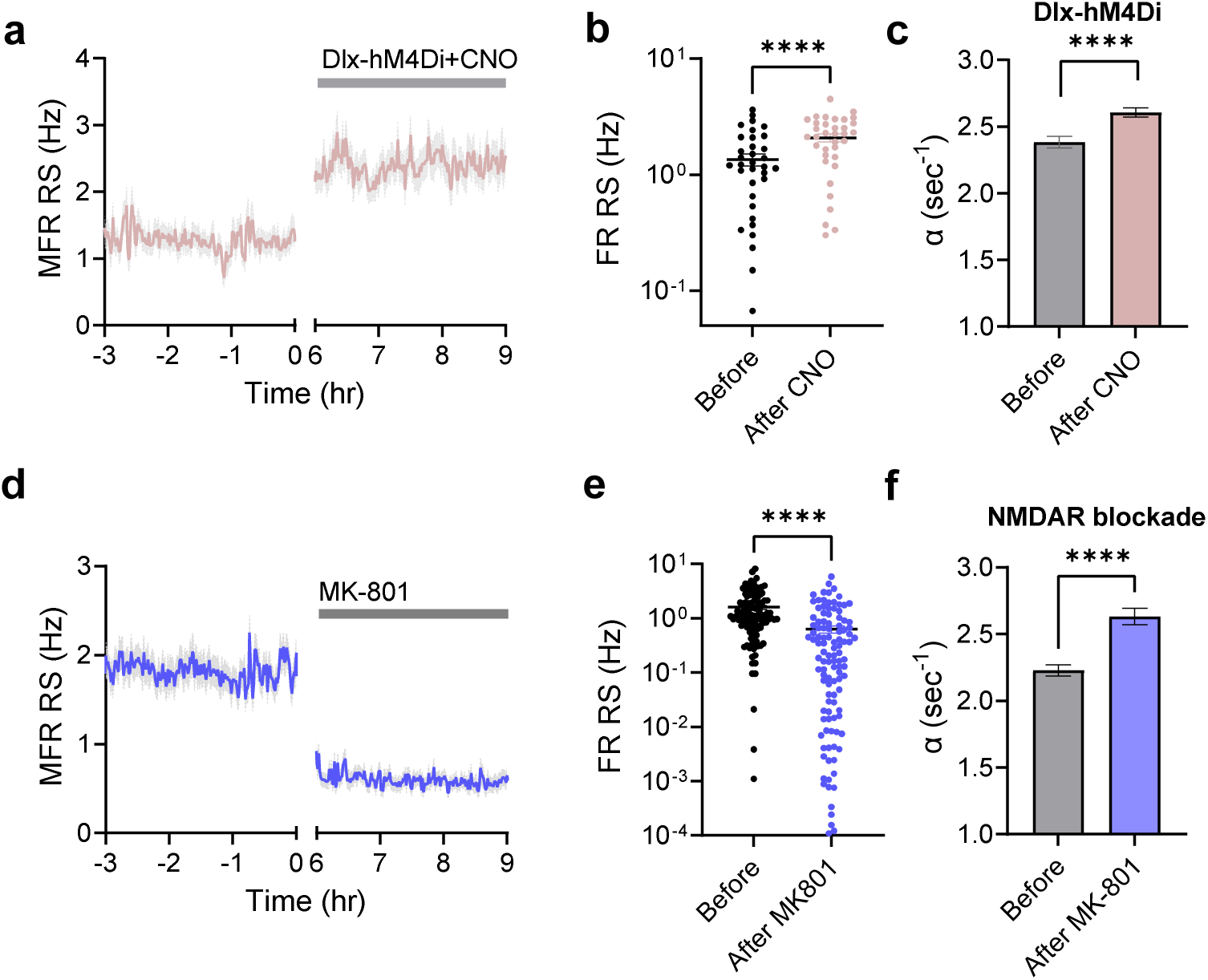
Modulation of stability index *α* (related to Fig. 3). **a-c**, Effect of CNO delivered via osmotic pump on MFR of RS units (a), single-unit FR (b, \*\*\*\**P* < 0.0001), and dynamical stability α (c, estimate 0.22 ± 0.04, mean ratio = 0.91, \*\*\*\**P* = 1.6228e-06) in behaving mice expressing Dlx-hMDi in the hippocampus (33 RS units, 3 mice). **d-f**, Effect of MK-801 NMDAR blocker delivered to the CA1 via osmotic pump at concentration of 1.25 mM on MFR of RS units (d), single-unit FR (E, \*\*\*\**P* < 0.0001), and dynamical stability α (F, estimate 0.40 ± 0.05, mean ratio = 0.85, \*\*\*\**P* = 6.5328e-13) in behaving mice (115 RS units, 4 mice). Analysis of data from Ruggiero et al^3^. Wilcoxon test (b,e), Linear mixed effects regression (LMER; c,f see Table S2g-h).

**Extended Data Fig. 9.**
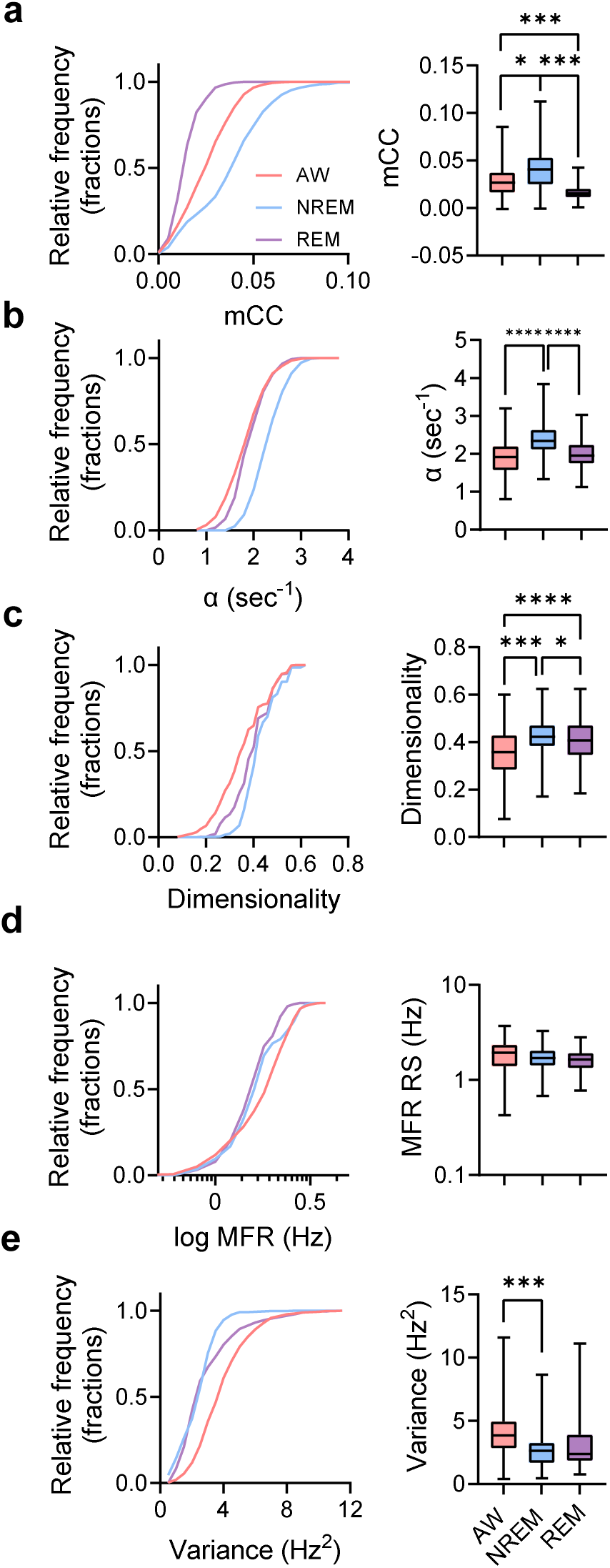
State dependency of network parameters under baseline conditions (related to Fig. 4). **a**, Mean correlation coefficient (mCC) across putative RS neurons during active wake (AW), NREM sleep, and REM sleep. Left: cumulative distributions show the highest synchrony in NREM. (\*\*\*\**P <* 0.0001 for all comparison). Right: group summary (boxplots) confirms significantly elevated mCC in sleep states (AW vs NREM: \**P =* 0.03; AW vs REM: *\*\*\*P =* 0.0005; NREM vs REM: *\*\*\*P =* 0.0004). **b**, Dynamical stability (α) differs across states. Left: cumulative distributions demonstrate stronger stability in NREM relative to AW and REM (\*\*\*\**P <* 0.0001 for all comparison). Right: boxplots reveal significantly higher α during NREM (AW vs NREM: \*\*\*\**P <* 0.0001; AW vs REM: *P =* 0.41; NREM vs REM: *\*\*\*\*P <* 0.0001). **c**, Dimensionality of population activity. Left: distributions show higher dimensionality in NREM and REM in comparison to AW (\*\*\*\**P <* 0.0001 for all comparison). Right: summary confirms significantly higher dimensionality in NREM relative to AW and REM (AW vs NREM: \*\*\**P =* 0.0005; AW vs REM: *\*\*\*\*P <* 0.0001; NREM vs REM: *\*P =* 0.02). **d**, Mean firing rate of RS neurons. Left: cumulative distributions of log-transformed firing rates show state dependence (\*\*\*\**P <* 0.0001 for all comparison). Right: summary statistics confirm no significant differences across states (*P* > 0.12 for all comparison). **e**, Variance of population activity. Left: cumulative distributions show reduced variance during NREM relative to AW (\*\*\*\**P <* 0.0001 for all comparison). Right: boxplots confirm significantly decreased variance in NREM (AW vs NREM: \*\*\**P =* 0.0002; AW vs REM: *P =* 0.17; NREM vs REM: *P =* 0.38). Kolmogorov–Smirnov tests with Bonferroni correction (a-e, left panel); Linear mixed effects regression (LMER; a-e, right panel), For pairwise statistical contrasts, see Table S3g-k.

**Extended Data Fig. 10.**
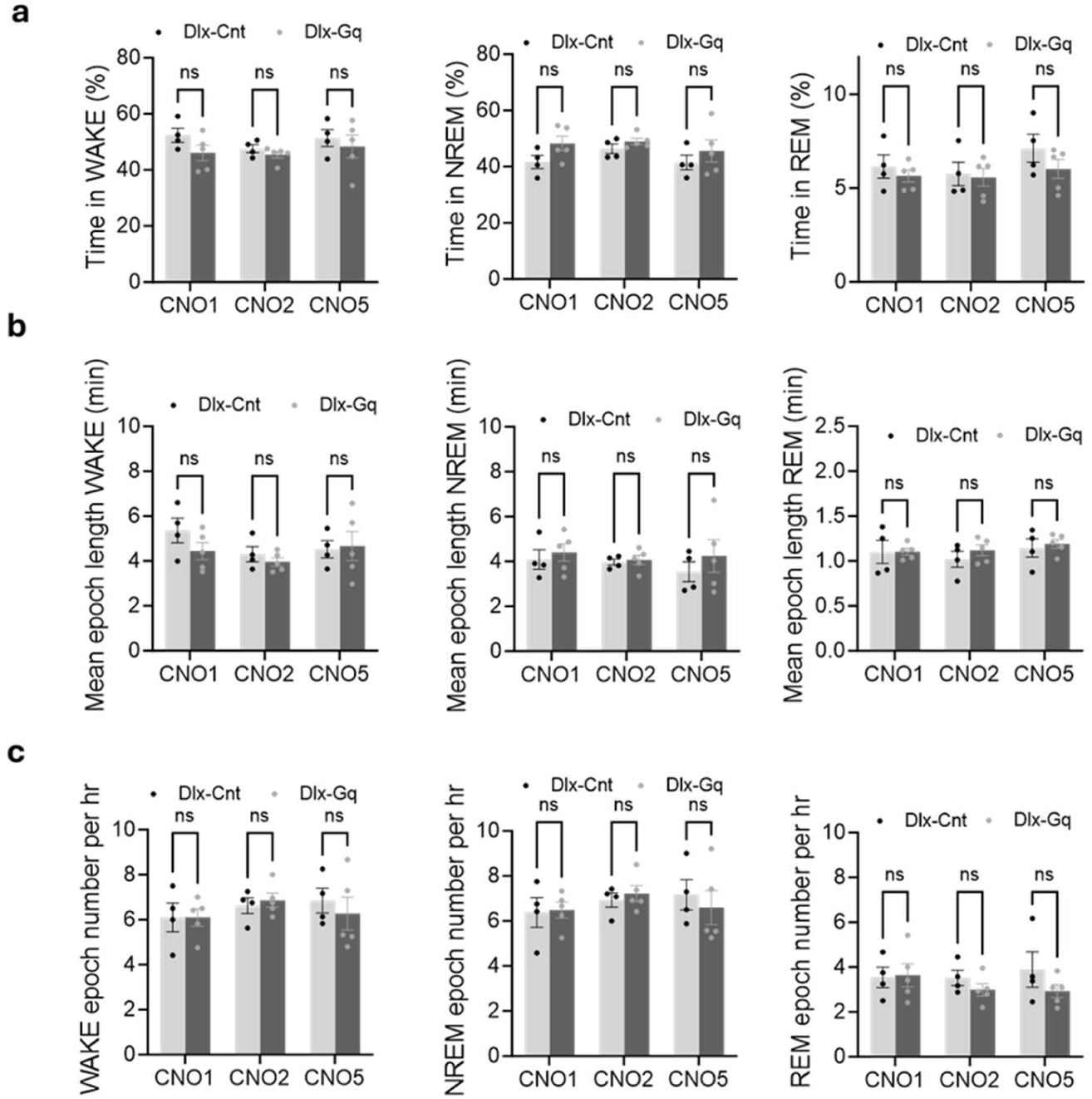
Sleep–wake architecture is preserved following chronic CNO in Dlx-Gq mice (related to Fig. 5). **a,** Percentage of time spent in WAKE (left; active + quiet wake), NREM (middle), and REM (right) states in Dlx-Cnt (n=4) and Dlx-Gq (n=5) mice on days 1, 2, and 5 of CNO delivery via osmotic pump. No significant effects of group or day were observed (WAKE *P* > 0.13; NREM *P* > 0.10; REM *P* > 0.22 for all comparison). **b,** Mean epoch duration for WAKE, NREM, and REM across the same days and groups. No significant effects of group or day were detected (WAKE *P* > 0.21; NREM *P* > 0.43; REM *P* > 0.07 for all comparison).**c,** Number of epochs per hour for WAKE, NREM, and REM across the same days and groups. No significant effects of group or day were found (WAKE *P* > 0.41; NREM *P* > 0.46; REM *P* > 0.16 for all comparison). Two-way ANOVA with Tukey multiple comparison test (a-c), Data are presented as mean ± s.e.m across mice.

**Extended Data Fig. 11.**
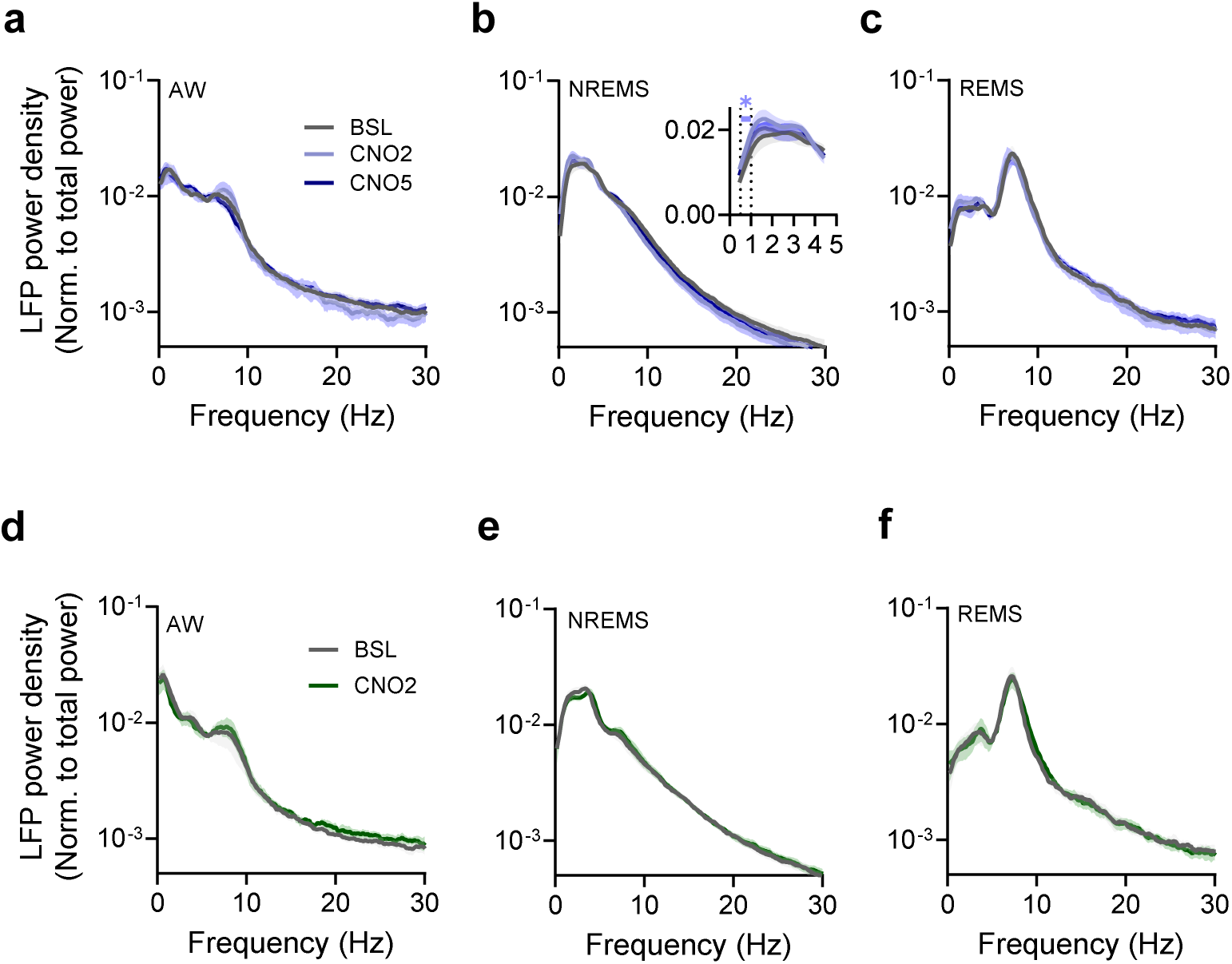
Power spectral densities in the hippocampus are largely unaffected by chronic hM3Dq activation in INs (related to Fig. 5). **a-c,** LFP power profile (mean ± SEM.) in Dlx-Gq group (n = 5) normalized to total power across different states: AW (a), NREMS (b) and REMS (c) at BSL, CNO2 and CNO5. No significant differences were observed across state frequency bands, except for a decrease in 8 −9 Hz theta band in CNO5 (\**P* < 0.04) and an increase in SWA (0.1-1 Hz) during NREMS at CNO2 (\**P* < 0.04). **d-f**, LFP power profile in Dlx-Cnt group (*n* = 4) normalized to total power across different states: AW (d), NREMS (e) and REMS (f) at BSL and CNO2. No significant differences were observed between days. *P* > 0.23 for all comparisons. Two-way ANOVA with Dunnett’s multiple comparisons (a-f).

**Extended Data Fig. 12.**
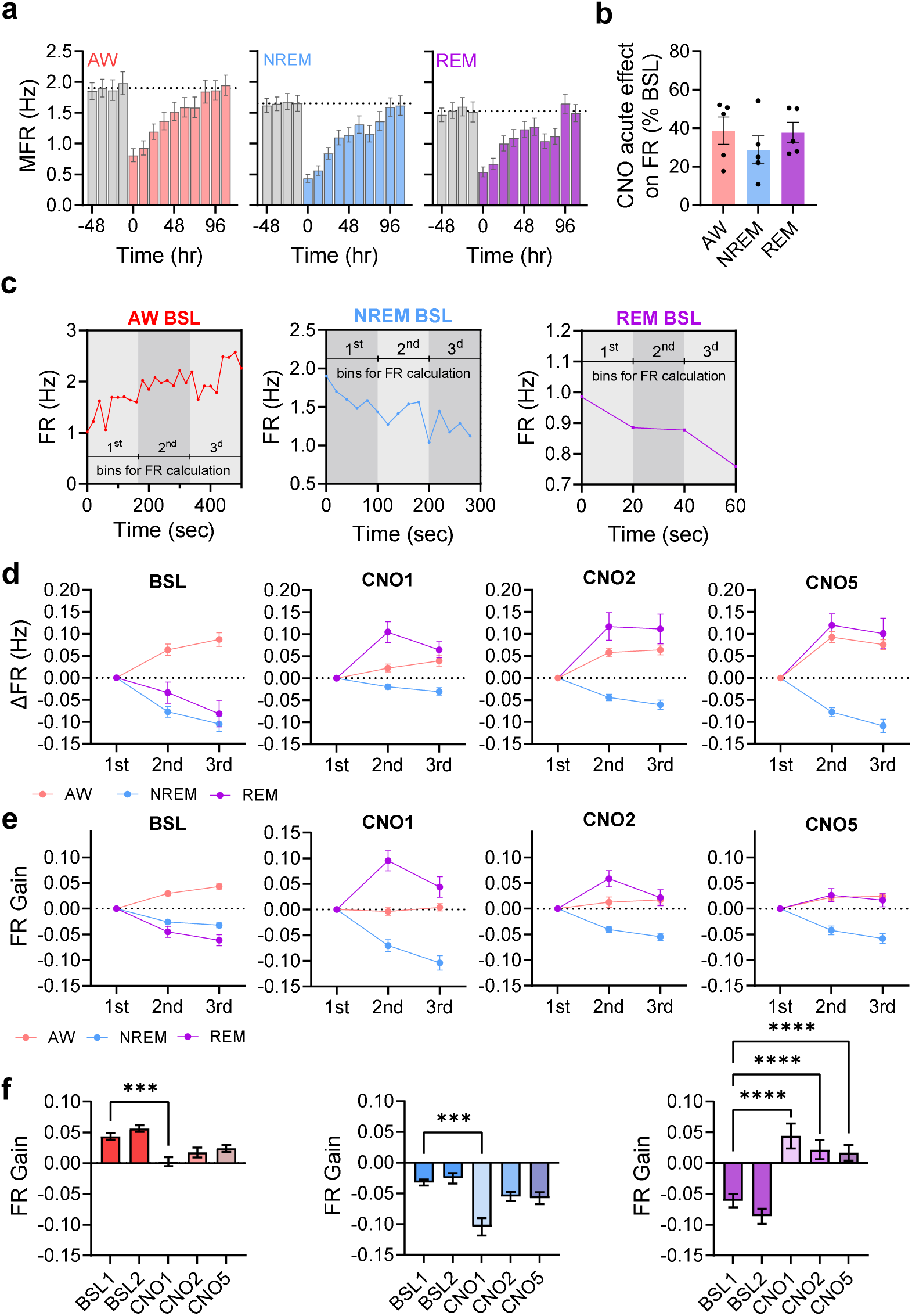
State-dependent firing rate changes across states (related to Fig. 5). **a-b**, MFR of RS units in the Dlx-Gq group across 12-hr blocks during BSL (gray) and CNO exposure in AW (left), NREMS (middle) and REMS (right). RS MFR decreased during 0–48 hr post-CNO (\**P* < 0.05 AW, REM; \*\**P* < 0.01 NREM) but recovered by 60–108 hr (*P* > 0.2 for all states) across all states (Kruskal-Wallis followed by Dunn’s multiple comparisons test). **c**, Example firing rate dynamics within representative AW, NREMS, and REMS epochs from one mouse, plotted in 20-s bins across all recorded neurons, illustrating typical within-epoch changes. **d**, Within-epoch FR changes across days (BSL, CNO1, CNO5). FR is plotted across epoch thirds after subtracting the first third. **e**, Within-epoch FR gain across days, plotted across epoch thirds after normalization to epoch firing rate. **f**, Average within-epoch FR gain for AW, NREMS, and REMS. In AW, FR gain was significantly reduced on CNO1 (CNO1 \*\*\**P* = 0.0009; CNO2 *P* = 0.2; CNO5 *P* = 0.9; Kruskal–Wallis test followed by Dunn’s multiple comparisons). In NREMS, FR gain was significantly decreased on CNO1 (CNO1 \*\*\**P* = 0.0003; CNO2 *P* = 0.7; CNO5 *P* = 0.1). In REMS, FR gain shifted from within-epoch decreases at baseline to increases across all CNO days (CNO1–5: \*\*\*\**P* < 0.0001). Data are presented as mean ± s.e.m across neurons; ns, not significant; \**P* < 0.05, \*\**P* < 0.01, \*\*\**P* < 0.001, \*\*\*\**P* < 0.0001.

**Extended Data Fig. 13.**
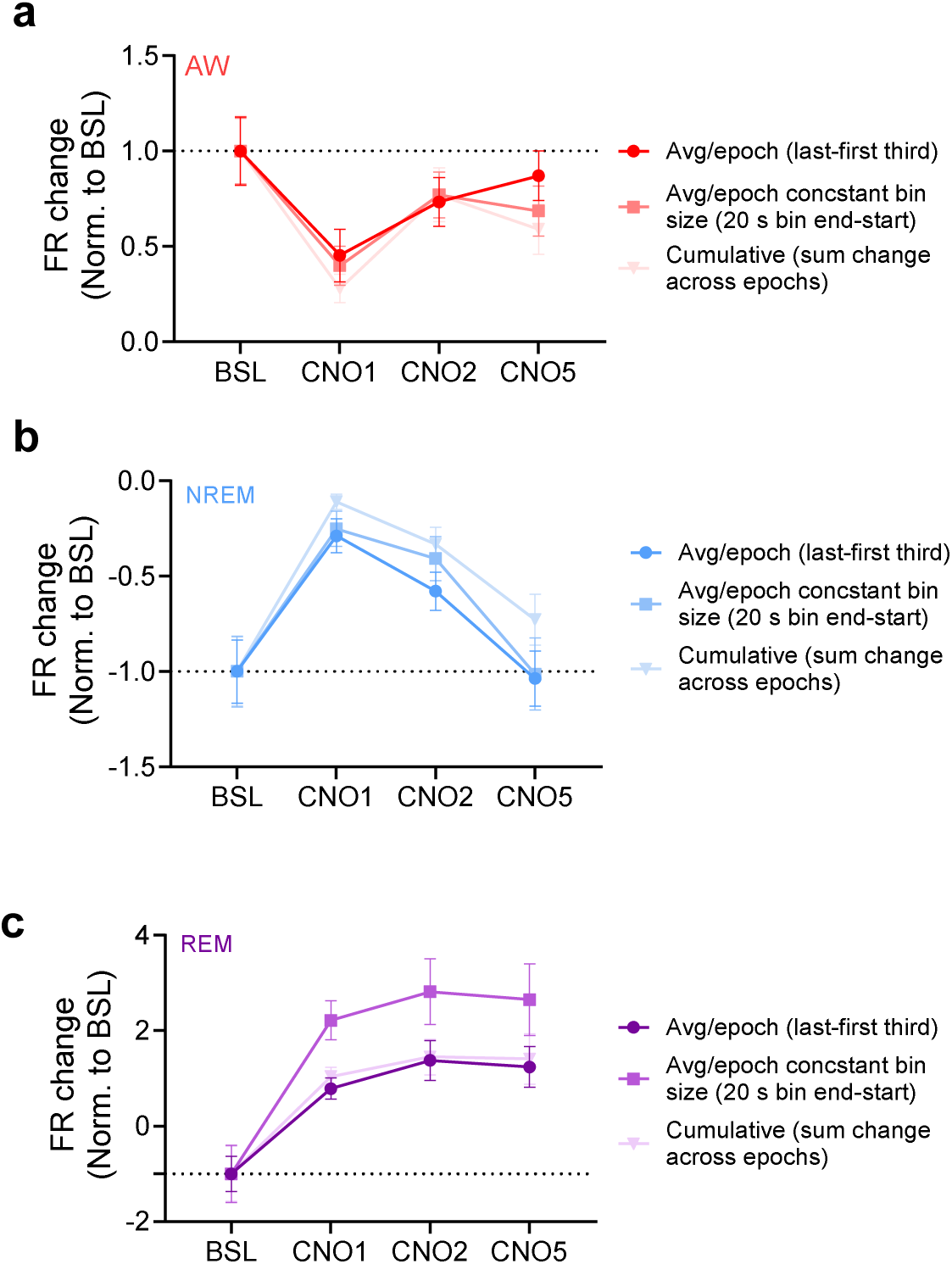
Robustness of state-dependent FR changes across analysis parameters (related to Fig. 5). **a-c,** Normalized FR changes are shown for AW (a), NREM (b), and REM (c) using three complementary quantification methods: (i) Avg/epoch (thirds)-difference between the FR in the last and first third of each epoch (as in Fig. 5); (ii) Avg/epoch (constant bins)-difference between the end and start of each epoch, where end/start are defined as fixed 20 sec windows; (iii) Cumulative change-sum of end–start differences across all epochs for each neuron. To enable comparison across states, values were normalized to the absolute baseline change (set to 1), while preserving the sign of the change (e.g., negative for NREM/REM at baseline). All methods revealed significant FR suppression in AW on CNO1 that recovered by CNO5 (CNO1: \**P* ≤ 0.01; CNO2: *P* ≥ 0.37; CNO5: *P* ≥ 0.1). NREM likewise showed suppression on CNO1 with renormalization by CNO5 (CNO1: \*\*\**P* ≤ 0.0006; CNO2: *P* ≤ 0.06; CNO5: *P* ≥ 0.33). REM exhibited a consistent sign reversal, shifting from baseline decreases to increases following CNO (CNO1: \**P* ≤ 0.03; CNO2: \*\**P* ≤ 0.002; **CNO5: \*\**P* ≤ 0.004). Two-way ANOVA with Dunnett’s multiple comparison test (a-c).

**Extended Data Fig. 14.**
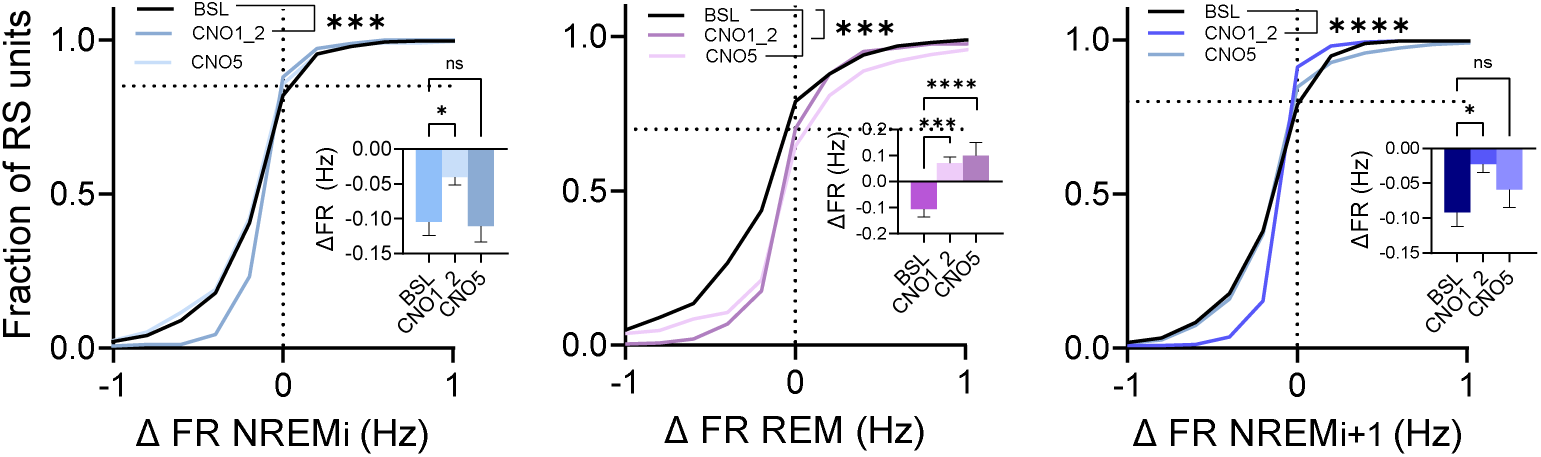
State dependent firing rate changes in NREM-REM-NREM triplets (related to Fig. 5). Cumulative distributions of within-epoch FR changes for NREM_i_, REM, and NREM_i+1_ from NREM_i_–REM–NREM_i+1_ triplets, with insets showing mean changes. NREM_i_ and NREM_i+1_ were suppressed on CNO1_2 (pooled CNO1–2), while REM shifted from decreases at baseline (BSL; pooled BSL1–2) to increases during CNO1_2 and CNO5 (Kolmogorov– Smirnov tests with Bonferroni correction: NREM_i_, BSL vs CNO1_2 \*\*\**P* < 0.001, BSL vs CNO5 *P* = 0.9; REM, BSL vs CNO1_2 \*\*\**P* < 0.001, BSL vs CNO5 \**P* < 0.05; NREM_i+1_, BSL vs CNO1 \*\*\**P* < 0.001, BSL vs CNO5 *P* = 0.9. Insets: NREM_i_ and NREM_i+1_, CNO1_2 \**P* < 0.05, CNO5 *P* > 0.3; REM, CNO1 and CNO5 \*\*\**P* < 0.001; linear mixed-effects model with random intercepts for mouse, comparing each day to BSL, followed by Bonferroni correction). For pairwise statistical contrasts, see Table S4e.

**Extended Data Fig. 15.**
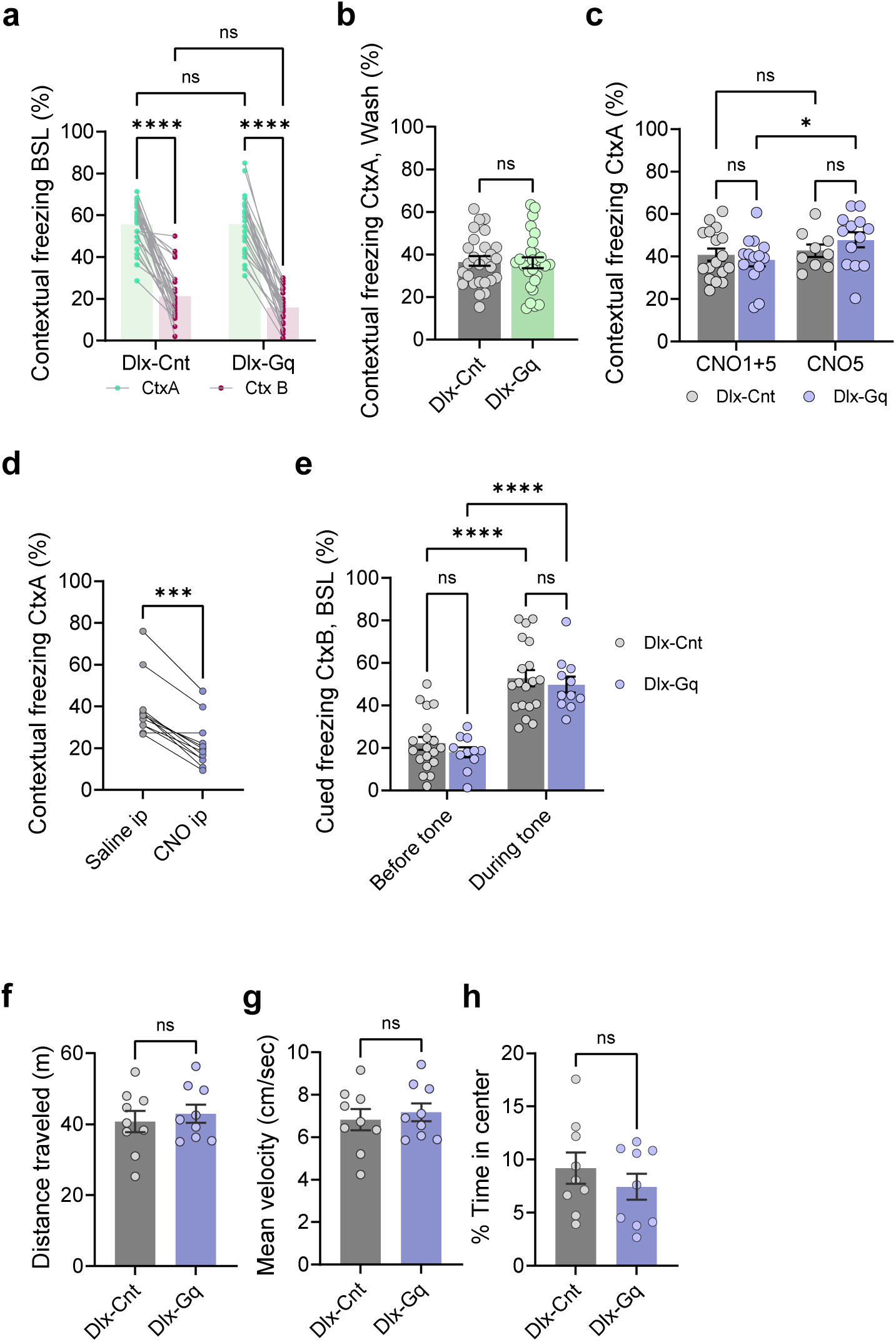
Behavioral effects of chronic hM3Dq activation in INs (related to Fig. 7). **a,** Contextual freezing levels during retrieval 0 (BSL) in CtxA and CtxB for the Dlx-Cnt (n = 27) and Dlx-Gq (*n* = 27) groups. (Dlx-Cnt ctxA vs. Ctxb \*\*\*\**P* < 0.0001; Dlx-Gq CtxA vs. CtxB \*\*\*\**P* < 0.0001; CtxA Dlx-Cnt vs. Dlx-Gq *P* = 0.97; CtxB Dlx-Cnt vs. Dlx-Gq *P* = 0.08). **b**, Contextual freezing levels during retrieval 3 (washout) for Dlx-Cnt (*n* = 27) and Dlx-Gq (*n* = 27) groups. No significant difference was observed (*P* = 0.86). **c**, Contextual freezing levels for two groups: Group I mice underwent both retrieval 1 and 2 (at CNO1+5; Dlx-Cnt: *n* = 17, Dlx-Gq: *n* = 14), and Group II mice underwent only retrieval 2 (at CNO5; Dlx-Cnt: *n* = 9, dlx-Gq: *n* = 13). Dlx-Gq mice that underwent both retrieval 1 and 2 showed reduced freezing compared to those that underwent only retrieval 2 (CNO1+5 Dlx-Cnt vs. Dlx-Gq *P* = 0.56; CNO5 Dlx-Cnt vs. Dlx-Gq *P* = 0.31; Dlx-Cnt CNO1+5 vs. CNO5 *P* = 0.68; Dlx-Gq CNO1+5 vs. CNO5 \**P* = 0.038). **d**, Contextual freezing levels in Dlx-Gq mice (*n* = 12) after intraperitoneal saline or CNO injection one week after the first retrieval. CNO caused a significant reduction in freezing, indicating impaired memory retrieval (\*\**\*P* = 0.001). **e**, Cued freezing levels at retrieval 0 (BSL) before and during the tone. Both Dlx-Cnt (*n* = 19) and Dlx-Gq (*n* = 11) groups exhibited increased freezing during the tone (Dlx-Cnt: \*\*\*\**P* < 0.0001, Dlx-Gq: \*\*\*\**P* < 0.0001). No significant group differences were found (before tone: *P* = 0.40; during tone: *P* = 0.55). **f-h**, Open field test parameters for Dlx-Cnt and Dlx-Gq groups (*n* = 9 per group) measured on CNO Day 1, showing no significant differences in distance traveled (f), velocity (g), or time spent in the center (h) (all comparison *P* > 0.38). Two-way ANOVA with Fisher’s LSD (a,c,e), Mann-Whitney test (b, f-h), Wilcoxon matched-pairs signed rank test (d).

### Supplementary tables

**Supplamental Table 1.**
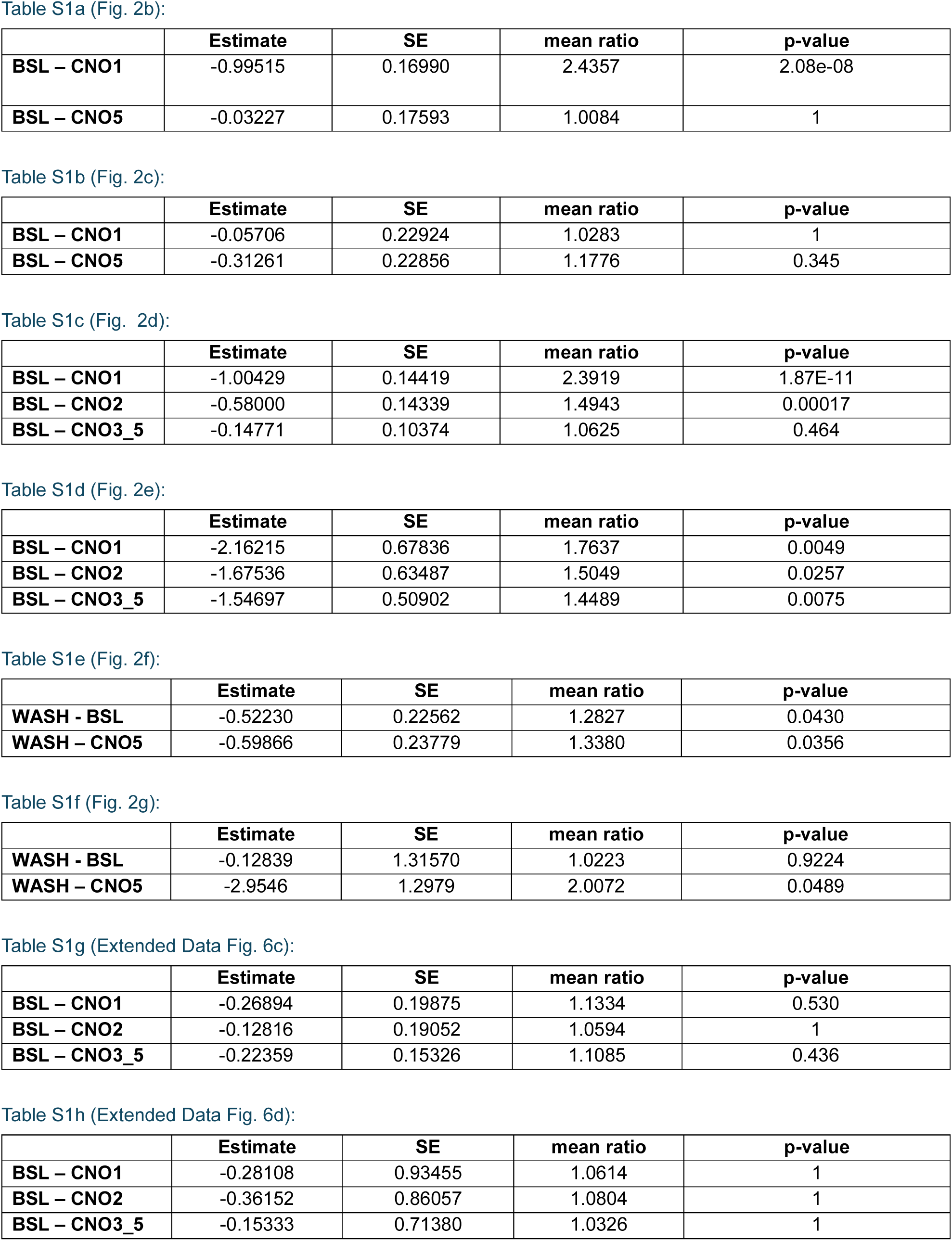
Statistical comparisons of MFR by experimental day, related to Figure 2 and Extended Data Fig. 6. Data were analyzed using a linear mixed-effects model (fr ∼ experimental day + (1|mouse_id)), with mouse identity included as a random effect to account for repeated measures and inter-animal variability. Tables present contrasts between experimental conditions, reporting the estimate (predicted difference), its standard error (SE), the mean ratio (estimate indicating fold-change), and adjusted p-values (adjusted for multiple comparisons using Bonferroni correction). BSL refers to –48 to −0 h, CNO1 to 12–24 h, CNO2 to 24–48 h,CNO3_5 to 60 to 108, CNO5 to 96 to 108 and WASH to 144–156 h.

| Table S1a (Fig. 2b): |  |  |  |  |
|  | Estimate | SE | mean ratio | p-value |
|---|---|---|---|---|
| **BSL – CNO1** | -0.99515 | 0.16990 | 2.4357 | 2.08e-08 |
| **BSL – CNO5** | -0.03227 | 0.17593 | 1.0084 | 1 |
| Table S1b (Fig. 2c): |  |  |  |  |
|  | Estimate | SE | mean ratio | p-value |
| **BSL – CNO1** | -0.05706 | 0.22924 | 1.0283 | 1 |
| **BSL – CNO5** | -0.31261 | 0.22856 | 1.1776 | 0.345 |
| Table S1c (Fig. 2d): |  |  |  |  |
|  | Estimate | SE | mean ratio | p-value |
| **BSL – CNO1** | -1.00429 | 0.14419 | 2.3919 | 1.87E-11 |
| **BSL – CNO2** | -0.58000 | 0.14339 | 1.4943 | 0.00017 |
| **BSL – CNO3_5** | -0.14771 | 0.10374 | 1.0625 | 0.464 |
| Table S1d (Fig. 2e): |  |  |  |  |
|  | Estimate | SE | mean ratio | p-value |
| **BSL – CNO1** | -2.16215 | 0.67836 | 1.7637 | 0.0049 |
| **BSL – CNO2** | -1.67536 | 0.63487 | 1.5049 | 0.0257 |
| **BSL – CNO3_5** | -1.54697 | 0.50902 | 1.4489 | 0.0075 |
| Table S1e (Fig. 2f): |  |  |  |  |
|  | Estimate | SE | mean ratio | p-value |
| **WASH - BSL** | -0.52230 | 0.22562 | 1.2827 | 0.0430 |
| **WASH – CNO5** | -0.59866 | 0.23779 | 1.3380 | 0.0356 |
| Table S1f (Fig. 2g): |  |  |  |  |
|  | Estimate | SE | mean ratio | p-value |
| **WASH - BSL** | -0.12839 | 1.31570 | 1.0223 | 0.9224 |
| **WASH – CNO5** | -2.9546 | 1.2979 | 2.0072 | 0.0489 |
| Table S1g (Extended Data Fig. 6c): |  |  |  |  |
|  | Estimate | SE | mean ratio | p-value |
| **BSL – CNO1** | -0.26894 | 0.19875 | 1.1334 | 0.530 |
| **BSL – CNO2** | -0.12816 | 0.19052 | 1.0594 | 1 |
| **BSL – CNO3_5** | -0.22359 | 0.15326 | 1.1085 | 0.436 |
| Table S1h (Extended Data Fig. 6d): |  |  |  |  |
|  | Estimate | SE | mean ratio | p-value |
| **BSL – CNO1** | -0.28108 | 0.93455 | 1.0614 | 1 |
| **BSL – CNO2** | -0.36152 | 0.86057 | 1.0804 | 1 |
| **BSL – CNO3_5** | -0.15333 | 0.71380 | 1.0326 | 1 |

**Supplamental Table 2.** Statistical comparisons of network properties (mCC, dimentionality, variance and stability index α) by experimental day, related to Fig. 3 and Extended Data Fig. 7-8. Data were analyzed using a linear mixed-effects model (mCC/dimensionality/variance/α ∼ experimental day + (1|mouse_id)), with mouse identity included as a random effect to account for repeated measures and inter-animal variability. Tables present contrasts between experimental conditions, reporting the estimate (predicted difference), its standard error (SE), the mean ratio (estimate indicating fold-change), and adjusted p-values (adjusted for multiple comparisons using Bonferroni correction). BSL refers to –48 to 0 h, CNO1 to 6–24 h, CNO2 to 24–48 h,CNO3 to 48 to 72, CNO4 to 72 to 96 and CNO5 to 96 −108 hr.

|  | **Estimate** | **SE** | **mean ratio** | **p-value** |
|---|---|---|---|---|
| **BSL - CNO1** | 0.01586 | 0.00119 | 0.6921 | 1.487E-38 |
| **BSL - CNO2** | 0.01097 | 0.00089 | 0.7542 | 3.277E-33 |
| **BSL - CNO3** | 0.00800 | 0.00089 | 0.8072 | 2.149E-18 |
| **BSL - CNO4** | 0.00092 | 0.00089 | 0.9678 | 1 |
| **BSL - CNO5** | -0.00152 | 0.00099 | 1.0278 | 0.617 |
|  | **Estimate** | **SE** | **mean ratio** | **p-value** |
| **BSL - CNO1** | -0.06455 | 0.00422 | 1.1763 | 4.383E-50 |
| **BSL - CNO2** | -0.00298 | 0.00316 | 1.0075 | 1 |
| **BSL - CNO3** | 0.01056 | 0.00315 | 0.9760 | 0.004 |
| **BSL - CNO4** | 0.02680 | 0.00317 | 0.9391 | 1.975E-16 |
| **BSL - CNO5** | 0.00694 | 0.00349 | 0.9535 | 0.235 |
|  | **Estimate** | **SE** | **mean ratio** | **p-value** |
| **BSL - CNO1** | -0.30287 | 0.01768 | 1.6186 | 4.20E-62 |
| **BSL - CNO2** | -0.17795 | 0.01323 | 1.2998 | 2.242E-39 |
| **BSL - CNO3** | -0.08194 | 0.01319 | 1.1210 | 2.987E-09 |
| **BSL - CNO4** | -0.01619 | 0.01325 | 1.0199 | 1 |
| **BSL - CNO5** | 0.03382 | 0.01462 | 0.9150 | 0.104 |
|  | **Estimate** | **SE** | **mean ratio** | **p-value** |
| **BSL - CNO1** | -0.15836 | 0.02537 | 1.0824 | 2.472E-09 |
| **BSL - CNO2** | -0.08975 | 0.01899 | 1.0482 | 1.193E-05 |
| **BSL - CNO3** | -0.01615 | 0.01893 | 1.0098 | 1 |
| **BSL - CNO4** | 0.05164 | 0.01901 | 0.9741 | 0.033 |
| **BSL - CNO5** | 0.01456 | 0.02098 | 0.9702 | 1 |
|  | **Estimate** | **SE** | **mean ratio** | **p-value** |
| **BSL - CNO1** | 0.01228 | 0.00772 | 1.0012 | 0.559 |
| **BSL - CNO2** | -0.01005 | 0.00578 | 0.9990 | 0.410 |
| **BSL - CNO3** | -0.00661 | 0.00576 | 0.9993 | 1 |
| **BSL - CNO4** | -0.00763 | 0.00578 | 0.9992 | 0.936 |
| **BSL - CNO5** | 0.00129 | 0.00630 | 1.0001 | 1 |
|  | **Estimate** | **SE** | **mean ratio** | **p-value** |
| **BSL - CNO** | 0.22277 | 0.04351 | 0.9145 | 1.623e-06 |
|  | **Estimate** | **SE** | **mean ratio** | **p-value** |
| **BSL - CNO** | 0.40224 | 0.05021 | 0.8471 | 6.533e-13 |

**Supplamental Table 3.**
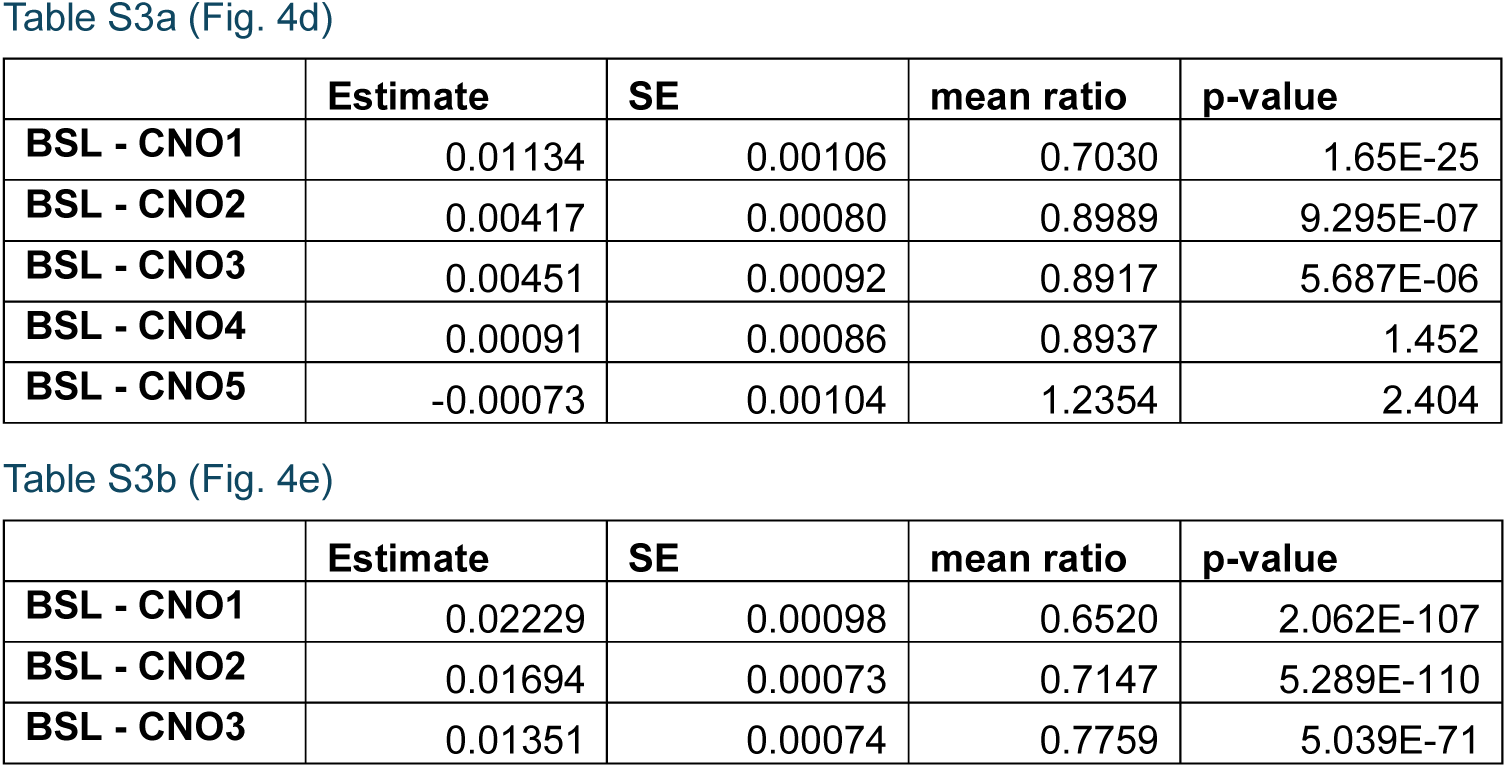
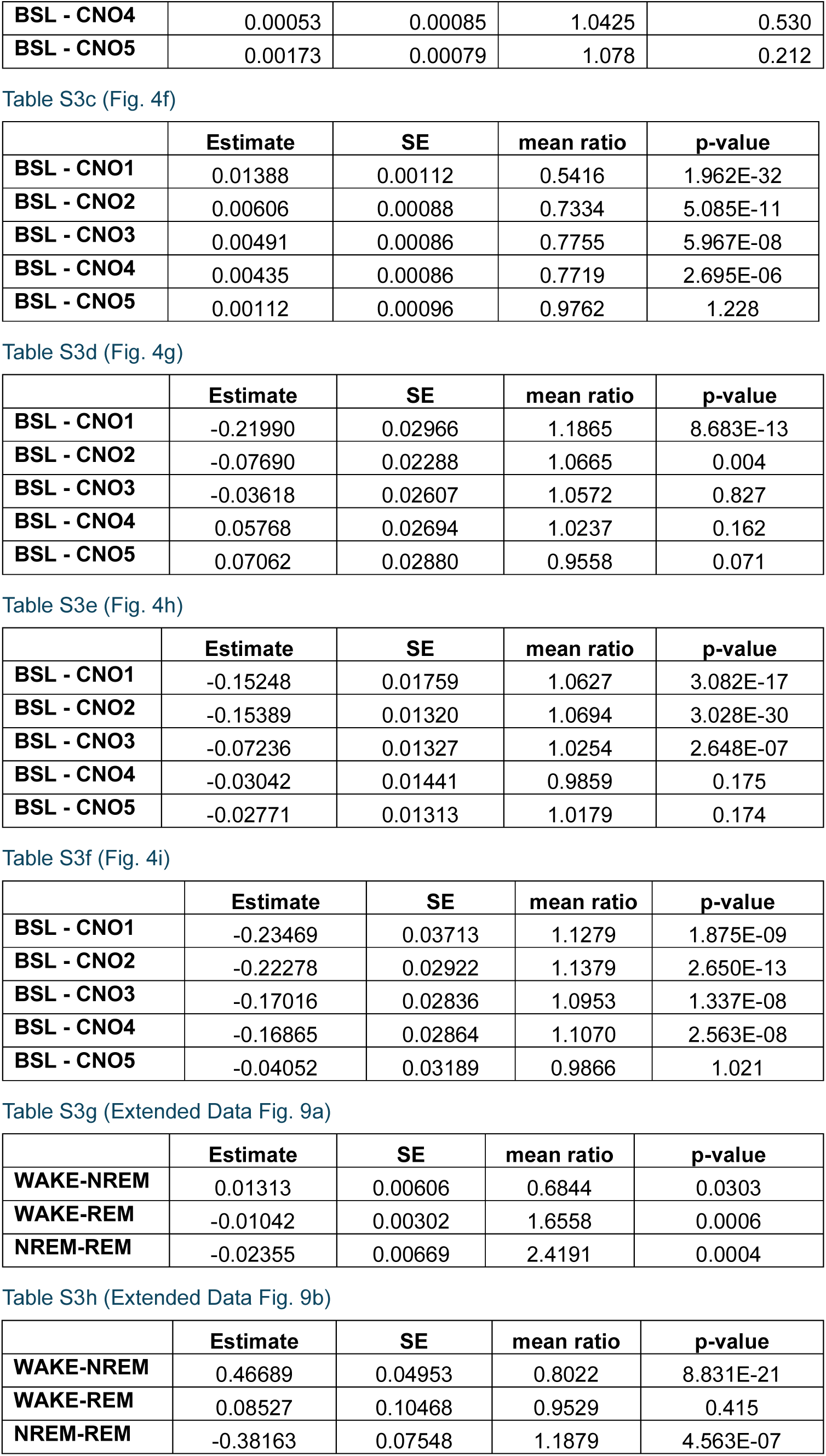
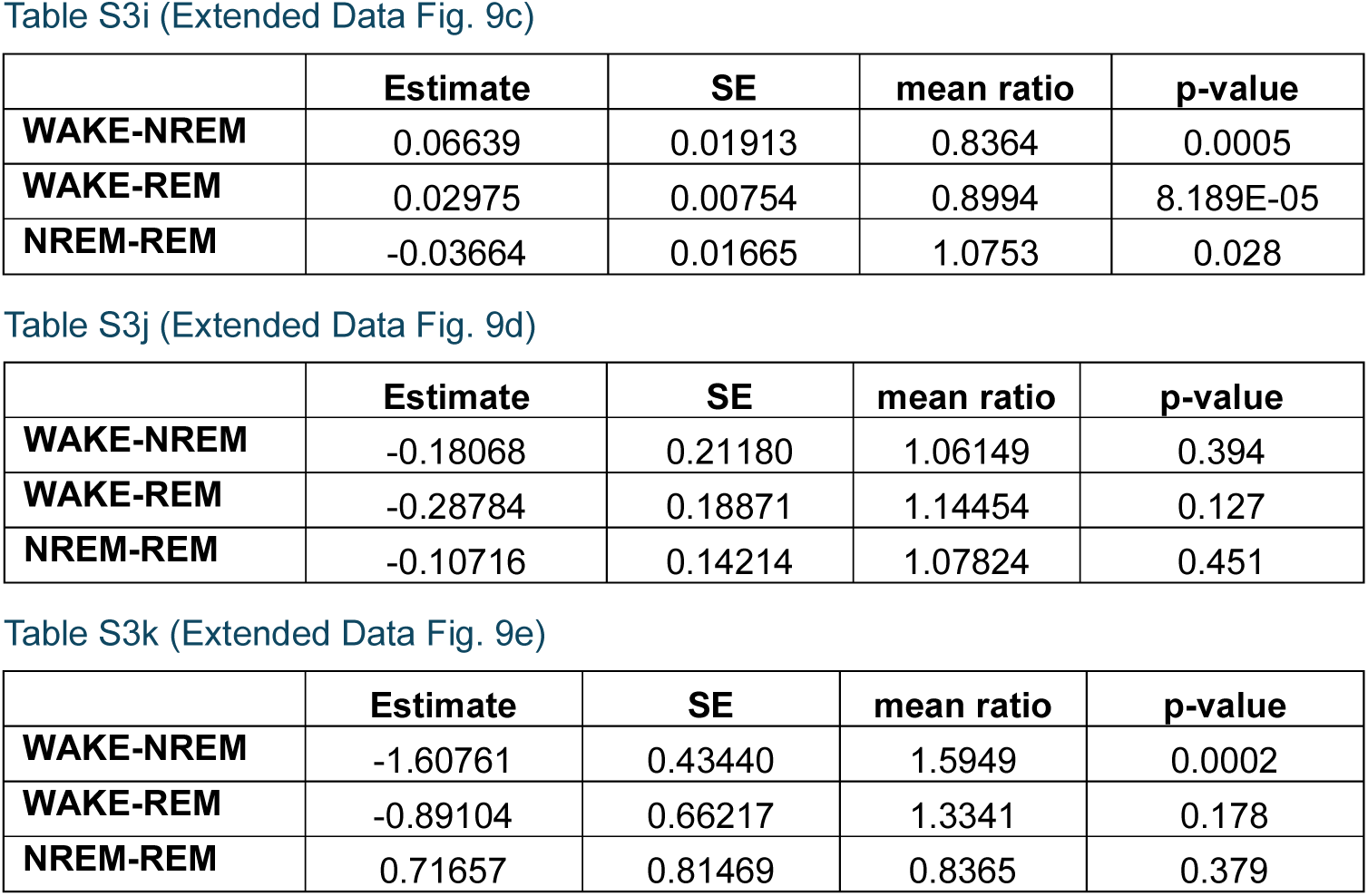
Statistical comparisons of mCC and stability index α by experimental day, and between brain states, related to Fig. 4 and Extended Data Fig. 9. Data were analyzed using linear mixed-effects models. For Figures 4d-i and Extended Data Fig. 9 the model was: network parameter (mCC/dimensionality/variance/α/MFR) ∼ brain state + (1|mouse_id); Mouse was included as random effects to account for repeated measures and inter-animal variability. Tables present contrasts between experimental conditions, reporting the estimate (predicted difference), its standard error (SE), the mean ratio (estimate representing fold-change), and Bonferroni-adjusted *p*-values. Time windows were defined as follows: BSL, –48 to 0 h prior to CNO; CNO1, 6–24 h post-CNO; CNO2, 24–48 h post-CNO; CNO3 to 48 to 72 post-CNO, CNO4 to 72 to 96 and CNO5 to 96 −108 hr post-CNO.

| Table S3a (Fig. 4d) |  |  |  |  |
|---|---|---|---|---|
|  | **Estimate** | **SE** | **mean ratio** | **p-value** |
| **BSL - CNO1** | 0.01134 | 0.00106 | 0.7030 | 1.65E-25 |
| **BSL - CNO2** | 0.00417 | 0.00080 | 0.8989 | 9.295E-07 |
| **BSL - CNO3** | 0.00451 | 0.00092 | 0.8917 | 5.687E-06 |
| **BSL - CNO4** | 0.00091 | 0.00086 | 0.8937 | 1.452 |
| **BSL - CNO5** | -0.00073 | 0.00104 | 1.2354 | 2.404 |
| Table S3b (Fig. 4e) |  |  |  |  |
|  | **Estimate** | **SE** | **mean ratio** | **p-value** |
| **BSL - CNO1** | 0.02229 | 0.00098 | 0.6520 | 2.062E-107 |
| **BSL - CNO2** | 0.01694 | 0.00073 | 0.7147 | 5.289E-110 |
| **BSL - CNO3** | 0.01351 | 0.00074 | 0.7759 | 5.039E-71 |
| **BSL - CNO4** | 0.00053 | 0.00085 | 1.0425 | 0.530 |
| **BSL - CNO5** | 0.00173 | 0.00079 | 1.078 | 0.212 |
| Table S3c (Fig. 4f) |  |  |  |  |
|  | **Estimate** | **SE** | **mean ratio** | **p-value** |
| **BSL - CNO1** | 0.01388 | 0.00112 | 0.5416 | 1.962E-32 |
| **BSL - CNO2** | 0.00606 | 0.00088 | 0.7334 | 5.085E-11 |
| **BSL - CNO3** | 0.00491 | 0.00086 | 0.7755 | 5.967E-08 |
| **BSL - CNO4** | 0.00435 | 0.00086 | 0.7719 | 2.695E-06 |
| **BSL - CNO5** | 0.00112 | 0.00096 | 0.9762 | 1.228 |
| Table S3d (Fig. 4g) |  |  |  |  |
|  | **Estimate** | **SE** | **mean ratio** | **p-value** |
| **BSL - CNO1** | -0.21990 | 0.02966 | 1.1865 | 8.683E-13 |
| **BSL - CNO2** | -0.07690 | 0.02288 | 1.0665 | 0.004 |
| **BSL - CNO3** | -0.03618 | 0.02607 | 1.0572 | 0.827 |
| **BSL - CNO4** | 0.05768 | 0.02694 | 1.0237 | 0.162 |
| **BSL - CNO5** | 0.07062 | 0.02880 | 0.9558 | 0.071 |
| Table S3e (Fig. 4h) |  |  |  |  |
|  | **Estimate** | **SE** | **mean ratio** | **p-value** |
| **BSL - CNO1** | -0.15248 | 0.01759 | 1.0627 | 3.082E-17 |
| **BSL - CNO2** | -0.15389 | 0.01320 | 1.0694 | 3.028E-30 |
| **BSL - CNO3** | -0.07236 | 0.01327 | 1.0254 | 2.648E-07 |
| **BSL - CNO4** | -0.03042 | 0.01441 | 0.9859 | 0.175 |
| **BSL - CNO5** | -0.02771 | 0.01313 | 1.0179 | 0.174 |
| Table S3f (Fig. 4i) |  |  |  |  |
|  | **Estimate** | **SE** | **mean ratio** | **p-value** |
| **BSL - CNO1** | -0.23469 | 0.03713 | 1.1279 | 1.875E-09 |
| **BSL - CNO2** | -0.22278 | 0.02922 | 1.1379 | 2.650E-13 |
| **BSL - CNO3** | -0.17016 | 0.02836 | 1.0953 | 1.337E-08 |
| **BSL - CNO4** | -0.16865 | 0.02864 | 1.1070 | 2.563E-08 |
| **BSL - CNO5** | -0.04052 | 0.03189 | 0.9866 | 1.021 |
| Table S3g (Extended Data Fig. 9a) |  |  |  |  |
|  | **Estimate** | **SE** | **mean ratio** | **p-value** |
| **WAKE-NREM** | 0.01313 | 0.00606 | 0.6844 | 0.0303 |
| **WAKE-REM** | -0.01042 | 0.00302 | 1.6558 | 0.0006 |
| **NREM-REM** | -0.02355 | 0.00669 | 2.4191 | 0.0004 |
| Table S3h (Extended Data Fig. 9b) |  |  |  |  |
|  | **Estimate** | **SE** | **mean ratio** | **p-value** |
| **WAKE-NREM** | 0.46689 | 0.04953 | 0.8022 | 8.831E-21 |
| **WAKE-REM** | 0.08527 | 0.10468 | 0.9529 | 0.415 |
| **NREM-REM** | -0.38163 | 0.07548 | 1.1879 | 4.563E-07 |
| Table S3i (Extended Data Fig. 9c) |  |  |  |  |
|  | **Estimate** | **SE** | **mean ratio** | **p-value** |
| **WAKE-NREM** | 0.06639 | 0.01913 | 0.8364 | 0.0005 |
| **WAKE-REM** | 0.02975 | 0.00754 | 0.8994 | 8.189E-05 |
| **NREM-REM** | -0.03664 | 0.01665 | 1.0753 | 0.028 |
| Table S3j (Extended Data Fig. 9d) |  |  |  |  |
|  | **Estimate** | **SE** | **mean ratio** | **p-value** |
| **WAKE-NREM** | -0.18068 | 0.21180 | 1.06149 | 0.394 |
| **WAKE-REM** | -0.28784 | 0.18871 | 1.14454 | 0.127 |
| **NREM-REM** | -0.10716 | 0.14214 | 1.07824 | 0.451 |
| Table S3k (Extended Data Fig. 9e) |  |  |  |  |
|  | **Estimate** | **SE** | **mean ratio** | **p-value** |
| **WAKE-NREM** | -1.60761 | 0.43440 | 1.5949 | 0.0002 |
| **WAKE-REM** | -0.89104 | 0.66217 | 1.3341 | 0.178 |
| **NREM-REM** | 0.71657 | 0.81469 | 0.8365 | 0.379 |

**Supplamental Table 4.** Statistical comparisons of single neuron FR by experimental day within brain states, related to Fig. 5 and Extended Data Fig. 14. Data were analyzed using linear mixed-effects models. For Figures 5c,j-l, the model was *fr ∼ time_point + (1|mouse_id) + (1|neuron_id),* to account for repeated measures within mice and within neurons. For Fig. 3d-f,m-n, Extended Data Fig. 14a the model was *Δfr ∼ day + (1|mouse_id)*; and Mouse and neuron identity were included as random effects to account for repeated measures and inter-animal variability. Tables present contrasts between experimental conditions, reporting the estimate (predicted difference), its standard error (SE), the mean ratio (estimate representing fold-change), and Bonferroni-adjusted *p*-values. Time windows were defined as follows: BSL1, –48 to –24 h prior to CNO; BSL2, –24 to 0 h prior to CNO; CNO1, 6–24 h post-CNO; CNO2, 24–48 h post-CNO; CNO5, 96–108 h post-CNO; BSL is the combination of BSL1 and BSL2, and CNO1_2, the combination of CNO1 and CNO2.

|  | **Estimate** | **SE** | **mean ratio** | **p-value** |
|---|---|---|---|---|
| **AW first – second third** | 0.05179 | 0.00957 | 0.9709 | 8.210e-08 |
| **AW first – last third** | 0.08949 | 0.00957 | 0.9507 | 1.487e-19 |
| **NREMS first – second third** | -0.07137 | 0.00945 | 1.0443 | 1.118e-13 |
| **NREMS first – last third** | -0.09045 | 0.00945 | 1.0569 | 2.235e-20 |
| **REMS first – second third** | -0.00571 | 0.01649 | 1.0039 | 0.73 |
| **REMS first – last third** | -0.06695 | 0.01649 | 1.0475 | 0.00011 |
| Table S4b (Fig. 5d,e,f) |  |  |  |  |
|  | **Estimate** | **SE** | **mean ratio** | **p-value** |
| **AW last third BSL1 - BSL2** | 0.00736 | 0.01861 | 0.9543 | 0.693 |
| **AW last third BSL1 - CNO1** | -0.05598 | 0.01934 | 2.8393 | 0.016 |
| **AW last third BSL1 - CNO2** | -0.02348 | 0.01942 | 1.3644 | 0.45 |
| **AW last third BSL1 - CNO5** | -0.01081 | 0.01960 | 1.1539 | 0.69 |
| **NREMS last third BSL1 - BSL2** | 0.00723 | 0.02079 | 1.3152 | 0.73 |
| **NREMS last third BSL1 - CNO1** | 0.06241 | 0.02155 | 3.4572 | 0.015 |
| **NREMS last third BSL1 - CNO2** | 0.02718 | 0.02169 | 1.7254 | 0.34 |
| **NREMS last third BSL1 - CNO5** | -0.02499 | 0.02189 | 0.9644 | 0.34 |
| **REMS last third BSL1 - BSL2** | -0.02127 | 0.04130 | 1.4451 | 0.607 |
| **REMS last third BSL1 - CNO1** | 0.11391 | 0.04283 | -1.2661 | 0.011 |
| **REMS last third BSL1 - CNO2** | 0.14483 | 0.04326 | -0.7250 | 0.003 |
| **REMS last third BSL1 - CNO5** | 0.13132 | 0.04348 | -0.8055 | 0.005 |
| Table S4c (Fig. 5j-l) |  |  |  |  |
|  | **Estimate** | **SE** | **mean ratio** | **p-value** |
| **BSL NREM<sub>i</sub> first - last third** | -0.10474 | 0.01935 | 1.0615 | 1.872e-07 |
| **BSL REM first - last third** | -0.10505 | 0.03032 | 1.0736 | 0.0006 |
| **BSL NREM<sub>i+1</sub> first - last third** | -0.09178 | 0.0202 | 1.0546 | 8.771e-06 |
| **BSL NREM<sub>i</sub> first - NREM<sub>i+1</sub> last third** | -0.12823 | 0.01843 | 1.0763 | 4.128e-11 |
| **CNO1_2 NREM<sub>i</sub> first - last third** | -0.04032 | 0.01138 | 1.065 | 0.0009 |
| **CNO1_2 REM first - last third** | 0.07114 | 0.02330 | 0.9123 | 0.0032 |
| **CNO1_2 NREM<sub>i+1</sub> first - last third** | -0.02196 | 0.01263 | 1.0358 | 0.083 |
| **CNO1_2 NREM<sub>i</sub> first - NREM<sub>i+1</sub> last third** | -0.04773 | 0.01246 | 1.0778 | 0.0006 |
| **CNO5 NREM<sub>i</sub> first - last third** | -0.11053 | 0.02292 | 1.0778 | 8.228e-06 |
| **CNO5 REM first - last third** | 0.09983 | 0.05024 | 0.9307 | 0.0477 |
| **CNO5 NREM<sub>i+1</sub> first - last third** | -0.05887 | 0.02558 | 1.0409 | 0.0292 |
| **CNO5 NREM<sub>i</sub> first - NREM<sub>i+1</sub> last third** | -0.09054 | 0.02816 | 1.0628 | 0.0028 |
| Table S4d (Fig. 5m-n) |  |  |  |  |
| Comparison | **Estimate** | **SE** | **mean ratio** | **p-value** |
| **NREM<sub>i</sub>-> REM BSL-CNO1_2** | 0.29109 | 0.05448 | -1.4371 | 1.239e-07 |
| **REM->NREM<sub>i+1</sub> BSL-CNO1_2** | -0.4859 | 0.07283 | -1.9597 | 5.147e-11 |
| **NREMi-> REM BSL-CNO5** | 0.0922 | 0.05880 | 2.1616 | 0.1170 |
| **REM->NREMi+1 BSL-CNO5** | -0.1940 | 0.07950 | 5.9045 | 0.0149 |
| Table S4e (Extended Data Fig.14a) |  |  |  |  |
| Comparison | **Estimate** | **SE** | **mean ratio** | **p-value** |
| **NREM<sub>i</sub> BSL-CNO1_2** | 0.05867 | 0.02396 | 2.5979 | 0.029 |
| **NREM<sub>i</sub> BSL-CNO5** | -0.02594 | 0.02613 | 0.9476 | 0.32 |
| **REM<sub>i</sub> BSL-CNO1_2** | 0.1762 | 0.04567 | -1.4766 | 0.00012 |
| **REM<sub>i</sub> BSL-CNO5** | 0.20488 | 0.04927 | -1.0523 | 7.212e-05 |
| **NREM<sub>i+1</sub> BSL-CNO1_2** | 0.06406 | 0.0254 | 5.7456 | 0.024 |
| **NREM<sub>i+1</sub> BSL-CNO5** | 0.00988 | 0.02766 | 1.5591 | 0.72 |

### Supplementary methods

#### Viruses

We used sAAV-5/2-hDlx-HBB-chI-hM4D(Gi)_2A_NLS_dTomato (Dlx-hM4Di group, v129, ETH Zurich Viral Vector facility) or the control viral vector sAAV-5/2-hDlx-HBB-chI-HA_NLS_dTomato (Dlx-Cnt group, made by ETH Zurich Viral Vector facility) in Figure S8.

#### Intrahippocampal MK801 administration

Local delivery of MK801 was achieved by subcutaneously implanting an osmotic pump (Alzet #1007D) on the animal’s back^4^. A vinyl catheter tubing (Durect #007760) was connected to the flow moderator of the osmotic pump on one end and to a 21 AWG cannula (5 mm length) on the other end. The 21 AWG cannula was positioned to encompass the 33 AWG cannula that was implanted in the CA1 region. The two cannulas were covered in dental cement to ensure stability and prevent leakage. One day prior to implantation, the pumps, catheter, and cannula were preloaded with 1.25 mM MK801 and incubated in 37℃.

#### Behavioral assays

##### Open Field Test

The open field test was conducted in an open plastic arena (40X40X40 cm). Mice were placed in the corner of the chamber and allowed to freely explore for 10 min. Activity in the center and periphery of the field was measured using an automated video-tracking system (EthoVision). The percentage of time in the center of the arena is defined as the percent of total time that was spent in the central 19x19 cm area (25%) of the open field.

##### Delta-Maze test

Spatial working memory was assessed using a continuous variant of the T-maze, the Delta-maze^5^. Mice traversed the central arm of a delta-shaped maze and alternated between left and right turns during consecutive trials. Correct trials were reinforced with a drop of condensed milk delivered at the end of the selected side arm. To prevent retracing, hinged doors blocked re-entry into the same side arm, forcing mice to return to the base of the central arm via a connecting arm before initiating a new trial. At the base, mice were confined for a specified delay before the central-arm door was reopened to start the next trial. In the first trial of each session, both side arms contained a reward. Prior to training, mice were habituated to the apparatus for 15 minutes a day over 5 consecutive days. During training, mice performed 10 trials of the task with a 10-s delay. During testing, they performed 10 trials at delays of 10, 60, and 90 s. To facilitate learning and assure consistent motivation, food intake was restricted to maintain body weight at 85-90% of ad libitum body weight. Success rate was defined as the percentage of correct alternations relative to the total number of trials. Dlx-Gq mice were first tested without pump implantation and were subsequently implanted with a saline-filled osmotic pump, after which testing was performed 4–7 hr post-implantation.

#### CA1 network model

We consider a network consisting of *N*^E^ excitatory and *N*^I^ inhibitory neurons. Due to the timescales of the recordings (from hours to days), we modeled the activity of each neuron, *i* in population α ∈ {*E*, *I*},*r*^α^, as a rate dynamics:

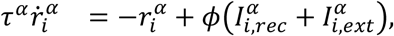

where r_α_ is the neuronal integration time, ф is the transfer function of the neuron and the total current consist of inputs from the recurrent interactions within CA1, 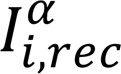, and inputs from outside CA1, external input, 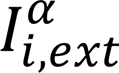. We take the transfer function to be a rectified linear unit (ReLU) activation function.

##### Recurrent interactions

The synaptic inputs from the recurrent interactions within CA1 is given by

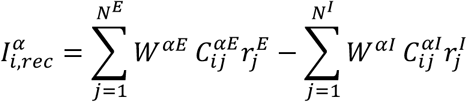

where 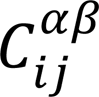 is the adjacency matrix of connections from *β* to α and *W*^αβ^ their synaptic strength. Connectivity is generated from a binominal distribution: each pair is independently connected with probability *p*^β^ (based on the population of the presynaptic neuron), generating on average *K*^E^ = *N*^E^*p*^E^ excitatory and *K*^I^ = *N*^I^*p*^I^ inhibitory inputs. We enforced *K*^E^ = *K*^I^ by setting *p*^I^ = *p*^E^*N*^E^/*N*^I^, and kept *p*^β^ low to ensure sparse connectivity.

##### External Input

The external input to population α consists of two terms:

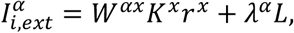

The first term is a constant external input to the network, which in the CA1 network can be thought of excitatory inputs from CA3 via the Schaffer Collateral. Thus, *W*^αx^ is synaptic strength of inputs from the external population, *K*^x^ is average number of external synaptic inputs per neuron, and *r*^x^ is the average firing rate of the external population. For simplicity, the same *W*^αx^ was used for both E and I populations.

The second term was used to model the CNO perturbation. We modeled the CNO perturbation as an additional external input, delivered only to the inhibitory population. Here, *λ*^α^ is the average input current per 1 nM CNO to each population and *L* the CNO concentration (*λ*^E^ was set to 0 to prevent additional input to the excitatory population). Since the osmotic pump is delivering a constant amount of CNO into the body, we assume a constant level of CNO once the perturbation started. Receptor activation was modeled as linear in *L*.

##### Balanced Regime

In large strongly connected random excitatory-inhibitory networks, each neuron receives excitatory and inhibitory inputs that are individually much larger than its firing threshold. In that case, the net current can overwhelm the cells, driving the population firing rate either into quiescence or saturation. Alternatively, inhibitory and excitatory currents can dynamically cancel each other at the leading order, leaving only a residual input of order one^6,7^. In that *balanced state*, this cancellation stabilizes firing rates away from the trivial fixed points and makes neuronal activity sensitive to fluctuations in synaptic input.

To ensure our simulations remained in the balanced regime, parameters were constrained to satisfy

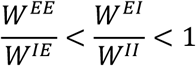

in the absence of CNO, and

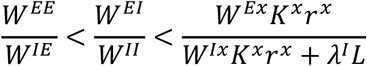

under CNO.

Fitting the CA1 network to data

We fitted network parameters using Simulation-Based Inference (SBI)^8–10^, which performs likelihood-free inference by training neural density estimators on simulated summary statistics.

Given priors over parameters, SBI infers posterior distributions; Here we used maximum a posteriori (MAP) estimates over those distributions to select the specific parameters to use.

Fitting CNO Day 1

We generated 7875 simulations, balanced across outcomes (41.6% paradoxical effect^1,2^, 58.4% not). Data were split into 80% training and 20% testing. A Neural Posterior Estimation model was trained to infer six parameters (*W*^αβ^, *r*^x^, *L*) from nine summary statistics: (1) mean and standard deviation of excitatory and inhibitory firing rates, before and after perturbation; (2) correlation between baseline firing rate and firing-rate change after perturbation,

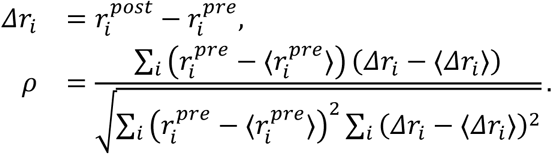

Note that here, for brevity, we dropped the population index. Model accuracy was evaluated by inferring parameters on the held-out test set, rerunning simulations, then comparing to their corresponding test simulations (Figure S5A–C). Any cases outside the balanced regime (see below) or non-physiological firing rates (< 10^−8^ Hz or > 30 Hz) were discarded (8.13%).

Finally, the trained SBI model was applied to experimental recordings from Day 1 of CNO treatment. Firing-rate statistics were computed across all recorded neurons pooled across animals, before osmotic pump implantation and 6 hours after implantation, for 5 hours. The following inferred parameters were used for further simulation (Extended Data Fig. 5h-i), which were compared to experimental data (Extended Data Fig. 5d-g).

##### Fitted and fixed model parameters

| Parameter | Description | Value |
|---|---|---|
| **Fitted Parameters** |  |  |
| *W*<sup>*EE*</sup> | Recurrent excitatory connection | 0.022 mA/Hz |
| *W*<sup>*EI*</sup> | Inhibitory → excitatory | 0.032 mA/Hz |
| *W*<sup>*IE*</sup> | Excitatory → inhibitory | 0.286 mA/Hz |
| *W*<sup>*II*</sup> | Recurrent inhibitory connection | 0.169 mA/Hz |
| *r*<sup>*x*</sup> | External population firing rate | 2.07 Hz |
| *L* | CNO amount/perturbation strength | 0.7 a.u. |
| **Fixed Parameters** |  |  |
| *N*<sup>*E*</sup> | Number of excitatory neurons | 7500 |
| *N*<sup>*I*</sup> | Number of inhibitory neurons | 2500 |
| *p*<sup>*E*</sup> | Excitatory connection probability | 1/15 |
| *p*<sup>*I*</sup> | Inhibitory connection probability | 1/5 |
| *θ* | ReLU threshold | 0 mA |
| *β* | ReLU gain (both populations) | 1 Hz/mA |
| *τ*<sup>*E*</sup> | Excitatory time constant | 1 s |
| *τ*<sup>*I*</sup> | Inhibitory time constant | 0.1 s |
| *K*<sup>*x*</sup> | External inputs per neuron | 500 |
| *W*<sup>*αx*</sup> | External synaptic strength | 0.045 mA/Hz |
| *λ*<sup>*E*</sup> | CNO current response (E) | 0 mA/nM |
| *λ*<sup>*I*</sup> | CNO current response (I) | 223.607 mA/nM |

